# The DYNAM-O Toolbox: Characterizing Individualized Neural Signatures in Sleep EEG

**DOI:** 10.64898/2026.08.26.747401

**Authors:** Mingjian He, Sophie R. Saremsky, Habiba Noamany, Shuqiang Chen, Michael J. Prerau

**Affiliations:** Department of Anesthesiology, Perioperative and Pain Medicine, Stanford School of Medicine, Palo Alto, CA, USA; Department of Psychology, Stanford University, Palo Alto, CA, USA; Division of Sleep and Circadian Disorders, Brigham and Women’s Hospital, Boston, MA, USA; Department of Medicine, Harvard Medical School, Boston, MA, USA

## Abstract

Here, we introduce the Dynamic Oscillation (DYNAM-O) Toolbox, an open-source, cross-platform (MATLAB, Python, and Rust) software package for data-driven characterization of individualized neural dynamics in sleep EEG. Conventional sleep electroencephalography (EEG) measures often rely on predefined bands, thresholds, and averages that incompletely capture transient oscillatory dynamics across the night. For example, sleep spindles, 12–16 Hz bursts of EEG activity related to memory consolidation and altered in disease states, are traditionally detected using fixed frequency and amplitude criteria. Recent studies, however, have shown that canonical spindles represent only a small subset of the tens of thousands of transient “spindle-like” oscillations occurring each night, comprising multiple event classes across a broad frequency range. Together, these classes of transient oscillations form highly individualized signatures of brain state with demonstrated utility as EEG biomarkers of disease. DYNAM-O substantially extends our transient oscillation framework and provides the first end-to-end pipeline for this analysis, from event detection to group-level statistics. It identifies transient oscillations as time-frequency peaks (TF-peaks) using a novel multi-resolution procedure that combines time- and frequency-optimized multitaper spectrograms to separate closely spaced events and refines each event’s peak frequency to resolve oscillations separated by less than the spectrogram resolution. It then computes intrinsic and sleep-state-dependent extrinsic features for each event and represents the overnight distributions of TF-peaks as feature histograms spanning oscillation frequency, slow oscillation power, and slow oscillation phase, preserving continuous brain-state variation obscured by stage-based averaging. New parametric and spline-based dimensionality reduction methods distill these histograms into a small number of interpretable modes, and integrated whole-histogram statistical tests support exploratory and hypothesis-driven group analyses. As a demonstration, we analyzed sex differences in polysomnography recordings from 112 adults (C3 electrode; 67 females, 45 males; ages 20–35 years) from the Cleveland Family Study. In addition to replicating the established higher fast-spindle frequency in females, we identified greater low-alpha transient oscillatory activity in females, a novel sex difference in events ignored in traditional analyses. DYNAM-O provides an accessible and interpretable framework for studying individualized sleep physiology, with applications to longitudinal studies, group comparisons, and feature extraction for deep learning models.

## Introduction

Since the onset of sleep research, electroencephalography (EEG) has been a key modality for the study of sleep physiology [1–6]. For decades, clinicians and researchers have carefully read EEG time traces (i.e., brain waves) to understand the dynamically changing brain states occurring during sleep [7–9]. Visual inspection of sleep EEG recordings has defined both macroscopic, architectural features of sleep, such as distinct sleep stages, and waveform-level features of sleep, such as k-complexes and sleep spindles, which form the cornerstone of standardized rules to review and interpret polysomnograms (PSG) in clinical practice and research [10]. However, PSG brain wave patterns remain noisy and difficult to quantify by human observers [7,11,12], often producing diverging results from repeated recordings or simply from two different raters interpreting the same recording [13–16]. To this end, automated signal processing tools and computerized sleep EEG measures drastically transformed PSG research [17] and generated critical insights that are challenging to obtain with visual inspection [2]. Studies have revealed numerous features of sleep that are stable and highly individualized, including aspects of the EEG power spectrum [18,19], waveform morphological traits [20], properties of sleep spindles [2,18,21,22], as well as beyond EEG signals, such as with temporal patterns of respiratory events [23].

A fundamental challenge in using common sleep EEG measures to capture neural dynamics is that most conventional measures were defined heuristically instead of from a principled basis [24–26]. A prominent example is sleep spindles, which are traditionally defined as a train of distinct waves oscillating within ∼11-16 Hz and lasting more than 0.5 seconds [10]. This definition stemmed from the earliest days of visual inspection of PSG [27] and motivated subsequent automatic detection methods to implement thresholds in order to mimic human scoring [9,28,29]. Automated EEG analysis algorithms are now ubiquitous in sleep research [7,8,14], including numerous sleep spindle detectors [30–32]. The majority of these algorithms utilize some forms of signal processing transforms, such as filtering spindle frequency bands, taking the squared voltage, and then applying pre-determined thresholding to isolate out signals of interest [33]. By trying to replicate human experts, most automated algorithms apply cutoff thresholds and inherit restrictions that stem from conventional practices in visual inspection of sleep EEG time traces [30]. Our earlier work has shown that sleep spindles visually detected by human experts only represent about 30% of spindle-like transient oscillations in the spindle frequency range during non-rapid eye-movement (NREM) sleep [24]. We later expanded this time-frequency analysis to all spindle-like transient oscillations beyond the spindle frequency range. By studying all transient oscillations on spectrograms, we coined the term “time-frequency peak” (TF-peak) to denote any electrophysiological event in the EEG that is spindle-like in its waveform morphology [34]. These TF-peak patterns are highly robust across multiple nights of sleep from the same subjects. Furthermore, TF-peaks allowed one to summarize the dynamics of tens of thousands of transient oscillations in a single image, which revealed previously unreported changes in several patient groups compared to controls [35]. These results highlight the importance of applying novel approaches to capture neural dynamics during sleep, in ways less restricted by prior assumptions and conventions rooted in visual inspections.

Despite many technological advances and the growing recognition that conventional EEG measures only capture a limited fraction of sleep neural dynamics [11,36–39], the field has been slow in moving away from clinical heuristics and towards data-driven methods [40,41]. Perhaps the greatest motivating factor of change in sleep analysis has been the release of accessible software toolboxes. As observed with a few prominent tools in the past: YASA for automatic sleep staging [42], LUNA for analyzing large volumes of sleep EEG data [43], and FOOOF for separating periodic oscillations and aperiodic activity [44], improved analytic methods are adopted rapidly and become increasingly utilized in sleep EEG studies once the software solutions are made readily available to researchers.

Here, we introduce the Dynamic Oscillation (DYNAM-O) Toolbox to provide an open-source, cross-platform, computational toolbox for characterizing individualized brain states during sleep in terms of the dynamics of tens of thousands of transient oscillations in EEG. DYNAM-O provides an interpretable, intuitive, and visually rich representation of a broad class of spindle-like waveform events at multiple time scales. Its outputs can be used to perform statistical tests and derive measures to both describe patterns at an individual level and discover changes at a population level. More broadly, DYNAM-O offers a powerful window into the collection of sleep features that are highly individualized and show general night-to-night stability. By providing novel tools to characterize neural dynamics during sleep more accurately, we aim to enhance the understanding of population variations and to establish a stronger connection between sleep and health at an individual level. We hope this toolbox distribution can help reveal crucial insights that would otherwise be overlooked due to limitations of conventional sleep EEG measures.

In this paper, we provide a general overview of the code, methods, and science behind the toolbox. We will describe key methodological steps involved in extracting transient oscillations and provide best practices for analysis and visualization of results. We also demonstrate the robustness of feature histograms as representations of neural dynamics during sleep in DYNAM-O, along with algorithms to parameterize the feature histograms and to compute group statistics all within the toolbox. With this comprehensive introduction of the DYNAM-O Toolbox, we aim to facilitate a broader adoption of data-driven approaches for studying neural dynamics during sleep and for identifying novel, interpretable, and reproducible electrophysiological biomarkers.

## Materials and Methods

The primary goal of the Dynamic Oscillation (DYNAM-O) Toolbox is to efficiently implement and expand upon our previously established approach of characterizing multidimensional dynamics of different classes of spindle-like transient oscillations in the sleep EEG spectrogram. This framework includes methods to extract time-frequency peaks and their properties, to visualize distributions of TF-peak features in an interpretable and efficient manner, and to conduct statistical tests to gain insights into sleep physiology. In doing so, we aim to provide a powerful tool through which researchers can explore stable features of sleep EEG, efficiently capturing the neural dynamics of tens of thousands of transient oscillatory events throughout a night.

To achieve these analytic goals, the processing pipeline of the DYNAM-O Toolbox is structured into four main parts:

1. Identifying transient oscillation events as TF-peaks from spectrograms.
2. Computing feature properties that describe each TF-peak.
3. Characterizing overnight dynamics of all TF-peaks with a distributional approach.
4. Conducting statistical tests of TF-peak dynamics on the new distributional characterization.

A graphical overview of the DYNAM-O pipeline is shown in Figure 1, and the analyses of TF-peaks can be applied autonomously to any single-channel electrophysiological recordings from overnight sleep EEG data. In the rest of the Methods sections, we describe each of the four parts in more details. A step-by-step flowchart of different functional modules underlying the DYNAM-O pipeline is also provided in Appendix 1, along with further implementation details describing computational steps involved in the processing pipeline.

**Figure 1.**
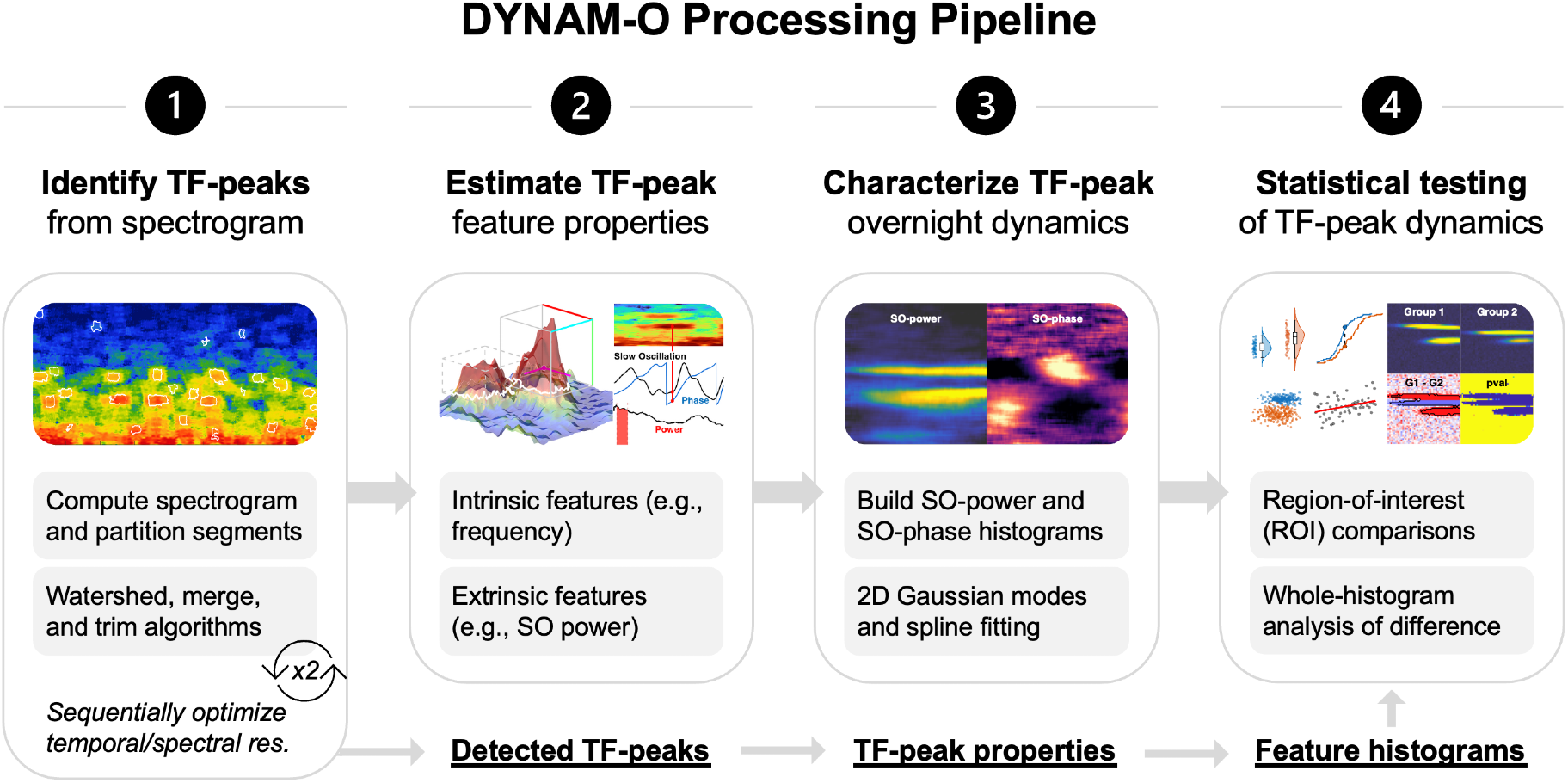
Schematic diagram of the DYNAM-O processing pipeline. The DYNAM-O pipeline is organized into four sequential parts, each feeding its output into the next. **Part 1.** Transient oscillation events are identified as TF-peaks by computing a multitaper spectrogram, partitioning it with the watershed algorithm, and then merging and trimming the resulting regions, with two spectrograms processed sequentially to optimize temporal and spectral resolutions. **Part 2.** Intrinsic feature properties (e.g., peak frequency, bandwidth, duration) and extrinsic feature properties (e.g., SO power and SO phase) are computed to describe each detected TF-peak. **Part 3.** The overnight dynamics of all TF-peaks are characterized using SO-power and SO-phase feature histograms. Optional dimensionality reduction can be performed on the histograms to represent them using a small number of two-dimensional Gaussian modes or spline coefficients. **Part 4.** Region-of-interest and whole-histogram analyses are performed on the feature histograms to detect group differences in transient oscillation dynamics. TF-peak = Time-frequency peak; SO = Slow oscillation.

### Code availability

When using the DYNAM-O Toolbox in published work, users should cite both the present paper and our earlier study that established the transient oscillation framework underlying the toolbox [34]; referencing these two papers is sufficient to acknowledge the DYNAM-O methods and software. An open-source distribution of the DYNAM-O Toolbox (version v3.1) is available at https://github.com/preraulab/DYNAM-O_toolbox.

The DYNAM-O Toolbox is available in MATLAB, Python, and Rust. DYNAM-O is developed in pure MATLAB, which remains the reference implementation and provides the complete visualization, dimensionality reduction, and statistical testing functionality described in detail in the following sections. In order to create multi-fold increases in performance, the computationally heavy components of DYNAM-O have been optimized and ported to Rust, which is used as a backend by Python wrapper, as well as the high-performance backend of the MATLAB library via MEX. While extra care was given to maximizing parity between all three versions, unavoidable differences in low-level libraries remain. Despite these differences, comparison runs on the same data for all versions are typically within a fraction of a percentage of each other for peak counts and remain virtually identical at the distributional level.

The pure MATLAB implementation of DYNAM-O also incorporates several external open-source toolboxes, which we acknowledge here. Time-frequency decomposition of sleep EEG data is performed using the Prerau Lab multitaper spectrogram [38,45]. Whole-histogram group comparisons control the false discovery rate using the Benjamini-Hochberg/Benjamini-Yekutieli procedures implemented in the mass univariate analysis code base [46]. Finally, some figures make use of the *colorcet* perceptually uniform and optimized rainbow colormaps [47].

### Data availability

Example data of a single-channel EEG recording of overnight sleep and its scored sleep stages used throughout this paper is obtained from a previous study [48] with original authors’ permission. This example data is distributed together with the DYNAM-O Toolbox and can be freely downloaded. Data from the Cleveland Family Study [49] presented in the group comparison section is available as part of the National Sleep Research Resource (NSRR) [50] database with free access upon request and compliance with NSRR’s data usage agreement.

### Part 1: Identifying TF-peaks from Spectrograms

#### What are TF-peaks?

To study the time-frequency information contained in sleep EEG, we first take a time-frequency transform of the recording time trace into a spectrogram (Figure 2a) to reflect electrophysiological activity at different frequencies and time points. By definition, a transient oscillation at a narrow-banded frequency will manifest as a confined region with elevated activity on a spectrogram with sufficient resolutions [24,34], i.e., a topographical peak in the time-frequency domain. Three of the regions showing increased activity likely correspond to canonical sleep spindles that are also visible on the time trace (Figure 2a). But the spectrogram reveals many additional regions of transient oscillatory activity at different frequencies. Based on this first principle, we detect transient oscillation events from electrophysiological recordings as *time-frequency peaks* (TF-peaks), which are three-dimensional peaks on the landscape of a spectrogram (Figure 2b). When this analysis is applied to overnight sleep EEG, there is a clear abundance of TF-peaks occurring throughout the night with structures over sleep cycles and dynamics at different frequencies that await further investigation (Figure 2c). It is the goal of the DYNAM-O Toolbox to facilitate all of these analyses.

**Figure 2.**
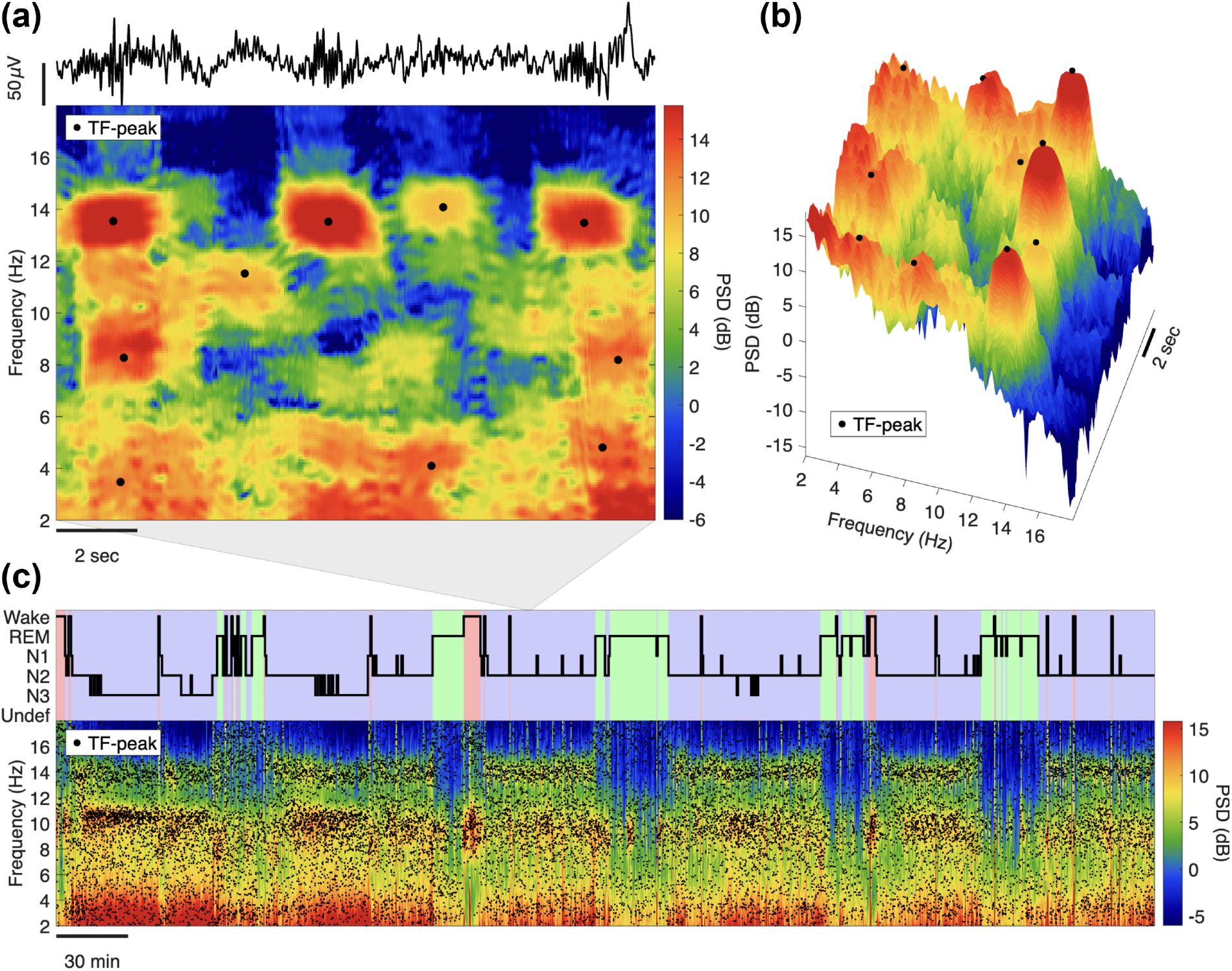
TF-peaks occur throughout a night with structured dynamics. **(a)** A short 15 s multitaper spectrogram (bottom) showing transient oscillatory activity at different frequencies, with warmer colors indicating greater power spectral density (PSD). Each TF-peak region is marked by a black dot at the center of the detected event. While three prominent TF-peaks temporally correspond to canonical sleep spindles that can be observed from the time trace (top), many other TF-peaks are only visible in the time-frequency domain. **(b)** The same 15 s spectrogram in (a) visualized in 3D space to demonstrate the distinctive topographical peaking shape of each detected TF-peak event (black dot) on the spectrogram landscape. While TF-peaks of typical sleep spindles display tall peaks, other TF-peaks also exhibit clear elevated activity within local peak regions. **(c)** Scored hypnogram (top) and the corresponding spectrogram (bottom) from a single EEG channel across an entire overnight recording. The continuous view captures both sleep macrostructure, such as NREM-REM cycling, and microstructure, such as bouts of sigma-band activity during NREM sleep, at a single glance. The spectrogram is overlaid with the tens of thousands of detected TF-peaks (black dots). These discrete transient oscillation events recur with organized structure across frequency and time rather than at random, motivating our analysis of them as individualized markers of sleep neural dynamics.

#### TF-peak detection

The peak detection procedure is summarized in Figure 3 and is comprised, at a high level, of the following steps:

1. For both a time resolution and a frequency resolution optimized spectrogram:
  a. Use the watershed procedure to extract peaks
  b. Merge the watershed regions to identify salient local peaks
  c. Trim the peaks to localize the peak area
2. Combine the peaks boundaries from both resolution spectrograms and extract TF-peaks

**Figure 3.**
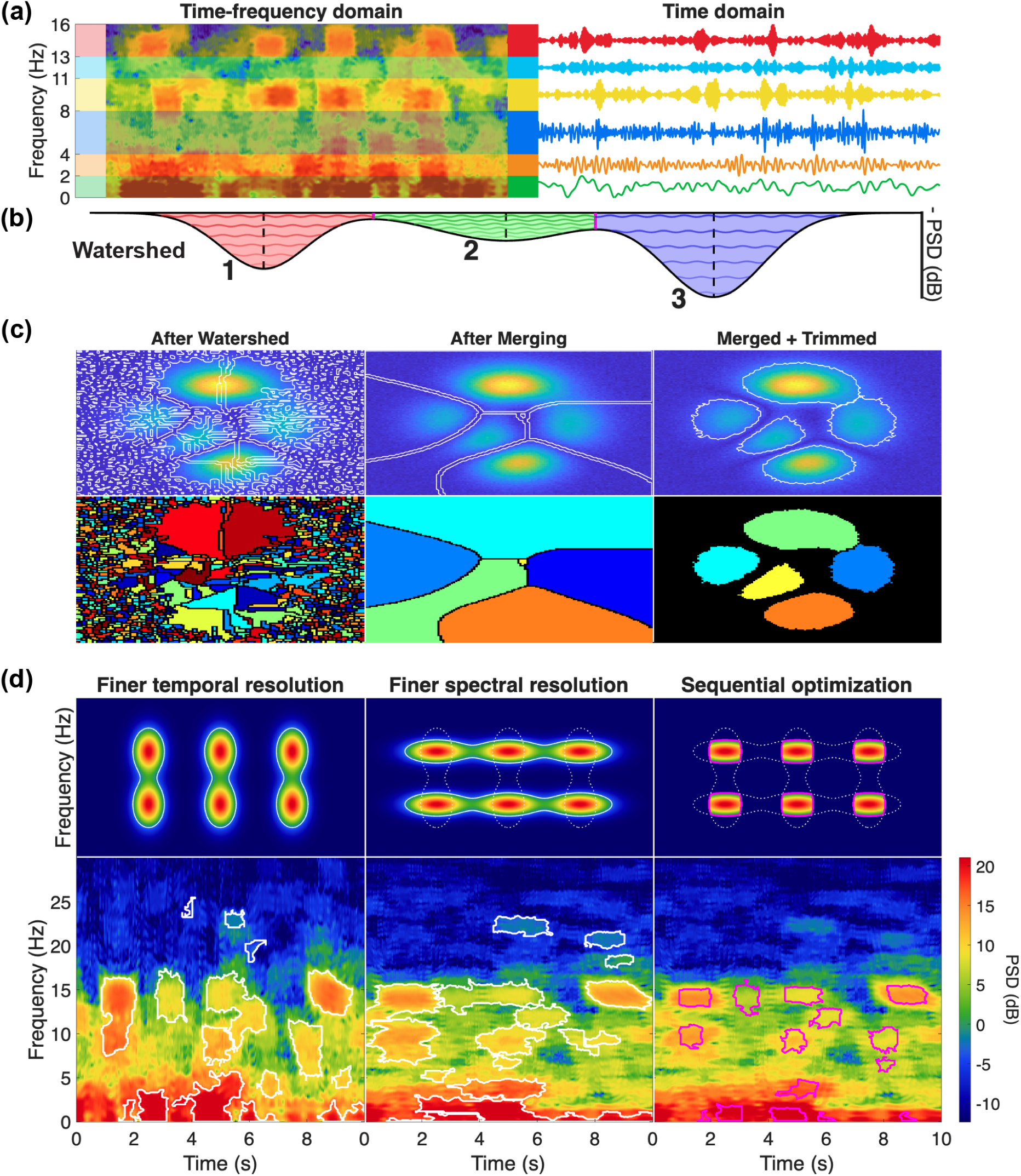
Identifying TF-peaks from spectrograms. **(a)** Correspondence between topographical peaks in the time-frequency domain and transient oscillations in the time domain; arbitrary frequency bands are used only for illustration. Confined regions of elevated PSD on the spectrogram (left) correspond to narrowband transient oscillations shown as bandpass-filtered time traces at illustrative frequencies (right). **(b)** Schematic of the watershed algorithm. Treating the inverted spectrogram as a topographic landscape, the watershed identifies local sinks (numbered 1–3). **(c)** The watershed-merging-trimming procedure, shown as peak contours (top row) and labeled regions (bottom row). A noisy spectrogram is initially over-segmented into many small regions (After Watershed); neighboring regions are iteratively recombined into larger, better-defined peaks (After Merging); and each peak is trimmed to retain 80% of its volume, tightly enclosing its central mass while excluding the extended flat surfaces (Merged + Trimmed). **(d)** Sequential optimization for temporal and spectral resolutions, illustrated with simulated peaks (top) and example sleep EEG spectrograms (bottom). Transient oscillations close in time are resolved under finer temporal resolution with multitaper parameters giving 1 s temporal and 4 Hz spectral resolutions (left), oscillations close in frequency are separated under finer spectral resolution with multitaper parameters giving 2 s temporal and 2 Hz spectral resolutions (middle), and processing the two spectrograms sequentially recovers adjacent events as distinct TF-peaks (right, pink outlines).

We will go over the theory and steps behind this process briefly, with implementation details covered in the Appendices. As explained above and illustrated in Figure 3a, there is a close correspondence between transient oscillations in the time domain and topographical peaks on spectrograms. Therefore, we first compute spectrograms using the multitaper method (MTM) (see Appendix 2 for a comparison with wavelet transform). We use a time-half-bandwidth product of 2 with 3 tapers, along with 1 s or 2 s analysis windows and 0.05 s window step size [38]. To accentuate transient oscillations and distinguish them from aperiodic activity [24], we remove the 1/f-like background baseline activity by normalizing the spectrogram with the 2^nd^ percentile of power spectral density at each frequency across non-artifact periods (Appendix 3).

We expand on our previous approach [34] to identify prominent 3D peaks on spectrogram segments (30s by default) using the watershed algorithm, an image segmentation method widely employed in image processing applications. The watershed algorithm identifies the boundaries between local sinks on a landscape that water would get trapped in when flowing downward (Figure 3b). Leveraging this ability to isolate local minima, we apply the watershed algorithm on the baseline-normalized MTM spectrogram on a linear scale with inverted power spectral density to identify prominent local peaks on the original spectrogram.

While the watershed is highly effective on a perfectly smooth surface, a noisy spectrogram will produce an enormous number of small watershed regions. This is because every tiny local maximum in the topography will trap water and get considered as a distinct peak by the watershed algorithm (Figure 3c). We address this challenge with an improved custom merging algorithm, which iteratively recombines neighboring 3D peaks obtained from the watershed into larger peaks with better defined peak structures (Appendix 4). Since prominent peaks are sparse in the time-frequency space, contours of the merged watershed peaks are too extensive and do not well represent the main body of spectral peaks in the time-frequency domain (Figure 3c). Therefore, we trim each detected peak to retain 80% of its volume, yielding contours that more tightly enclose the central mass of the peak and excluding the extended flat surfaces due to watershed and merging (Figure 3c). We then keep merged and trimmed 3D peaks with properties above detection thresholds as TF-peaks, with the rest discarded as noise. These thresholds are theoretically grounded by MTM parameters and not freely adjusted (Appendix 5). Together, this watershed-merging-trimming processing robustly identifies TF-peak events from a spectrogram.

In the DYNAM-O Toolbox implementation, we employ an additional downsampling of the spectrogram 2D image to accelerate the watershed and merging computations. Once merged watershed regions are obtained to partition the spectrogram into distinct peak regions, the downsampled masks are interpolated using nearest categorical neighbors back to full resolution prior to volume trimming. To perform the downsampling, we decimate the spectrogram (default to a factor of 2) along both time and frequency axes. As the spectrogram is computed with overlapping windows and much larger spectral resolution than frequency steps (default to 0.1 Hz), this downsampling exploits overlapping information in consecutive pixel values to greatly speed up the TF-peak detection process.

#### Sequential optimization for temporal and spectral resolutions

The ability to extract TF-peaks is ultimately limited by the intrinsic properties of the spectral estimator (i.e. the spectrogram) from which they are being derived. In spectral estimation, there is a distinct trade-off between the temporal resolution and frequency resolution in a spectrogram. That is, it is possible to either make a spectrogram that can separate events that are close in time or a spectrogram that can separate events that are close in frequency, but not at the same time. Note that in this case, the term resolution has nothing to do with how many pixels a spectrogram has, but rather the ability to see and differentiate between closely spaced events (see [38] for more details). This issue impacts sleep EEG in particular, since there are complex neural dynamics during sleep, potentially generating distinct electrophysiological events that are close in time and/or in frequency. For example, simultaneous transient oscillations close in frequency may appear as one unified TF-peak with a broad bandwidth. Similarly, if two transient oscillations occur close in time at the same frequency, they may appear as a single TF-peak with a long duration. However, by combining information from different spectral estimators, we can better separate events. To do this, DYNAM-O estimates two different spectrograms: one optimized for time (time-half-bandwidth product of 2 with 3 tapers) with 1 s analysis window, and another for frequency with 2 s analysis window (Figure 3d). By effectively looking at the intersection of peaks boundaries identified by both spectrograms (see Appendix 6 for details), we can separate events that would otherwise be linked in each individual spectrogram in time or frequency. This procedure allows us to better identify transient oscillation events adjacent to each other in the time-frequency domain as distinct TF-peaks, providing improved resolutions for studying sleep neural dynamics through discrete TF-peak events.

#### Part 1 Summary

By the end of this first part, we have extracted tens of thousands of TF-peaks occurring at different frequencies and points in time (Figure 2c). These TF-peaks carry rich information about neural dynamics in sleep EEG as a data-driven decomposition of the time-frequency spectrogram into discrete transient oscillation events. In the next part, we compute feature properties describing each TF-peak to begin quantifying neural dynamics during sleep.

### Part 2: Computing Feature Properties of TF-peaks

For each identified TF-peak, we compute several feature properties (Table 1) that are descriptive statistics to help characterize the 3D peaks for subsequent analyses. The feature properties can be divided into two types: 1) intrinsic properties that describe the geometry and location of each topological peak on the spectrogram (Figure 4a); and 2) extrinsic properties that capture the contextual information about the peak in terms of overnight sleep dynamics.

**Figure 4.**
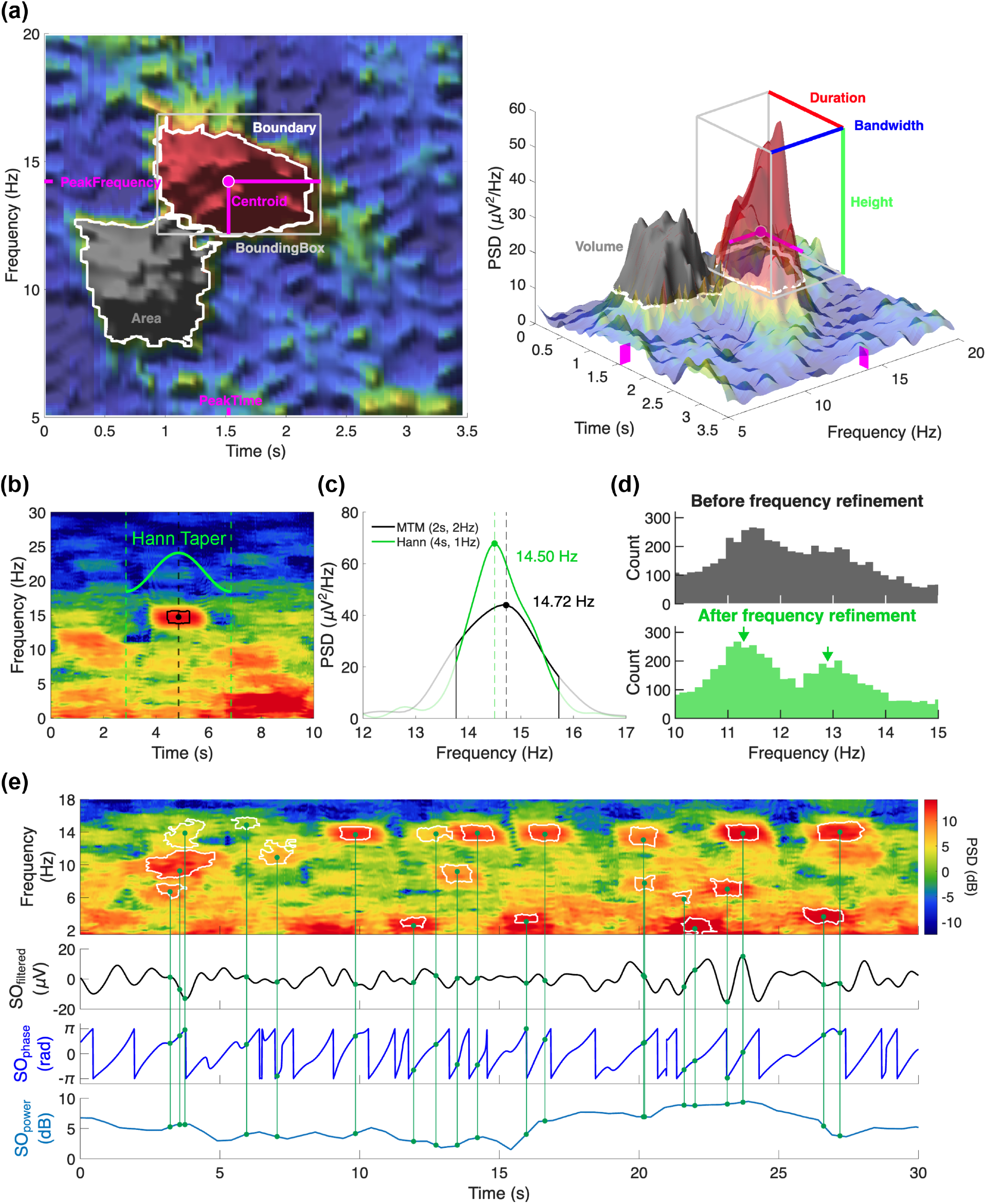
Intrinsic and extrinsic feature properties describing each TF-peak event. **(a)** Intrinsic feature properties illustrated on a single TF-peak, shown as a two-dimensional region on the spectrogram (left) and as a three-dimensional surface (right). **(b)** Peak-frequency refinement uses a 4 s Hann-windowed segment centered at the peak time to obtain a higher-resolution (1 Hz) spectral estimate for each TF-peak. **(c)** Power spectra for an example peak estimated using the detection multitaper spectrogram (2 s window, 2 Hz resolution; black) versus the Hann-window refinement (4 s, 1 Hz resolution; green). **(d)** Distributions of estimated peak frequencies across all TF-peaks before (top) and after (bottom) refinement; refinement better resolves separable clusters of TF-peaks at close frequencies (arrows). **(e)** Extrinsic feature properties place each TF-peak in the context of overnight sleep dynamics. The spectrogram with overlaid TF-peaks (top) is shown together with the slow-oscillation-filtered signal, the continuous SO phase, and the SO power (lower traces); vertical lines link example TF-peaks to the SO phase and SO power values assigned to them at their peak times.

**Table 1.** Feature properties of TF-peaks computed in the DYNAM-O Toolbox.

| Feature | Description | Units |
| --- | --- | --- |
| *Intrinsic* |  |  |
| Area | Time-frequency area of TF-peak | sec*Hz |
| Bandwidth | Peak bandwidth along the frequency axis | Hz |
| Boundaries | (time, frequency) points of peak surrounding boundary | (sec, Hz) |
| BoundingBox | (top-left time, top-left frequency, width, height) box points | (sec, Hz, sec, Hz) |
| Duration | Peak duration along the time axis | sec |
| Height | Peak height above background baseline spectrum | $\mu V^2/Hz$ |
| HeightData | Height of all points in peak region within peak boundary | $\mu V^2/Hz$ |
| PeakFrequency | Peak frequency estimated at peak time with a Hann taper | Hz |
| Peakiness | Sharpness (0-1) of a peak computed as (max-mean)/(max-min) | unitless |
| PeakTime | Peak time based on weighted centroid | sec |
| SegmentNum | Spectrogram segment number | # |
| Volume | Time-frequency volume of TF-peak | sec*$\mu V^2$ |
| *Extrinsic* |  |  |
| PeakStage | Sleep stage at peak time based on input scoring information<br>(6=artifact, 5=Wake, 4=REM, 3=N1, 2=N2, 1=N3, 0=undefined) | Stage # (0-6) |
| SOpower | Slow oscillation (0.3-1.5 Hz) power at peak time | dB |
| SOphase | Slow oscillation (0.3-1.5 Hz) phase at peak time | rad |

#### TF-peak frequency estimation

One of the intrinsic TF-peak features, peak frequency, requires special attention. Neural activity during sleep generates complex electrophysiological signals that are often close in frequency on scalp EEG. This phenomenon is particularly prominent in the alpha and sigma frequency ranges, exemplified by the close frequencies of slow and fast sleep spindles [51]. During the sequential optimization for temporal and spectral resolutions, the second frequency-optimized spectrogram is computed with MTM parameters giving a spectral resolution of 2 Hz. This parameter choice reflects a compromise to avoid smearing spectral peaks by window lengths beyond 2 s during TF-peak detection. However, many electrophysiological patterns induced by transient oscillations during sleep manifest with frequency separations less than 2 Hz [52,53]. Separately, it is difficult to accurately estimate the peak frequency from a MTM spectrum due to the higher bias from spectral leakage in exchange for lower spectral variance. To address these issues, for each identified TF-peak, we refine its peak frequency estimate to 1 Hz spectral resolution: for each identified TF-peak event, we perform a tapered spectral estimation with a Hann window on the 4 s original recording centered at the peak time, which is estimated from the weighted centroid in the watershed-merging-trimming processing (Figure 4b). This procedure updates the initial centroid-based estimate of each event’s peak frequency (Figure 4c) and better reveals separable clusters of TF-peaks at close frequencies (Figure 4d). This refinement step is critical for the next part of the DYNAM-O processing pipeline, where distributions of the frequency property form a key dimension to encode sleep dynamics of TF-peaks. Details of the Hann-windowed frequency refinement are described in Appendix 7.

#### Slow oscillations as a proxy marker of sleep depth, timing mechanisms, and brain state during sleep

We focus on three sleep-state-dependent extrinsic properties: scored sleep stage, slow oscillation (0.3-1.5 Hz) power (SO power), and slow oscillation phase (SO phase). Steps to compute these three properties are described in Appendix 8. These properties are motivated by extensive existing literature on distinct brain states across clinically defined, discretized sleep stages [1,54–56], continuous measures of depth-of-sleep [57–61], and cross-frequency coupling of fast transient oscillations with cortical up/down states reflected by slow frequency oscillations [62–65]. Past research has shown that changes in SO features during sleep are observed in sleep disorders and associated with neuropathology [66,67]. Sleep stages and SO features are therefore critical dimensions to consider when characterizing neural dynamics during sleep. Concretely, TF-peaks observed under different contexts with respect to these extrinsic properties can capture transient oscillations occurring at varying brain states during sleep, even when the peaks have similar frequencies (Figure 4e). Thus, measuring these extrinsic feature properties of TF-peak events provides one way to quantify sleep dynamics viewed through the lens of transient oscillations.

#### Part 2 Summary

We have computed feature properties for each TF-peak event. However, challenges remain as the large numbers of TF-peaks cannot be sufficiently captured by simple averages. The next part of the DYNAM-O processing pipeline addresses this issue by deriving a histogram-based characterization of distributional structures of TF-peaks, visualizing their multi-dimensional feature properties, and providing novel bases for statistical tests on sleep dynamics.

### Part 3: Characterizing TF-peak Dynamics with Feature Histograms

The first two parts of the DYNAM-O pipeline yield a table of identified TF-peaks, each described by a collection of local (intrinsic) and contextual (extrinsic) feature properties. Traditional analyses would at this point select a fixed frequency band for which to define a spindle event and then examine spindle properties as a function of sleep stage and experimental condition. With DYNAM-O, we advocate for a different approach to study sleep dynamics of transient oscillations for several reasons. First, canonical frequency ranges established by clinical sleep scoring guidelines are only generally accurate at a population level. Individuals exhibit substantial variability in their neural oscillation frequencies [18,34,51,68], which are inconsistent with rigid frequency cutoffs [11]. This is especially true in development, aging, and patient populations in which sleep spindles are used as indices of thalamocortical activity [69–71]. Second, brain states during sleep vary continuously and are arbitrarily discretized by sleep stages [8,72]. As a result, there are still substantial variations in neural dynamics within the same sleep stage [38,56,73–75]. Such information will be lost if one simply averages across TF-peaks within a conventionally defined sleep stage. Third, most studies using average spectral power carry an implicit assumption about the importance of neural activity based on strength. Yet, weaker transient oscillations with more consistent occurrence patterns could also represent important neural dynamics. These events get weighted down when averaging measures and are often overlooked in conventional analyses [24]. Thus, DYNAM-O aims to study identified TF-peaks in a more comprehensive and data-driven manner through representing the multi-dimensional feature properties efficiently and facilitating their interpretations. Most importantly, characterizing clusters of repeated occurrences of TF-peaks in the multi-dimensional feature space is a fundamental way to study TF-peaks with minimal assumptions on signal properties including peak strength.

#### Feature histograms

We characterize structures of TF-peak properties with a *distributional* approach, based on the construct of “feature histograms” first introduced in [34]. As the name suggests, feature histograms focus on the occurrence of discrete TF-peak events and display patterns in the numbers of TF-peaks identified across the multi-dimensional feature space. While higher-dimensional histograms can be generated along axes of many combinations of feature properties, here we describe two primary feature histograms featured in DYNAM-O to investigate sleep neural dynamics, namely slow oscillation (SO)-power and SO-phase histograms. As explained in the last section, SO power and SO phase represent two different time scales of extrinsic factors influencing transient oscillations, and both are important for understanding the functional roles and dynamics of TF-peaks observed during a night [58,64].

Both SO-power and SO-phase histograms use the peak frequency estimated during the last part of the pipeline as one of the feature axes. Peak frequencies are particular relevant for studying sleep physiology, given that electrophysiological signals of different frequencies are putatively generated by neural activity with distinct functions during overnight sleep [76–81]. In addition to frequency, sleep research questions are often concerned with varying brain states during sleep [1,7,57]. SO-power histograms use a two-dimensional histogram to encode the density of TF-peak occurrence (events / minute) as a function of peak frequency and SO power, which is an extrinsic feature approximating the depth of overnight sleep. Similarly, SO-phase histograms encode the density of TF-peaks as a function of peak frequency and SO phase, another extrinsic feature describing the contextual cortical up/down state at the peak time of each TF-peak. As SO phase is circular within -π to π, we employ proportional densities by normalizing the densities on a SO-phase histogram, so each row sums to one across a given peak frequency.

Figure 5 illustrates how histogram density values at different overlapping bins are computed from raw counts of TF-peak events, and how SO-power and SO-phase histograms can reveal detailed structures of transient oscillation dynamics, superseding averages within conventional sleep stages. Briefly, the numbers of TF-peaks whose SO features fall inside each SO-power or SO-phase bin along the x-axis are counted separately for each frequency bin along the y-axis. These raw counts are divided by the time spent in the SO-power or SO-phase bin during the overnight recording to produce density measures. This time-in-bin denominator captures the total duration (in minutes) of sleep showing SO features within a particular range, regardless of the occurrence of TF-peaks. Compared to a coarse aggregation of transient oscillation events based on discrete sleep stages, these histograms offer greater resolutions in capturing how often TF-peaks are observed as a function of peak oscillation frequency and SO features. Further details of SO-power/phase histogram computations are described in Appendix 9.

**Figure 5.**
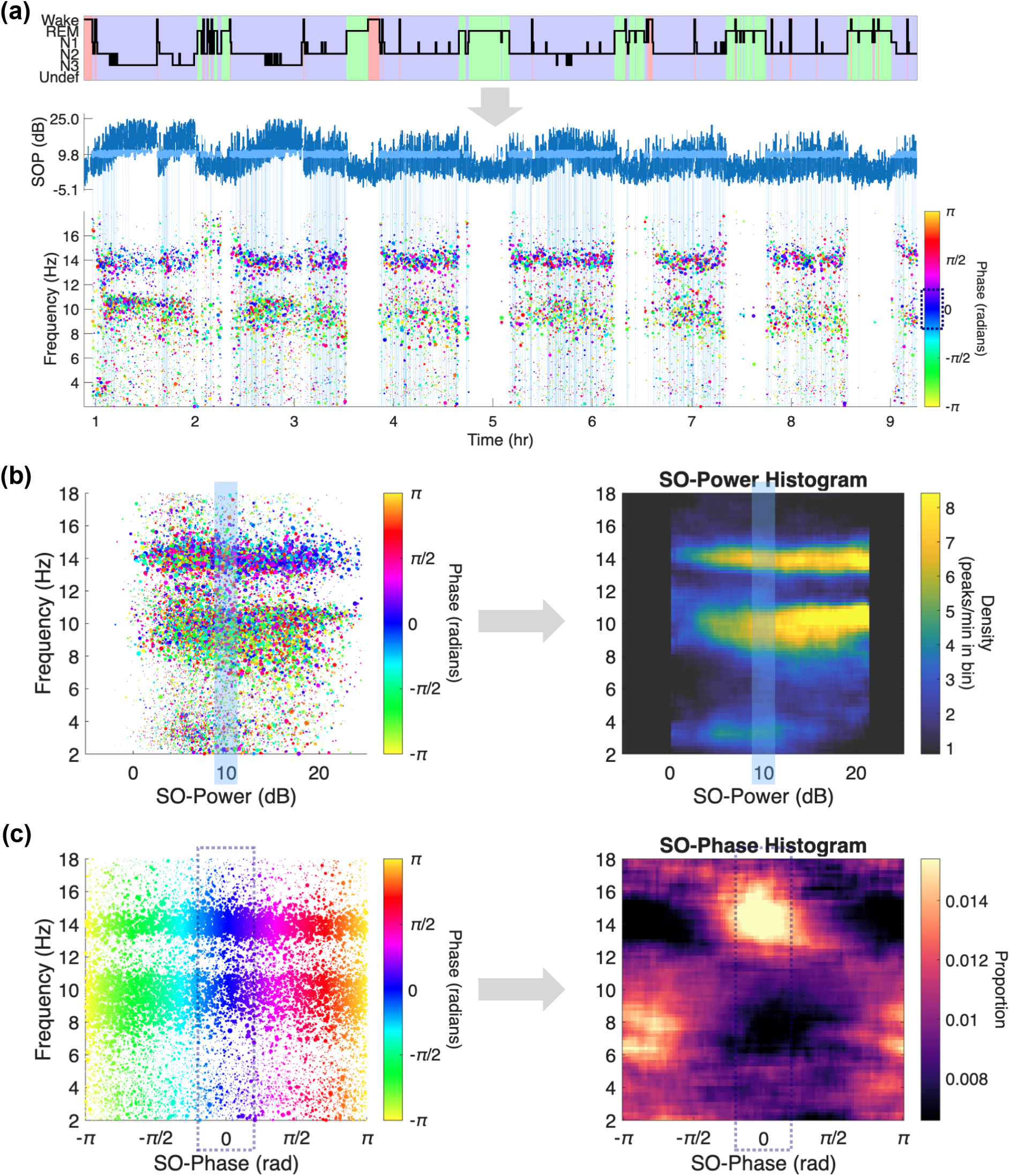
Computation of SO feature histograms from raw counts of TF-peak events. **(a)** An overnight sleep EEG recording can be displayed as a scored hypnogram (top). The sleep stages discretize the underlying brain states based on conventional scoring guidelines. A continuous SO-power (SOP) time series (middle) generalizes the discrete stages to a proxy measure that fluctuates with depth of sleep. A scatter plot of detected TF-peaks (bottom) during NREM sleep shows each TF-peak event colored by the corresponding SO phase at each peak’s center time. TF-peaks from REM and wake periods are omitted in this visualization for illustration clarity. Time periods within an example SOP bin (8.75 to 11.25 dB) are colored in light blue on the SOP time trace and propagate to the scatter plot as vertical bars to indicate which TF-peak events fall within this bin of SOP metric. An example phase bin (-π/5 to π/5 rad) is also marked by a dotted rectangle on the color bar to indicate the range of color for TF-peak events that fall within this bin of SO phase metric. **(b)** Left: scatter points of detected TF-peaks colored by SO phase on a space of SO power by frequency. Right: a 2D histogram showing event densities computed from overlapping bins. The example SOP bin (8.75 to 11.25 dB) is highlighted by vertical light blue bars to indicate the group of TF-peaks that contribute to event counts underlying the histogram, along the bin center column, separately at each frequency. Thus, the SO-power histogram on the right characterizes TF-peak density as a continuous function of peak frequency and SOP**. (c)** Left: scatter points of detected TF-peaks colored by SO phase on a space of SO phase by frequency. Right: a 2D histogram showing proportional event densities computed from overlapping bins. The example phase bin (-π/5 to π/5 rad) is again highlighted by dotted rectangles to indicate the group of TF-peaks that contribute to event counts along the bin center. However, the actual proportional density values on the histogram depend on all TF-peaks occurring at other SO phase values due to the use of a proportional metric. A large proportion indicated by a bright hotspot on the histogram means that TF-peaks at that frequency preferentially occur at that particular phase relative to other phase values. Thus, the SO-phase histogram on the right characterizes the proportional density of TF-peaks as a function of peak frequency and SO phase, normalized across each frequency.

Exemplified by the SO-based histograms, feature histograms enable a rich characterization of feature properties of tens of thousands of TF-peaks occurring at different points in time during sleep. In addition, these histograms provide intuitive visualizations of structures of TF-peak distributions along selected feature axes. This is an important step forward in terms of quickly observing the sleep dynamics across an entire night of sleep at a glance compared to scrolling through 30 s epochs of sleep EEG. However, these feature histograms are still just non-parametric plots of TF-peak activity patterns using a very large number of variables (∼10k bins). Therefore, we would prefer to further reduce their dimensionalities to increase their practical utilities beyond a visualization tool and to facilitate subsequent quantitative studies such as hypothesis testing, clustering, regression analyses, and deep learning.

#### Dimensionality reduction of feature histograms

Going beyond the raw feature histograms we previously employed in [34], here we introduce two different approaches to reduce the dimensionality of SO-power and SO-phase histograms, focusing on preserving prominent patterns of TF-peak distributions (referred to hereafter as “modes”) characterized by these histograms. Analyzing SO feature histograms by modes of densities instead of individual pixel values is a critical way to extend the distributional approach of studying TF-peaks. While each TF-peak is well defined by their local spectral maximum in the time-frequency domain, there is no guarantee that every TF-peak event is necessarily meaningful as a functionally relevant transient oscillation. However, distributional hotspots where TF-peaks tend to occur are much more interpretable, stable, and robust to noise within an individual. Therefore, measuring modes on SO feature histograms makes these dynamics amenable to subsequent analysis.

The first approach fits a parametric model to the feature histograms, decomposing them into Gaussian modes with interpretable parameters. As illustrated using a simulated SO-power histogram in Figure 6a, each of the fitted modes represents a region of increased TF-peak activity, in terms of the number of events occurring per minute, in the feature space. Rather than using the thousands of bins in a histogram, this approach allows us to describe the same data structure with only a few parameters per observed mode. Figure 6b shows that the SO-power histogram obtained from the example data in the DYNAM-O Toolbox can be well represented with only 4 Gaussian modes. Similarly, Figure 6c shows that the SO-phase histogram contains 2 modes. We describe each of these modes and their interpretations later in the **Results** section.

**Figure 6.**
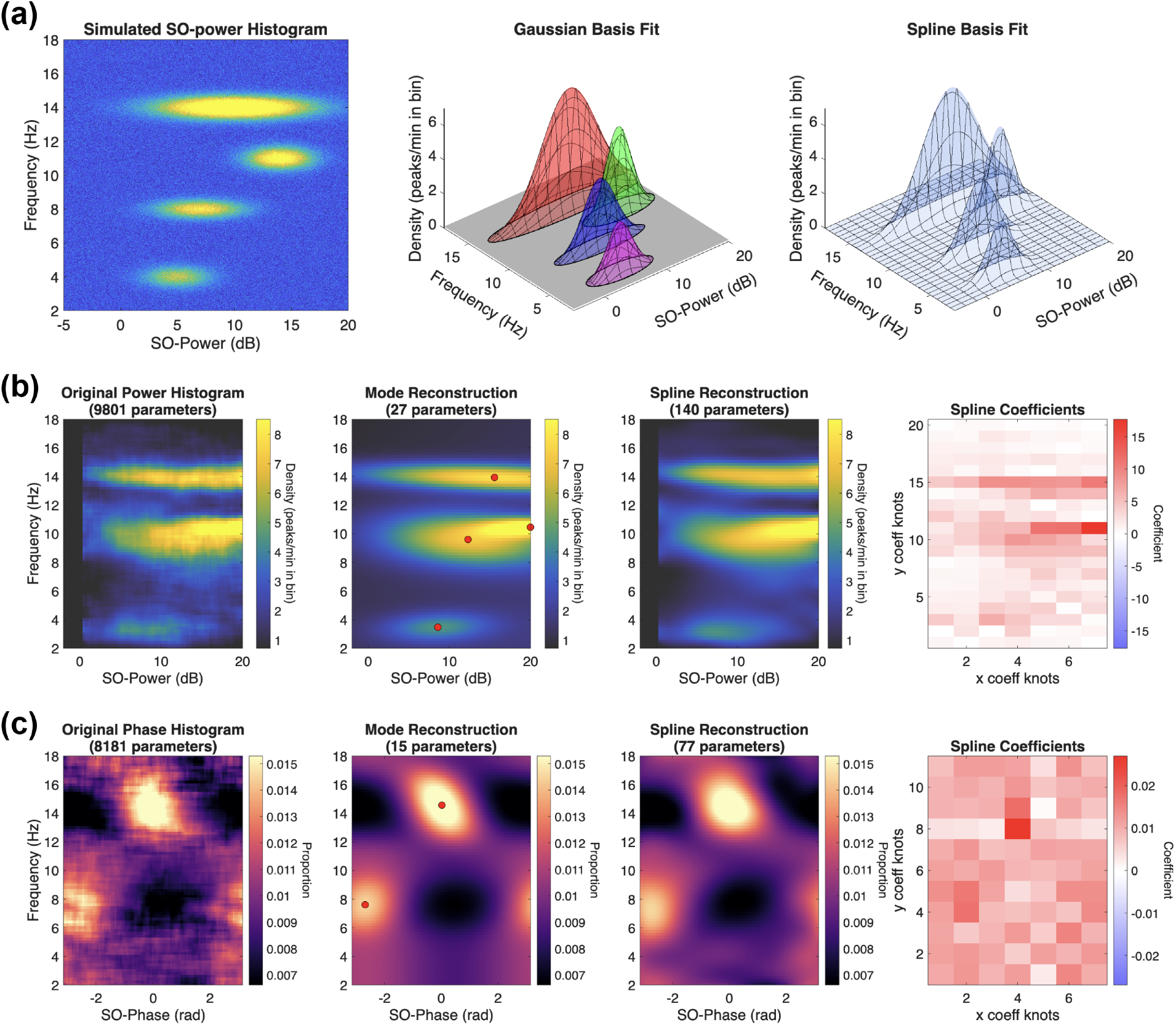
Dimensionality reduction of SOPHs using parametric Gaussian modes and spline fitting. **(a)** A simulated SO-power histogram to illustrate the fitting of histograms using Gaussian or spline basis functions. **(b)** SO-power histogram dimensionality reduction. From left to right: the original histogram, its reconstruction from parametric two-dimensional Gaussian modes, its reconstruction from a two-dimensional spline basis, and the corresponding spline coefficient map. Both reductions recapitulate the major modes of TF-peak density with far fewer parameters. **(c)** SO-phase histogram dimensionality reduction, shown in the same layout: the original histogram, the parametric von Mises-Gaussian mode reconstruction, the spline reconstruction, and the spline coefficients.

Detailed Gaussian basis functions used to model SO-power and SO-phase histograms are described in Appendix 10 along with the parametric model fitting procedure. In addition to fitted Gaussian mode parameters, a number of properties can also be computed to describe each mode. For example, based on the fitted mode center frequency on the SO-power histogram, one can look up the observed extent of phase coupling at that frequency from the SO-phase histogram, which helps describe whether a particular cluster of TF-peak events tends to be phase-locked to a particular SO phase. A full list of automatically computed properties describing each fitted parametric mode is shown below in Table 2 with brief explanation of each property.

**Table 2.** Mode properties of fitted parametric Gaussian modes on SO feature histograms.

| Property | Description | Units |
| --- | --- | --- |
| *Fitted Gaussian mode parameters - available for both histogram types* |  |  |
| Density | Mode height at its center. SO-power: raw Gaussian amplitude. SO-phase: empirical row-normalized full-model height after removing background | power: peaks/min<br>phase: proportion |
| FreqMean | Center frequency of the fitted mode | Hz |
| FreqStd | Gaussian standard deviation of the fitted mode along the frequency axis | Hz |
| Theta | Mode tilt angle as a direct rotation of the SO-power Gaussian or as a frequency-dependent phase shift of the SO-phase mode | rad |
| *Fitted Gaussian mode parameters – SO power only* |  |  |
| SOpowerMean | Center of the fitted mode along the SO-power axis | SO power (dB) |
| SOpowerStd | Gaussian standard deviation along the rotated SO-power-oriented axis | SO power (dB) |
| *Fitted Gaussian mode parameters – SO phase only* |  |  |
| SOphaseMean | Circular phase center of the fitted mode at FreqMean, wrapped to $[-\pi, \pi]$ | rad |
| SOphaseStd | Reciprocal-square-root of concentration of the von Mises term, as a local phase-width scale measure | rad |
| *Derived mode property - available for both histogram types* |  |  |
| Volume | Area under the standalone mode kernel: exact full-plane integral for SO-power; empirical-height area surrogate integral for SO-phase | power: peaks/min*Hz<br>phase: proportion*Hz |
| *Derived phase-coupling properties – SO power only* |  |  |
| PrefPhase | SO-phase bin center where the fitted SO-phase model is maximal at the frequency nearest to FreqMean | rad |
| Coupling | Fitted SO-phase model value at PrefPhase and FreqMean on the SO-phase histogram | proportion |
| *TF-peak summary properties - available for both histogram types* |  |  |
| PkCount | Number of TF-peaks within the 95% confidence density contour of a mode based on Gaussian density | # |
| PkFreq | Mean frequency of assigned TF-peaks | Hz |
| PkDuration | Mean duration of assigned TF-peaks | sec |
| PkBandwidth | Mean bandwidth of assigned TF-peaks | Hz |
| PkHeight | Mean TF-peak height above the fitted background | $\mu V^2/Hz$ |
| PkVolume | Mean time-frequency volume of assigned TF-peaks | $sec * \mu V^2$ |
| PkArea | Mean time-frequency area of assigned TF-peaks | $sec * Hz$ |
| PkPeakiness | Mean TF-peak sharpness: $10 * \log_{10}(\text{Area} * \text{Height} / \text{Volume})$ | dB |
| PkSOpower | Mean normalized SO-power at the peak center times of assigned TF-peaks | SO power (dB) |
| PkSOphase | Circular mean SO-phase at the peak center times of assigned TF-peaks | rad |

The second approach of dimensionality reduction performs 2D spline interpolation to reconstruct the 2D histograms (Figure 6a), giving a flexible approximation of the multi-dimensional structure of TF-peak modes with still fewer parameters than the original, albeit being less interpretable (Figure 6b&c). Details of feature histogram fitting using spline basis functions are described in Appendix 11. With either of the above two approaches, overall structures of TF-peak distributions on the SO-power (Figure 6b) and SO-phase (Figure 6c) histograms are captured by a small number of variables, which can be used to test how dynamics of transient oscillations vary as a function of frequency and cortical brain states.

#### Part 3 Summary

With feature histograms constructed, DYNAM-O has obtained highly distilled characterizations of TF-peak distributions and their feature properties, which capture consistent patterns of sleep neural dynamics based on how often a certain type of TF-peaks occur during overnight sleep. These SO feature histograms can be easily visualized to compare between conditions as well as summarized as modes of TF-peaks. The next section offers strategies for conducting statistical tests on TF-peak dynamics using the novel representations provided by feature histograms. This distributional approach aims to derive data-driven insights into sleep physiology and identify patterns that may be overlooked by conventional sleep EEG measures.

### Part 4: Statistical Tests on TF-peak Dynamics

The outputs from DYNAM-O are naturally suited for statistical tests in different forms. Below we describe two flavors of statistical tests that are currently supported in the DYNAM-O Toolbox. Many other hypothesis testing tools may be suitable for different analytic goals.

#### Feature and mode analysis

First, researchers can perform hypothesis testing after aggregating discrete events using the table of TF-peaks and their feature properties, as long as one is willing to accept assumptions to define a priori cutoff thresholds. This approach is identical to how sleep spindles are often studied in conventional sleep EEG analyses, except now extending to general TF-peaks. Distributional comparisons such as Kolmogorov-Smirnov tests [82] can also be conducted to compare one or more feature property distributions between conditions or cohorts. Second, feature histograms provide a more efficient characterization to perform statistical tests while being more robust to noise. Analogous to the assumptions in selecting and averaging discrete TF-peaks, one could define regions of interest (ROIs) on feature histograms to compute mean densities for each region and compare these researcher-defined modes based on prior domain knowledge and/or visual inspections of histograms from the control group (Figure 7a). Note that one should not define ROIs after inspecting the differences in histograms to avoid biasing one’s statistical testing with implicit inferences. Third, while this “top-down” approach can be highly effective with known EEG changes in certain frequency ranges [34], we advocate for a “bottom-up” analysis using empirical modes identified through the unsupervised dimensionality reduction techniques introduced in DYNAM-O. This mode-based approach abstracts beyond individual oscillation events and bounds, which is more appropriate when lacking strong ROI priors on feature histograms. We showcase an example analysis using this approach in the **Results** section.

**Figure 7.**
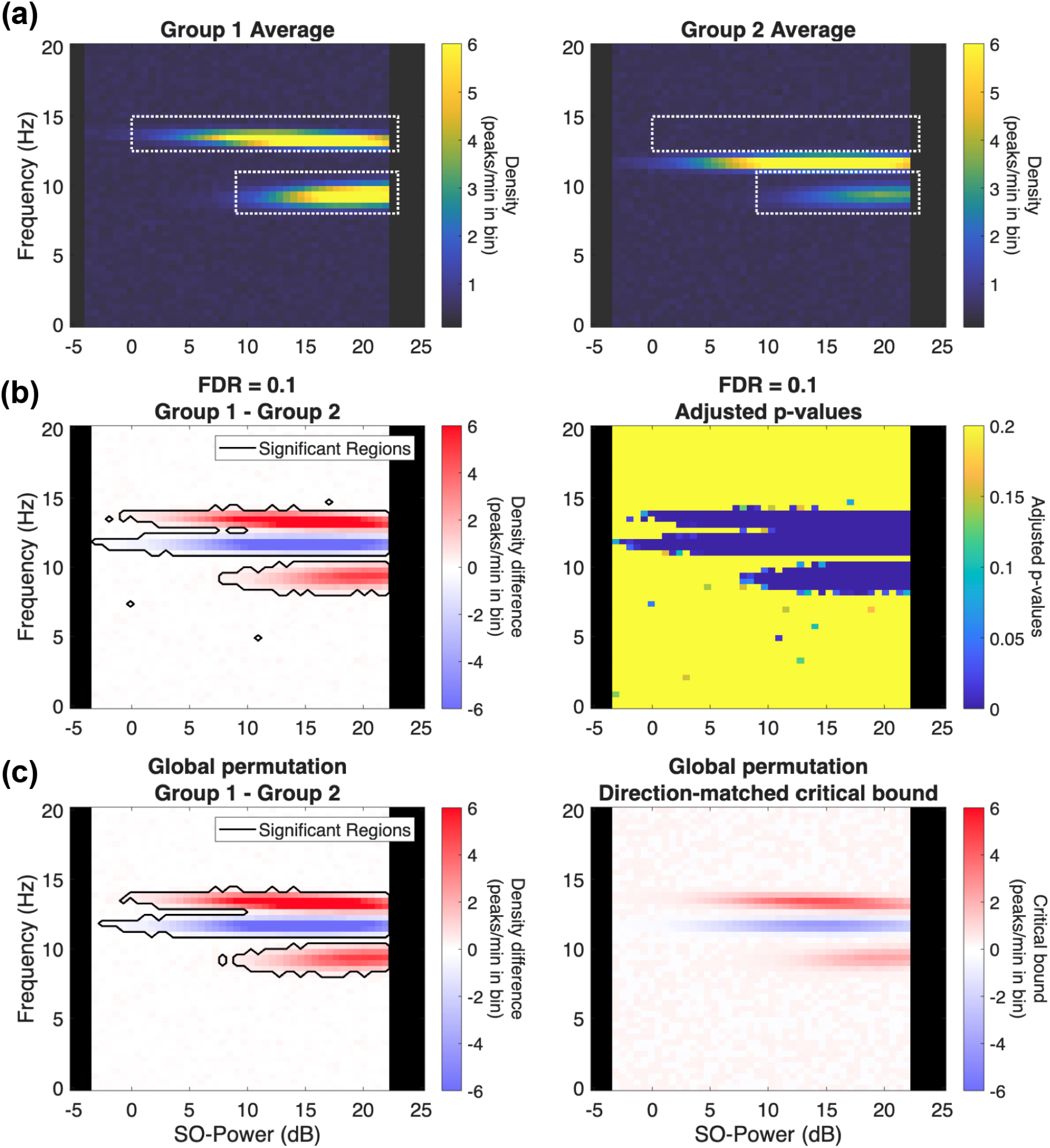
Simulated SO-power histograms demonstrated with two different techniques of statistical testing to detect group differences on feature histograms. **(a)** Group-averaged SO-power histograms for two simulated groups (Group 1 and Group 2), each displaying TF-peak density as a function of peak frequency and SO power; outlined boxes mark example researcher-defined regions of interest (ROIs) for a top-down comparison. **(b)** Bin-wise false discovery rate (FDR) analysis. Left: the Group 1 minus Group 2 difference histogram, with outlined regions surviving FDR control at 10%. Right: the corresponding map of adjusted p-values. **(c)** Global permutation testing on the linearized histograms. Left: the same difference histogram with significant regions identified by the permutation procedure. Right: the spatially varying acceptance bounds against which observed differences are compared. Actual bounds are continuous and symmetric in positive and negative values; displayed bounds are directionally matched to observed differences on the left such that darker color than the right panel indicates significant differences.

#### Whole-histogram analysis

Given multiple feature histograms from different individuals and/or repeated testing, we can also conduct a “whole-histogram” analysis (akin to whole-brain analysis of voxels in fMRI) to conduct group comparisons of sleep dynamics using the efficient representations encoded by feature histograms. Next, we briefly describe two general techniques to handle multiple comparisons that have been implemented within DYNAM-O to facilitate group comparisons of TF-peaks using the feature histogram outputs. The first technique conducts two-sample or paired-sample statistical tests at each bin on the histograms while controlling liberally at 10% false discovery rate (FDR) for exploratory analyses using the Benjamini-Yekutieli procedure for dependent tests [83]. One can update the FDR level to 0.05 to follow more standard statistical conventions. The second technique performs a global permutation testing on the linearized feature histograms based on the numbers of bins exceeding one-dimensional acceptance bounds [84], which provides greater sensitivity to detect small but consistent differences between feature histograms. Figure 7b and c show an illustrative example with the two techniques to conduct whole-histogram analysis, applied to two groups of simulated SO-power histograms.

## Results

Having explained the DYNAM-O processing pipeline, we now demonstrate actual outputs from the toolbox to characterize transient oscillatory neural dynamics during sleep. Throughout the sections below, we use code snippets and example data outputs to illustrate key function calls following the MATLAB implementation with a Rust backend for computation acceleration. The pure MATLAB DYNAM-O Toolbox exposes the same functions. While the Python implementation differs by adopting object-oriented programming conventions, the same functionalities are available with closely matching outputs (< 1% difference in TF-peak counts).

In the following sections, we first present the main function to run the full DYNAM-O processing pipeline and how to interpret its outputs. We then dissect the core function within DYNAM-O to extract TF-peaks from EEG data along with measures of each TF-peak event, which can also be used independently outside the DYNAM-O pipeline. We then explain the SO feature histograms constructed from TF-peaks, how to interpret them, and how to reduce the dimensionality of these histograms for efficient statistical testing. Finally, we conduct an illustrative group comparison analysis on sex differences to showcase how to integrate the various functions together to draw inferences on subtle changes in neural dynamics during sleep under a multi-dimensional distributional approach. Readers are encouraged to follow the sections with DYNAM-O installed to explore the introduced functions on the example data distributed with the toolbox.

### Using the DYNAM-O Toolbox to analyze overnight EEG from a recording channel

Programmatically, the entry-point function for the DYNAM-O Toolbox is `runDYNAMO()`. An overnight sleep recording yields a time trace of EEG signals (`data`) and its sampling frequency (`Fs`). Scoring of sleep stages by polysomnography technologists or automated scoring programs generates epochs of sleep stages and associated timing (`stage_vals, stage_times`). With these four input variables, `runDYNAMO()` can be called directly to analyze sleep EEG data from a single channel. The simplest use case of the function is as following:

~~~
[stats_table, …, SOPHs, …] = runDYNAMO(data, Fs, stage_times, stage_vals)
~~~

Figure 8 provides an algorithmic flow chart of the encapsulated function calls that are executed in sequence for a standard DYNAM-O processing pipeline. `runDYNAMO()` takes as inputs an overnight EEG time trace from a channel (`data`), its sampling frequency (`Fs`), and scored sleep stages. It performs all the computations of Parts 1-3 described in the **Methods** section: TF-peaks are identified on multitaper spectrograms estimated from the input data; feature properties of each of these identified TF-peaks are then computed; SO feature histograms are generated to capture the overnight dynamics of TF-peaks, and they are parameterized with a small number of coefficients. All detected TF-peaks and their computed features, including peak time and peak frequency, are returned as a table in `stats_table`. Generated feature histograms and their parameterization are included in `SOPHs`. Detailed results on the TF-peaks (`stats_table`; Parts 1-2 of the **Methods** section) and feature histograms (`SOPHs`; Part 3 of the **Methods** section) are described in later sections.

**Figure 8.**
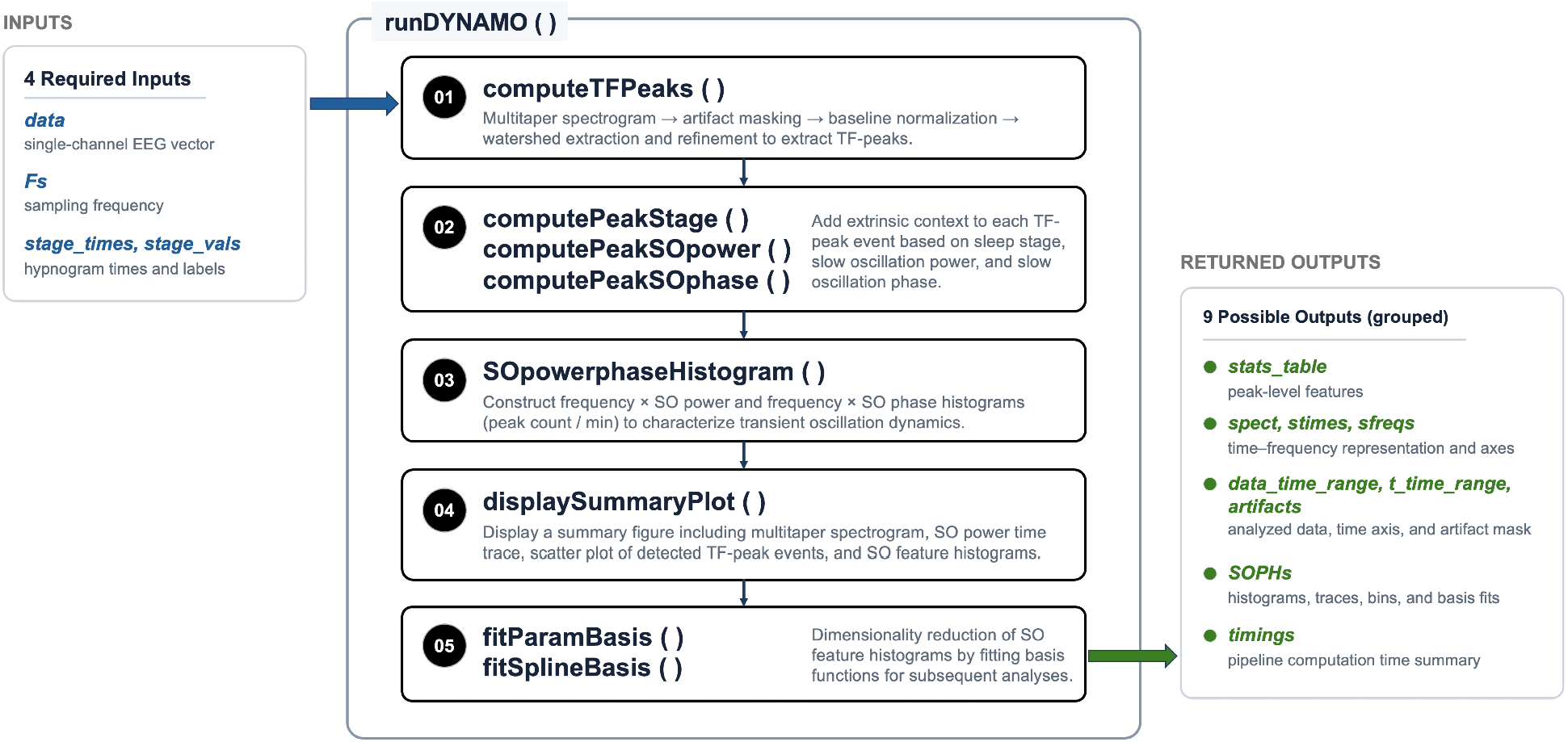
Algorithmic flow chart of the runDYNAMO() function with required inputs and all possible outputs. This entry-point function orchestrates the core parts of the DYNAM-O processing pipeline and calls relevant downstream computation methods to extract TF-peaks, calculate extrinsic TF-peak feature properties, construct SO feature histograms, and fit parametric and spline basis functions to represent the histograms using a handful of modes and coefficients. A summary figure is also produced using a dedicated function to visualize key pipeline outputs. Note that the `computeTFPeaks()` function is invoked first and depends only on the input variables. Therefore, it can be used as a general-purpose method to extract TF-peaks from any time series data beyond sleep EEG recordings.

A summary figure is automatically generated to display key findings by DYNAM-O on the input data across the multiple processing steps within `runDYNAMO()`. One typical summary figure obtained from DYNAM-O using the example data is shown in Figure 9, which can be easily reproduced by calling `runDYNAMO(’night’)`. The summary figure is divided into three subpanels organized from top to bottom.

**Figure 9.**
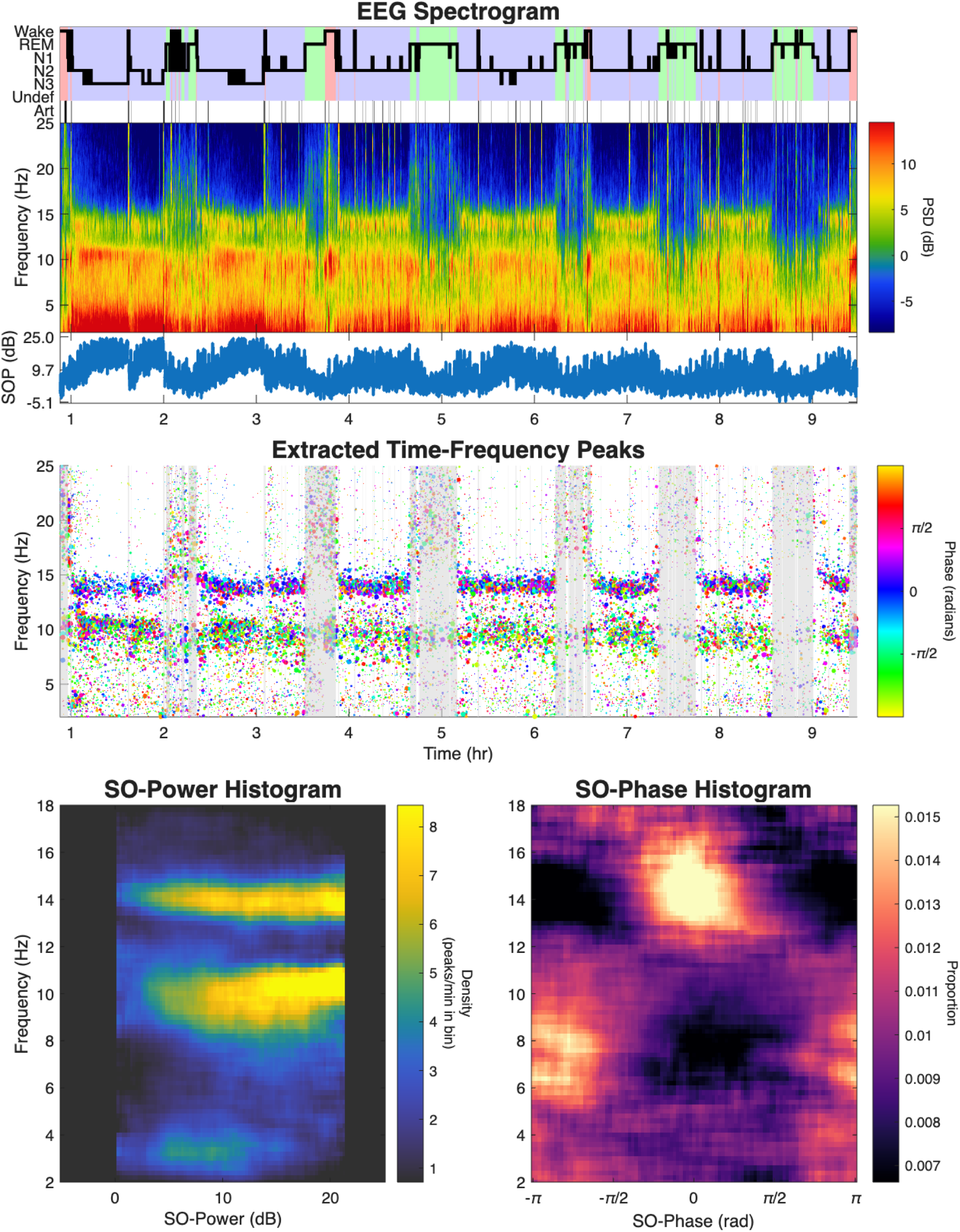
**Summary figure of DYNAM-O processing on one overnight sleep EEG recording,** organized into three stacked panels. The top panel shows the scored hypnogram, the raw multitaper spectrogram, and the continuous SO-power time series (a proxy for depth-of-sleep), giving a whole-night view of the single-channel recording in place of individual 30 s epochs. The middle panel shows all TF-peaks as scatter points, with marker size scaled by TF-peak volume and with marker color denoting the SO phase at each peak’s weighted centroid time. TF-peaks not included in the SO feature histograms are masked by semi-transparent vertical gray bands corresponding to wake, REM, and artifact periods that are excluded by the default NREM-only histograms. The bottom panel shows the resulting SO-power histogram (left) and SO-phase histogram (right), in which hotspots of warmer color mark the frequency and SO-power/phase regions where TF-peaks recur most densely across the night.

The first panel shows a standard hypnogram depicting scored sleep stages throughout the night, a raw estimated multitaper spectrogram, and a time series of computed slow oscillation power (SO-power), which gives a proxy continuous measure of depth-of-sleep. This panel provides an efficient visualization of the single channel recording over the entire night, in stark contrast to the traditional display of sleep EEG time traces in 30 s epochs. Both macrostructures such as sleep cycles and microstructures such as bouts of sleep spindles during NREM sleep can be easily seen on the spectrogram [38]. In addition, this view can highlight apparent issues such as abnormal scored sleep architecture (low sleep efficiency), artifacts in the recording, and potential errors during data preprocessing.

The second panel shows detected TF-peaks as scatter points, spanning across the recording duration and frequency range. The sizes of TF-peak scatter points are scaled by the volumes of extracted TF-peaks to accentuate strong transient oscillations. In addition, the TF-peak scatter points are colored according to the phases of slow oscillation at the peak time of each TF-peak, which can reveal clustering of transient oscillations at distinct frequencies within specific slow oscillation phases. It should be noted that only TF-peaks included in generating the SO feature histograms in the next panel are displayed in this scatter plot to allow matching visualizations. The default SO feature histograms focus on NREM sleep, with gray regions in indicating time periods during wake, REM sleep stages, or detected artifact periods that remain unused in the analyses.

The third panel shows two feature histograms focused on TF-peak dynamics in terms of slow oscillations, specifically slow oscillation power (SO-power) and slow oscillation phase (SO-phase). The histograms are generated by computing the densities of TF-peaks (counts per minute) within each bin of a SO-power/phase range and a frequency range, pooled across the night (default limits to NREM sleep) as SO-power/phase vary over different values. The densities under SO-phase histograms are normalized as proportions at a frequency (across a row). This panel highlights notable patterns in the occurrence of transient oscillations relative to slow oscillation properties. Hotspots on the feature histograms indicate increased numbers of detected TF-peaks within regions in the frequency x SO-power/phase space, characterizing consistent dynamics of transient oscillation activity across the night and providing one representation of stable features in sleep EEG. The SO feature histograms are also parameterized during `runDYNAMO()` to reduce the number of variables needed to capture prominent patterns on the histograms. We describe more details on the SO feature histograms and their dimensionality reduction in a later section.

There are several options that can be set when calling `runDYNAMO()` in order to adjust the various processing steps in DYNAM-O. The setting options are conveniently organized into five callable sets: `baseline_opts()`, `detection_opts()`, `SOpowerphasehist_opts()`, `param_basis_opts()`, and `spline_basis_opts()`. All default values can be invoked by directly calling the option functions. We explain each of these options in later **Results** sections during detailed results of TF-peak extraction and feature histogram analysis.

We recommend users to start with the default options, as they have been tested on a number of large datasets in NSRR [50,85]. In the most general use cases, one can apply `runDYNAMO()`on sleep EEG recordings with default settings to 1) obtain an in-depth visualization of individualized brain states during sleep as reflected by tens of thousands of transient oscillations in EEG, and 2) extract data-driven measures of these transient oscillations in output tables of TF-peak events and efficient histogram representations of their dynamics.

### Characterizing tens of thousands of transient oscillation events in sleep EEG

We now break down each of the key results obtained from running the top-level toolbox function `runDYNAMO()` and explain finer controls that can be exerted on each part of the processing steps. Detailed descriptions of all parameters and their defaults are available in our open-source code distribution.

First and foremost, the DYNAM-O Toolbox identifies TF-peaks from spectrograms and computes various peak properties to characterize transient oscillations during sleep. The primary TF-peak extraction function invoked by `runDYNAMO()` is `computeTFPeaks()`, which can also be called independently. Its simplest use case is as following:

~~~
[stats_table] = computeTFPeaks(data, Fs, stage_times, stage_vals)
~~~

`computeTFPeaks()` takes the same inputs as `runDYNAMO()` and computes TF-peaks as well as their features based on spectrogram estimation from the entire input EEG recording. Transient oscillation events with brief durations and narrowband frequencies will manifest as distinct local peaks in the time-frequency landscape of a spectrogram, when they are strong enough to be above the level of background noise. Accordingly, `computeTFPeaks()` employs the watershed algorithm to detect and outline these transient oscillation events in terms of individual TF-peaks. Figure 10a illustrates the boundaries of TF-peaks detected from the example sleep EEG data, zoomed into a short segment and overlaid on top of a spectrogram for display. Some of the TF-peaks correspond to well-known fast and slow sleep spindles, while others are weaker and have peak frequencies in the alpha/theta ranges. Less is known about the neural origins of the latter types of TF-peaks, but they are nevertheless highly discernible on spectrograms and indeed reliably identified by the DYNAM-O Toolbox.

**Figure 10.**
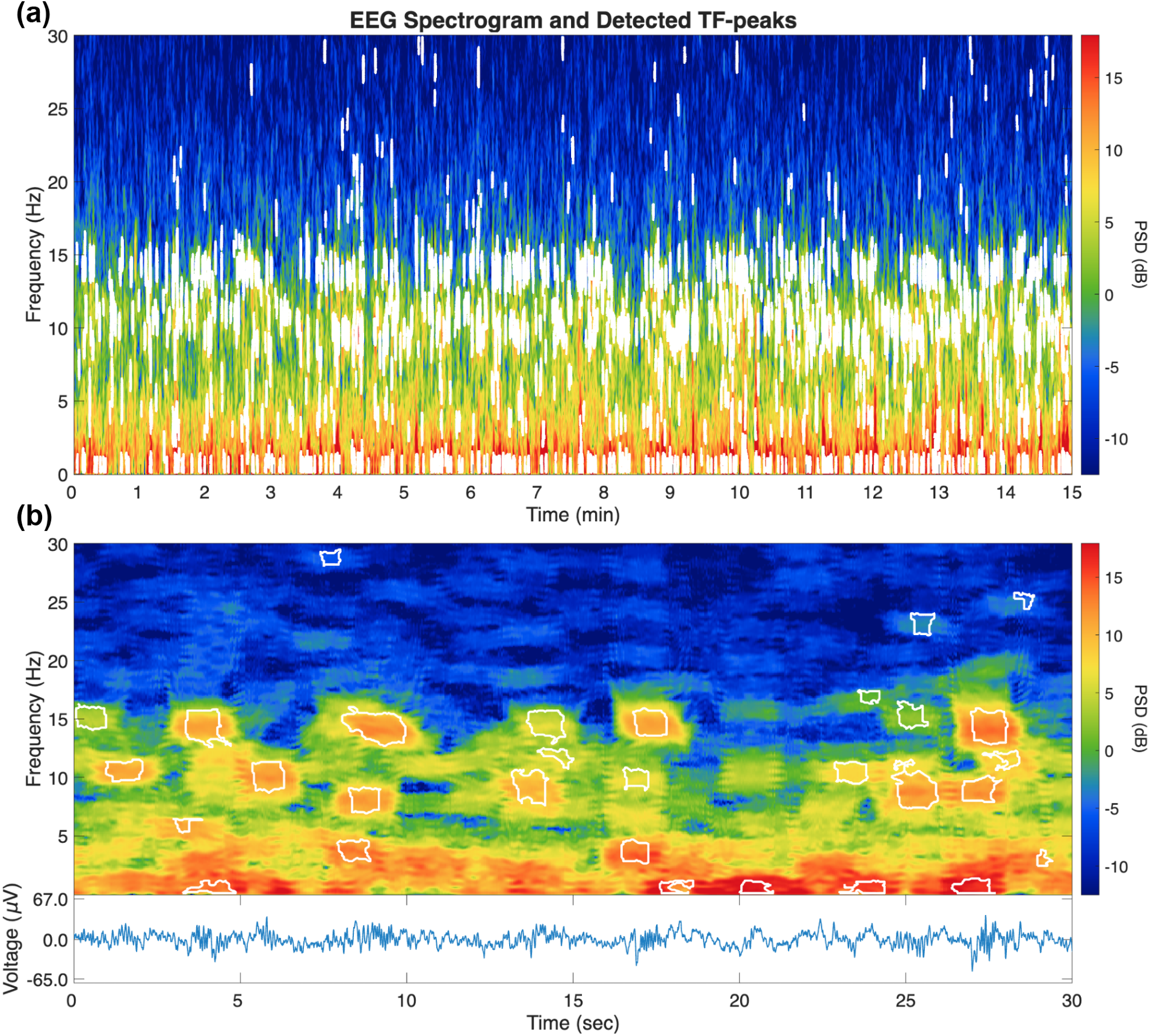
Boundaries of detected TF-peak events from overnight EEG during stage 2 sleep. **(a)** Detected TF-peaks across a 15 min window of stage 2 sleep, shown as boundaries overlaid on the unnormalized, finer-spectral-resolution multitaper spectrogram. **(b)** A 30 s segment from the same recording, with TF-peak boundaries overlaid on the spectrogram (top) and the corresponding EEG voltage trace shown below. Drawn after trimming each peak to 80% of its volume, the boundaries tightly outline transient oscillation events of differing morphology.

While `computeTFPeaks()` expects inputs of scored sleep staging, one can use this function to analyze time series data from any field with transient oscillation events. Users are encouraged to use the interactive viewer in DYNAM-O to inspect detected TF-peaks overlaid on the spectrogram to understand the forms of TF-peaks being captured. This important exploratory step can be invoked with the name-value pair `(’display_peaks’, true)` when using `computeTFPeaks()`. We recommend that users run `computeTFPeaks()` and display detected TF-peaks first in a handful of recordings before following later sections that describe characterizing TF-peak features and dynamics. This visualization can highlight promising analytic directions as well as flag any limitations of TF-peak detection in a given dataset.

It should be noted that the spectrogram being overlaid when displaying TF-peaks is only one of the two spectrograms (finer spectral resolution, see **Methods** section **Sequential optimization for temporal and spectral resolutions**) in the TF-peak computations. It is also the unnormalized version to match typical spectrogram visualization of sleep EEG, while the watershed algorithm is applied to baseline-corrected spectrograms with the 1/f background trend removed in order to isolate distinctive oscillations (see **Methods** section **TF-peak detection**). Finally, the TF-peak boundaries depict detected TF-peak events after trimming to 80% of the peak volume (see **Methods** section **TF-peak detection**), which compensates for smearing from spectrogram estimation and less reliable peak edges from merging. Despite these subtle differences, the displayed TF-peak boundaries provide highly accurate outlining of transient oscillation events being detected from the input data, and they give an informative visualization of this fundamental TF-peak detection step in the DYNAM-O Toolbox.

In a typical overnight sleep EEG recording lasting several hours, tens of thousands of TF-peaks are identified. As can be seen even from the short segment in Figure 10b, different TF-peaks often have distinct morphologies, with some being stronger and lasting longer while others being weaker and brief. Therefore, it is important to extensively characterize detected TF-peaks in terms of different properties, as they likely contain useful information about sleep physiology and neural dynamics. To this end, `computeTFPeaks()` outputs computed features of TF-peaks in an organized table `stats_table`. All intrinsic features listed in Table 1 are outputted by default as separate columns. In this tabulated sheet, each row corresponds to one detected TF-peak event. In addition to usual properties used to describe sleep microstructure events such as peak frequency, duration, and bandwidth, we also provide three different size measures of TF-peaks: Height (PSD), Area (extent), and Volume (both). Height measures the PSD distance from the peak to the noise floor, while Area is the two-dimensional area on the time-frequency spectrogram regardless of spectral power. Volume considers both PSD and area on spectrogram by calculating the total volume of the TF-peak in 3D. In addition, by default settings we provide the raw height data across all pixels in the region detected in a TF-peak, which could be used for further analysis of TF-peak morphology. Table 3 shows the first 10 rows of `stats_table` from running `computeTFPeaks()` on the full overnight example data. Note that TF-peaks detected from the entire recording are outputted in this table, including those identified from artifact periods, as users could be interested in studying artifacts using TF-peaks. Additional peak feature properties such as slow oscillation features and scored sleep stages are appended using functions `computePeakStage()`, `computePeakSOpower()`, and `computePeakSOphase()` to the output table. In the next section, we describe these three extrinsic features as they pertain to the generation of SO feature histograms.

**Table 3.** Feature properties of TF-peaks are provided in the stats_table output.

|  | PeakTime | PeakFrequency | Duration | Bandwidth | Height | Volume | SegmentNum | BoundingBox |  |  |  | Area | Peakiness | Boundaries | HeightData | PeakStage | SOpower | SOphase |
| --- | --- | --- | --- | --- | --- | --- | --- | --- | --- | --- | --- | --- | --- | --- | --- | --- | --- | --- |
|  |  |  |  |  |  |  |  | 1 | 2 | 3 | 4 |  |  |  |  |  |  |  |
| 1 | 3.1816e+03 | 28.1732 | 0.8500 | 1.8555 | 17.1806 | 27.0071 | 1 | 3.1813e+03 | 27.3438 | 0.8500 | 1.8555 | 0.9521 | -2.1773 | 67×2 double | 195×1 double | 6 | NaN | NaN |
| 2 | 3.1820e+03 | 18.1982 | 0.6000 | 1.2695 | 11.1660 | 5.3652 | 1 | 3.1818e+03 | 17.5781 | 0.6000 | 1.2695 | 0.4395 | -0.3878 | 40×2 double | 90×1 double | 6 | NaN | NaN |
| 3 | 3.1827e+03 | 23.4660 | 1.2500 | 1.8555 | 15.2074 | 39.5183 | 1 | 3.1821e+03 | 22.6562 | 1.2500 | 1.8555 | 1.5723 | -2.1822 | 79×2 double | 322×1 double | 6 | NaN | NaN |
| 4 | 3.1832e+03 | 26.5768 | 0.8500 | 2.5391 | 18.1047 | 28.1141 | 1 | 3.1827e+03 | 25.7812 | 0.8500 | 2.5391 | 1.4697 | -0.2390 | 74×2 double | 301×1 double | 6 | NaN | NaN |
| 5 | 3.1842e+03 | 26.2757 | 0.5500 | 1.7578 | 22.1722 | 4.2756 | 1 | 3.1840e+03 | 26.1719 | 0.5500 | 1.7578 | 0.2783 | 1.5936 | 47×2 double | 57×1 double | 6 | NaN | NaN |
| 6 | 3.1847e+03 | 20.6135 | 0.5000 | 1.0742 | 9.1083 | 6.1399 | 1 | 3.1844e+03 | 19.8242 | 0.5000 | 1.0742 | 0.4248 | -2.0054 | 32×2 double | 87×1 double | 6 | NaN | NaN |
| 7 | 3.1862e+03 | 26.1295 | 0.9000 | 1.8555 | 10.8057 | 25.0774 | 1 | 3.1858e+03 | 25.1953 | 0.9000 | 1.8555 | 1.0156 | -3.5890 | 59×2 double | 208×1 double | 5 | 0.3234 | 0.3148 |
| 8 | 3.1867e+03 | 23.3248 | 1.1000 | 2.5391 | 25.6077 | 51.8662 | 1 | 3.1862e+03 | 21.9727 | 1.1000 | 2.5391 | 1.9287 | -0.2125 | 87×2 double | 395×1 double | 5 | 0.3234 | 1.2481 |
| 9 | 3.1873e+03 | 18.5806 | 2.0500 | 2.5391 | 18.2603 | 74.2170 | 1 | 3.1864e+03 | 17.6758 | 2.0500 | 2.5391 | 3.2275 | -1.0012 | 117×2 double | 661×1 double | 5 | 0.3234 | 2.0511 |
| 10 | 3.1874e+03 | 28.8268 | 0.9000 | 1.8555 | 17.2384 | 24.7467 | 1 | 3.1870e+03 | 27.9297 | 0.9000 | 1.8555 | 0.9814 | -1.6515 | 66×2 double | 201×1 double | 5 | 0.3234 | 2.0748 |

The computations within `computeTFPeaks()` are also controlled using various parameters organized into two sets, specified by `baseline_opts()` and `detection_opts()`. The first set of parameters impact how the estimated spectrogram gets normalized prior to the watershed algorithm via removing the spectral slope (see **Methods** section **TF-peak detection**). Users can also exclude additional periods from being used for baseline spectrum calculation. The second set of parameters control granular settings of the TF-peak detection process (see **Methods** section **Part 1: Identifying TF-peaks from Spectrograms**), including whether to sequentially use two spectrograms with different optimized resolutions, multitaper parameters for spectrogram estimation, downsampling of spectrogram segments for faster processing speed, merging of over-segmented TF-peak regions, trimming of TF-peaks by volume, maximum duration and bandwidth allowed for detected TF-peaks, and whether to refine peak frequency estimation using a Hann window. Note that the minimum peak duration, bandwidth, height used to define valid TF-peak events are theoretically grounded by MTM parameters, which determine the minimum temporal and spectral resolutions. Therefore, these minimum thresholds (unlike the maximum) are not exposed for user adjustment to prevent TF-peak extraction that would be technically incorrect. All other parameters have default values that have been extensively tested on multiple NSRR [50] datasets of sleep EEG recordings and hence should provide effective and generalizable detection of TF-peaks across studies.

In general, we believe no fine tuning of these parameters is needed when the DYNAM-O Toolbox is used to analyze human sleep EEG from overnight recordings under standard clinical sleep lab environments or mobile EEG setups. Apart from edge cases with substantial electrical noise that may manifest as discrete peaks in the time-frequency domain, `computeTFPeaks()` can be readily employed off the shelf to identify TF-peaks and to quantify their feature properties, providing an efficient characterization of transient oscillations during overnight sleep EEG recordings. Future work can also test for optimal parameters that can work effectively on rodent and non-human primate data of sleep EEG recordings that contain transient oscillations.

### Describing transient oscillation dynamics using feature histograms

With the table of TF-peaks provided by `computeTFPeaks()`, one can compute summary statistics and perform inferences regarding transient oscillations. For example, an average peak strength could be calculated for all TF-peak events whose peak frequency falls between 12-14 Hz, which can be used as a crude metric of fast spindle activity similar to other spindle measures used in the literature [68,86]. However, there are at least two important caveats to such approach, especially when applied to analyze all TF-peak events. First, without further analysis, it is not immediately clear how to best define frequency cutoffs to select subsets of TF-peaks, especially given that TF-peak properties change across sleep stages and show substantial inter-individual variabilities. Second, averaging feature properties across events assumes the importance of how an arbitrarily defined type of transient oscillations manifests *on average* in EEG data, but it overlooks any information about dynamics during sleep. For these reasons, we instead employ histograms to study the ensemble densities of TF-peaks along some dimensions that are intrinsically meaningful in terms of sleep physiology. In other words, in the absence of a ground-truth threshold to decide whether a transient oscillation is relevant, we focus on patterns of repeated occurrence of TF-peaks with similar properties. This distributional approach can be applied to high-dimensional states that encompass all possible peak properties, but as a starting point, DYNAM-O studies clusters of increased rates of TF-peaks along two dimensions during sleep: oscillation frequency and slow oscillation features. The former is known to differ between distinct electrophysiological activity; the latter provides a surrogate continuous measure of sleep depth, separates apart clinically established NREM sleep stages, and reflects global cortical states of excitation and inhibition. Throughout the DYNAM-O Toolbox, we refer to such distributional embeddings as SO feature histograms.

To obtain SO feature histograms, runDYNAMO() uses the SOpowerphaseHistogram() function in the DYNAM-O Toolbox, whose simplest use case is as following:

~~~
[SOpower_mat, SOphase_mat, SOpower_bins, SOphase_bins, freq_bins] = SOpowerphaseHistogram(data, Fs, TFpeak_freqs, TFpeak_times)
~~~

The peak frequencies and times of TF-peaks are obtained directly from the `stats_table`. The output variables specify the SO feature histograms, which are displayed as the lower part of the summary figure shown in Figure 9. The SO feature histograms provide an intuitive visualization of one particular form of transient oscillation dynamics: the frequency and depth of sleep (or cortical state) at which TF-peaks occur most frequently, which can be easily discerned as hotspots of warmer color on the histograms. Currently, the DYNAM-O Toolbox focuses on studying TF-peak dynamics during NREM sleep, as TF-peak events in NREM show more organized clustering compared to REM and WAKE periods. Nevertheless, these restrictions can be easily adjusted using parameters that control feature histogram generation (see below). All measures related to the SO feature histograms are returned by `runDYNAMO()` as a structure in `SOPHs`.

The SO-power histogram in Figure 9 indicates that there are several prominent clusters of TF-peak events, with one occurring sharply around 14 Hz frequency and spanning the majority period of NREM sleep, while the other occurring sharply around 10 Hz frequency and more frequently during deeper sleep. There is perhaps yet another diffuse cluster of TF-peaks around the alpha frequency range that can be seen during shallower NREM sleep. The first two forms of TF-peaks match well with known characteristics of fast and slow sleep spindles, while the third type is less understood as it has not been reported with traditional spindle detectors. Finally, there is also a weaker delta cluster centered at around 3Hz that may reflect delta waves distinct from slow oscillations [87].

Similarly, the SO-phase histogram in Figure 9 indicates that TF-peaks at around 15 Hz tend to exhibit a peak-max phase-density coupling with SO. There is also another cluster of TF-peaks in the low-alpha/theta range exhibiting trough-max phase-density coupling with SO. It should be noted that the bandwidth (along the frequency dimension) of prominent clusters may not match between the SO-power and SO-phase histograms. This is because values displayed on the SO-phase histograms are normalized across rows to show the proportion of TF-peaks at a specific frequency occurring within a SO-phase bin. Thus, the two SO feature histograms provide complementary information about transient oscillation dynamics during sleep.

As with other parts of the DYNAM-O Toolbox, processing in `SOpowerphaseHistogram()` can be finely controlled using parameters specified by `SOpowerphasehist_opts()`. These parameters dictate the estimation of SO power and SO phase, the sleep stages for which histograms are computed, and the binning resolutions of the SO-power and SO-phase histograms. We again recommend users to start with SO feature histograms generated using default settings in their own datasets, as the default values have been extensively tested across a wide range of recordings to yield high-quality representations of transient oscillation dynamics in sleep EEG data.

### Dimensionality reduction of feature histograms for statistical comparisons

Visualizing the SO feature histograms can already provide a lot of insights into transient oscillations, neural dynamics, and overnight sleep physiology. But a natural need for subsequent analyses with these feature histograms is to quantify the patterns more formally and to allow statistical tests. The raw feature histograms as returned by `runDYNAMO()` each contains about 10,000 pixels of density or proportion values. DYNAM-O provides two built-in ways to perform dimensionality reduction to capture the major patterns on the histograms with fewer parameters, which could be used for better powered statistical analysis and hypothesis testing. By default, `runDYNAMO()` has arguments `fit_param_basis` and `fit_spline_basis` both set to `true`, which will perform 1) parametric and 2) spline fitting of the histograms and return fitted results in the `SOPHs` output structure.

As detailed in Appendix 10, parametrizing SO-power histograms in DYNAM-O is achieved by fitting two-dimensional Gaussian peaks to the histograms on a baseline plane in order to quantify prominent modes using parameters that define each of these modes. A single mode is specified by six parameters: density amplitude, mean and standard deviations specifying the Gaussian densities along the frequency and SO-power dimensions, as well as a rotation angle that captures any positioning with a tilt on the histograms. In the case of SO-phase histograms, we use a von Mises-Gaussian function to accommodate circular phases. Modes are fitted iteratively by adding one mode at a time until achieving plateauing R^2^ values in explaining variances on histograms.

Fitting SO feature histograms uses the `fitParamBasis()` function to reduce the multi-dimensional distributions into parametric Gaussian modes. The simplest use case is as following:

~~~
SOPHs = fitParamBasis(SOPHs)
~~~

where fitted modes for both power and phase histograms are appended as sub-structures into the `SOPHs` output. Figure 6 shows the results of parametric fitting of SO feature histograms on the example data in DYNAM-O. Contours of the fitted 2D Gaussians are also displayed to highlight identified modes in Appendix 10. A total of 4 modes are identified on the SO-power histogram (Figure 6b): the first two modes at 10 Hz and 14 Hz range correspond approximately to the slow and fast spindle-like oscillation events. An overlapping cluster of slow spindle-like oscillations is also separated into a third mode at 9 Hz, extending to shallower sleep depth and slightly lower frequency. This mode potentially reflects a distinct form of low alpha activity that has been previously reported [34]. Finally, a weak delta cluster at 3 Hz is also identified. A total of 2 modes are identified on the SO-phase histogram (Figure 6c): the first mode captures peak-max phase-density coupling of transient oscillations at around 14 Hz with SO, while the second mode captures a trough-max coupling in the low-alpha/theta range, which we also identified previously by inspecting the output SO-phase histogram in the summary figure.

While parametric fitting of SO feature histograms yields satisfactory results in many cases, there could be complex TF-peak patterns that are not easily represented as Gaussian peaks but are nevertheless useful to quantify, as they may still contain valuable information regarding transient oscillations and neural dynamics. DYNAM-O provides a complementary non-parametric approach with two-dimensional spline curve fitting to feature histograms. This functionality is accessed through the `fitSplineBasis()` function, which has a parallel simple usage:

~~~
SOPHs = fitSplineBasis(SOPHs)
~~~

The spline curve fitting reduces the dimensionality of histograms from close to 10,000 pixels to about 100 spline coefficients. A default display of spline fitting of SO feature histograms generated by `runDYNAMO()` on the example data is shown in Figure 6. These fitted spline coefficients effectively capture the variations in the feature histograms at a coarser resolution. Figure 6b shows that as few as 140 spline coefficients can accurately recapitulate major patterns on SO-power histograms, while the smoother SO-phase histograms only require 77 parameters (Figure 6c). Statistical comparisons can be better powered with these smaller variable sets compared to the raw histogram values. Such non-parametric spline fitting could be useful when tracking changes in SO feature histograms over time in patient cohorts, whose feature histograms may not be easily parameterized as 2D Gaussian peaks by `fitParamBasis()`.

Both parametric and spline fitting procedures can be tuned using hyper-parameters. We organize these hyper-parameters into two sets that are easily accessible at `param_basis_opts()` and `spline_basis_opts()`, which take in `’power’` or `’phase’` as the first input to load default settings that can be subsequently overwritten. Similar to other parts of DYNAM-O, default parameters work well in many overnight sleep EEG datasets. However, some tuning of the watershed parameters and/or Gaussian peak criteria, such as parameter bounds used in the nonlinear least squares fit, might be needed depending on the nature of signal qualities and transient oscillation dynamics in specific datasets. When fitting SO feature histograms, DYNAM-O focuses on frequencies between 2-18 Hz, which represent the primary frequency range of interest for transient oscillations during sleep and can be adjusted using the option functions.

We recommend users to begin with parametric fitting of SO feature histograms for subsequent analyses, as it provides a stronger dimensionality reduction and can yield more interpretable fitted parameters. The spline fitting serves as a fallback to handle complex and noisy feature histograms that may be observed in patients. Both procedures are viable ways to quantify the dynamics of tens of thousands of transient oscillations summarized in the feature histograms, and both carry great potential to reveal novel insights into ensemble patterns of neural activity during sleep and how they change as a function of biological variables, such as aging.

### Group comparisons of transient oscillation dynamics

With feature histograms computed from multiple recordings, a natural next step is to conduct group comparisons to investigate changes in overnight dynamics of transient oscillations as functions of other biological/clinical variables. One approach is to conduct a whole-histogram analysis without assuming a priori mode structures and perform bin-wise statistical comparisons between feature histograms from two groups. We next perform an illustrative analysis on sex differences in transient oscillation dynamics to showcase this approach.

It has been previously described in the sleep literature that females exhibit different sleep spindle properties from age-matched males, including higher frequencies of fast spindles [68,88,89]. We applied the DYNAM-O Toolbox to PSG recordings from 133 subjects within the Cleveland Family Study [49] in the NSRR database [50], which were selected solely based on a typical young-adult age range (20-35). 14 subjects were excluded due to apnea/hypopnea index (AHI) exceeding 15 that suggests at least moderate severity of sleep apnea. Additional 7 subjects were excluded due to poor data quality with severe artifacts that interfered with TF-peak detection. The final analyzed sample included 67 female and 45 male subjects (age min/max = 20.22/34.97, mean ± standard deviation = 26.1 ± 4.3). All results held when including all 133 subjects, but we chose to report from the restricted sample for better interpretability. We first extracted TF-peaks and computed SO feature histograms using the C3 channel in these subjects. Figure 11a shows the group-averaged SO-power histograms for female and male subjects respectively. Outlined regions on the difference histogram indicate significant differences between females and males in TF-peak densities, as examined under Wilcoxon rank sum tests controlled for false discovery rates set at 0.05 with a conservative assumption of non-zero dependence across comparisons [83] (see **Methods** section **Part 4: Statistical Tests on TF-peak Dynamics**).

**Figure 11.**
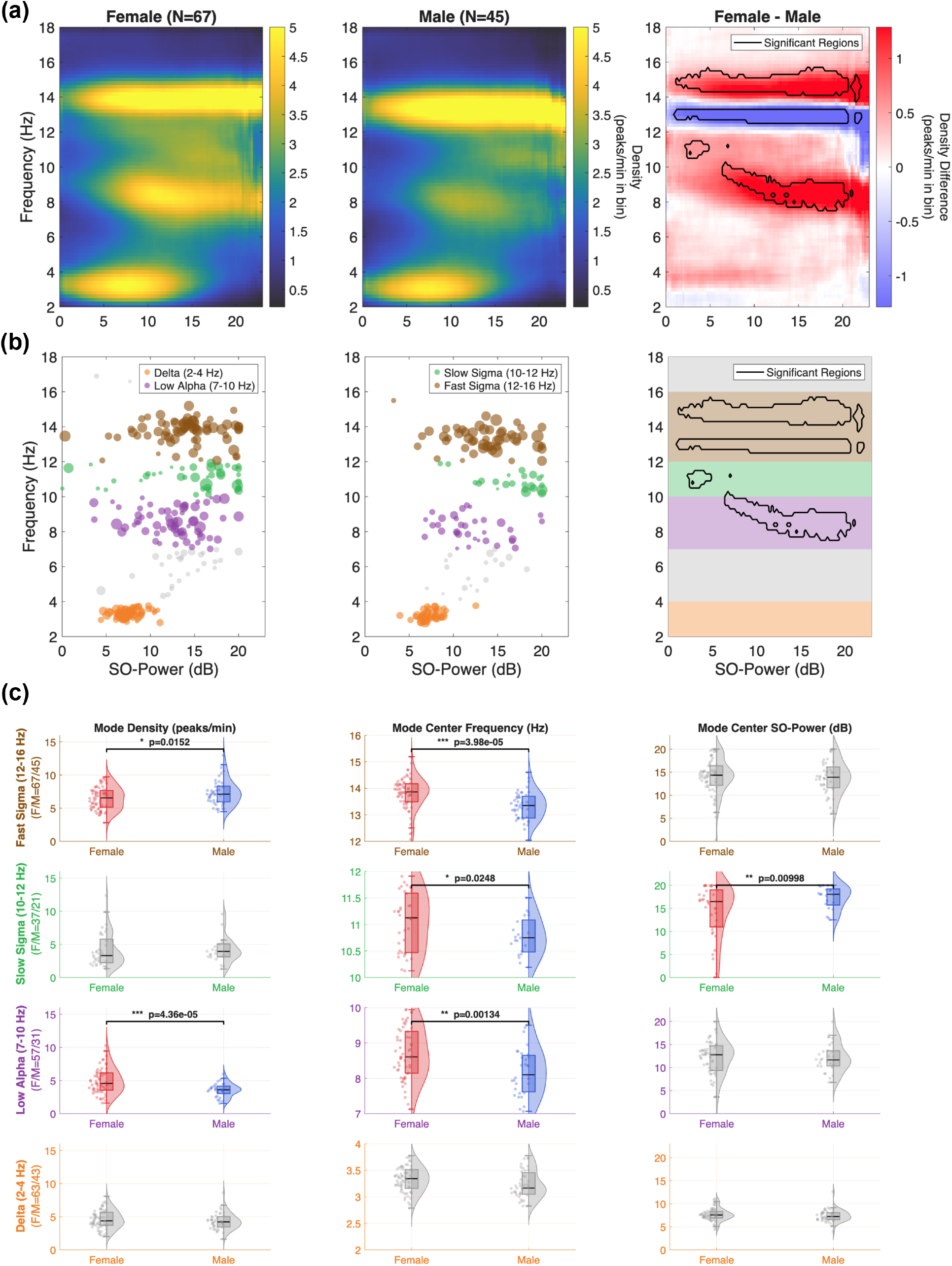
Sex difference analysis with whole-histogram and parameterized mode comparisons. **(a)** Group-averaged SO-power histograms for female (N = 67) and male (N = 45) subjects in the 20-35 age range from the Cleveland Family Study, as well as the female-minus-male difference histogram (right). Outlined regions on the difference histogram mark significant sex differences in TF-peak density (Wilcoxon rank-sum tests, FDR-controlled), including increased density near 15 Hz and decreased density near 13 Hz in females that together reflect a higher fast-sigma mode frequency centered at 14 Hz, along with increased low-alpha (7–10 Hz) activity in females. **(b)** Parameterized modes obtained by fitting two-dimensional Gaussian peaks to each individual’s SO-power histogram, plotted as mode center frequency versus center SO-power for female (left) and male (middle) subjects. Modes are colored following a frequency-based labeling scheme into delta (2-4 Hz), low alpha (7–10 Hz), slow sigma (10–12 Hz), and fast sigma (12–16 Hz); Outlined regions on top of shaded bands (right) denote the labeling ranges and the significant regions within each range. **(c)** Group comparisons of fitted mode parameters, including mode density, mode center frequency, and mode center SO power between females and males across the four TF-peak types. Raincloud plots show the group distributions of TF-peak mode measures in each of the four frequency ranges, and p-values denote significant sex differences based on two-sample t-tests uncorrected for multiple comparisons.

Two statistical testing functions have been implemented in DYNAM-O for whole-histogram group comparisons. First, the sex difference analysis illustrated above can be achieved using the function `FDR_2D()` with stacked feature histograms as inputs:

~~~
[sigbins, p_adj, p_values] = FDR_2D(group1_SOPHs, group2_SOPHs)
~~~

Second, global permutation testing (see **Methods** section **Part 4: Statistical Tests on TF-peak Dynamics**) can be achieved using the function gpermtest2() with the same inputs, whose results are omitted here for brevity.

From Figure 11a, one can observe that the fast sigma frequency mode (generalizing traditional fast spindles) has a slight but significantly higher mode frequency in females. This is reflected as a pair of increase/decrease in TF-peak densities on the difference histogram, where the shift in mode frequency centered at 14 Hz results in significantly increased density around 15 Hz and significantly decreased density around 13 Hz in females. Beyond this well-documented frequency change in fast sigma activity [90], additional regions of increased TF-peak densities are also identified by the FDR-group comparisons in the low alpha (7-10 Hz) and slow sigma (10-12 Hz) frequency ranges. Indeed, the low alpha difference is highly visible when comparing the averaged SO-power histograms, which show increased activity in females. In contrast, the sex difference in slow sigma frequency (generalizing traditional slow spindles) is much more focal and locates in a low SO-power region without strong TF-peak activity in either group. The theta (4-7 Hz) range is skipped, as no clear pattern of activity can be observed. Finally, a single strong mode is visible in the delta wave (2-4 Hz) range in both groups, although no sex difference reached FDR-controlled statistical significance. In summary, this example of whole-histogram analysis of sex differences demonstrates how DYNAM-O feature histograms can offer intuitive visualizations to spotlight group differences in multi-dimensional transient oscillation dynamics via simple 2D images.

In addition to whole-histogram analysis, one can leverage the powerful dimensionality reduction tools in DYNAM-O to perform group comparisons. To illustrate this analytic approach, we next employ mode parameters obtained from applying `param_basis_power()` to individual SO-power histograms and re-examine the sex differences in transient oscillatory activity. Figure 11b shows all Gaussian peaks extracted from the female and male subjects, each fitted with mode parameters including mode density, center frequency, and center SO power (see **Methods** section **Dimensionality reduction of feature histograms**). To test for sex differences, we adopt a frequency-based labeling of parameterized modes, where the strongest mode within 12-16 Hz is used to represent an individual’s fast sigma activity, 10-12 Hz for slow sigma, 7-10 Hz for low alpha, and 2-4 Hz for delta (Figure 11b). Events whose center frequency overlaps exactly with a frequency boundary is classified into the higher frequency mode. If an individual does not express a mode in a range, they are treated as missing data for that comparison. These four frequency ranges are drawn from where increased activity was observed in the grand-averaged histogram across all subjects (not shown) to roughly separate distinct modes of activity. While they are not circularly defined based on where significant differences were observed in the whole-histogram analysis, they are still post-hoc ROIs. It should also be cautioned that each individual’s SO-power histogram may not follow these frequency cutoffs. Nevertheless, this simple labeling scheme enables mode matching across subjects. More advanced labeling techniques such as data-driven clustering on mode parameters should be considered when typical structures of sleep transient oscillations may not hold or when a grand-averaged histogram across all subjects is not representative of groups in comparison, such as in developmental or patient populations.

Figure 11c compares females and males on mode density, center frequency, and center SO power across the four TF-peak types. Separate two-sample t-tests were conducted to contrast females and males on the four different mode types. The numbers of subjects included in each comparison are displayed in the y-axis label of each row, and displayed p-statistics are uncorrected for multiple comparisons. Overall, results from mode-based statistical tests strongly agree with the whole-histogram analysis, but they offer better clarifications on how histogram modes differ between groups. First, parameterized modes confirm that fast sigma activity (12-16 Hz) in females indeed shows higher frequencies than males. No difference is observed regarding the depth of sleep proxied by SO power. However, fast sigma mode density is higher in males compared to females, suggesting more TF-peaks are observed per minute in this frequency range in the males analyzed in this sample. Second, significant slow sigma (10-12 Hz) mode differences are also identified, but they substantially revise our earlier FDR-controlled whole-histogram analysis results: females exhibit higher frequency in slow sigma TF-peaks similar to the finding in fast sigma frequency, but slow sigma modes in males occur at deeper sleep indicated by the significantly higher mode SO power. No significant difference in mode density is found, despite the average histograms appear to show stronger activity in females. Third, females show much stronger low-alpha (7-10 Hz) activity than males, which likely indicates sex differences in a novel type of transient oscillations distinct from spindle-range activity. Interestingly, low-alpha mode frequencies are also elevated in females, consistent with the sigma-range findings. Finally, delta (2-4 Hz) frequency modes did not differ between groups, suggesting similar delta wave activity across females and males, which again agrees with the whole-histogram comparison results.

In summary, we have shown in this section that feature histograms provide efficient encoding of overnight sleep dynamics via TF-peaks. DYNAM-O offers a powerful framework to identify novel changes in transient oscillation dynamics during sleep that might have been overlooked due to constraints of traditional EEG measures. We have implemented the two group comparison methods in order to facilitate hypothesis testing and scientific discoveries. Other dimensionality reduction techniques such as principal component analysis and statistical methods such as support vector machine analysis are all relevant options to analyze feature histograms for studying changes in sleep oscillations. We aim to add support for these methods in the future.

## Discussion

Influential studies and major large data repositories of polysomnography (PSG) [50,91] have created an urgent need to automate the analysis of sleep EEG recordings. Making computerized measures of sleep oscillation features available is a critical step toward enabling sleep and neuroscience research at scale. Here, we provide a detailed introduction to the DYNAM-O Toolbox that supports the investigation of transient sleep oscillations within an efficient, data-driven, and rigorous framework for studying sleep physiology. This work builds on the foundation we introduced in [34], while providing key methodological improvements, computational optimizations, and software support for new built-in analysis methods. We present the four core parts of the DYNAM-O pipeline, which complete the full analysis cycle from extracting TF-peaks in raw data to computing summary statistics for scientific inference and hypothesis testing. We also describe visualization tools and modular processing steps that can be flexibly used as needed.

DYNAM-O has many potential use cases, suiting distinct analytic goals when studying sleep EEG data. Given the highly individualized and stable patterns of transient oscillations revealed by DYNAM-O, this approach can be especially well positioned to analyze longitudinal data and uncover subtle changes in sleep dynamics that may relate to health outcomes within an individual [92]. As detailed analyses of change trajectories on SO feature histograms are beyond the scope of the current software paper, we leave the relation of altered TF-peak modes to longitudinal observations within the same individual for future empirical studies. Next, we discuss four additional example scenarios where DYNAM-O could significantly advance the characterization of sleep dynamics and aid sleep research.

First, the multitaper spectrogram and SO feature histograms visualize an entire night worth of electrophysiology during sleep at a single glance. These visualizations are particularly useful in a clinical sleep lab for obtaining a general sense of a patient’s sleep physiology, or during exploratory analysis in sleep research studies to identify patterns in PSG data. In addition, there are often insidious artifacts from electrical noise sources in PSG that are difficult to observe on conventional time traces, spectral estimates, or even spectrograms. These systematic artifacts nevertheless last across nights and can severely hinder accurate data analysis. Such non-physiological regularities are often obvious on feature histograms, allowing researchers to troubleshoot data collection prior to committing substantial resources in large datasets.

Second, as demonstrated in the **Results** section **Group comparisons of transient oscillation dynamics**, DYNAM-O outputs can be directly used to compare between two cohorts such as males and females, patients and controls, or before and after treatment. Such comparisons can be achieved at multiple levels, including granular features of individual discrete TF-peak events, non-parametric whole-histogram analysis, and interpretable mode parameterization. Conducting group comparisons across all these levels gives a comprehensive depiction of changes in transient oscillation dynamics and can reveal previously unseen patterns of sleep physiology.

Third, DYNAM-O measures can also be used within a single cohort to model impacts of continuous covariates, such as biological age and levels of pathology from established biomarkers. This type of analysis is facilitated by the data-driven distributional approach DYNAM-O takes to quantify transient sleep oscillations: while some variability is expected from any quantitative measure, the reproducible DYNAM-O outputs, especially SO feature histograms that are largely invariant across multiple nights within an individual [34], provide a robust basis to investigate trends in sleep physiology as a function of continuous or ordered-categorical clinical variables.

Fourth, machine learning methods, including deep neural networks, have been recently applied to sleep EEG data to extract feature patterns and latent embeddings that achieved encouraging successes in predicting health outcome variables [93,94]. Unsurprisingly, many ML applications in sleep medicine [95–97] continue to use conventional EEG measures for feature extraction. Incorporating DYNAM-O features into existing ML frameworks could enhance the way in which these networks capture intersubject variability in a substantial way. Moreover, the recent adoption of transformer-based architectures led to the emergence of a series of *foundation models* trained using large PSG datasets [94,98]. Models like SleepFM [94] or SleepGPT [98] have been shown to very comprehensively represent brain wave signals using vector embeddings and obtain superior success in sleep-related pathology classification. These powerful models trade interpretability for dimensionality and inform scientific understanding by providing strong evidence for information content embedded within the data. Simultaneously, it is also important to explore orthogonal approaches, such as DYNAM-O, that conversely trade dimensionality for interpretability, to extract meaningful metrics of an individual’s neural dynamics during sleep. Future work combining both approaches could bridge the gap between these methodological advances.

For all of the above use cases, the DYNAM-O Toolbox can be fine-tuned using various parameters, but we recommend users to start with default parameters and exploit the variety of visualization tools provided by DYNAM-O to confirm analysis validity in a handful of pilot subjects first. A general workflow for studying sleep EEG recordings within DYNAM-O entails the following steps: *1) complete raw data export and sleep stage scoring*: this is usually done on clinical or research PSG systems, whose outputs are manually scored by trained polysomnography technologists or automated sleep stager; *2) apply DYNAM-O processing on each single-channel recording and obtain a summary figure*: inspect the spectrogram and scatter plots of extracted TF-peaks to verify performance and identify any data loading error, e.g., wrong sampling rate; *3) compare obtained SO feature histograms with previous DYNAM-O outputs*: explore normal or abnormal structures and utilize the TF-peak per-event boundary display as needed to investigate outstanding patterns; *4) batch analyze all sleep EEG data*: use saved option sets (`*_opts()`) to ensure identical parameter settings across runs and uniform output formats; *5) conduct group-level analysis*: leverage parameterized modes of SO feature histograms in conjunction with non-parametric whole-histogram analysis to perform statistical testing on transient oscillation dynamics.

Additionally, DYNAM-O is suited to applications beyond standard sleep EEG. Within EEG, this could be used to study transient EEG oscillation activity as a function of features ranging from drug effect site level, depth of consciousness, aperiodic slope, and many others. In general, DYNAM-O can be generalized to study any set of transient oscillations as a function of any continuous covariate.

We aim to expand on the DYNAM-O Toolbox in the near future with full GUI support as a standalone application for running all analysis steps described in this paper with native visualization tools, new functional modules to characterize event history timing information [22], and statistical methods for simulating and quantifying sleep EEG time series [99]. Ultimately, we believe a heavily-optimized and easily-accessible software is necessary to propel the sleep research field to further embrace data-driven analysis of PSG data and explore subtle patterns in neural dynamics during sleep.

## Acknowledgements

We would like to express our gratitude toward Dr. Dara Manoach for access to the data from Wamsley et al. for the example data in the DYNAM-O Toolbox. The Cleveland Family Study (CFS) was supported by grants from the National Institutes of Health (HL46380, M01 RR00080-39, T32-HL07567, RO1-46380). The National Sleep Research Resource was supported by the National Heart, Lung, and Blood Institute (R24 HL114473, 75N92019R002).

## References

1. Carskadon MA, Dement WC. Normal human sleep: an overview. Princ pract sleep med. 2005;4(1):13–23. doi:10.1016/B0-72-160797-7/50009-4

2. De Gennaro L, Ferrara M, Vecchio F, Curcio G, Bertini M. An electroencephalographic fingerprint of human sleep. Neuroimage. 2005;26(1):114–122. doi:10.1016/j.neuroimage.2005.01.020

3. Deak M, Epstein LJ. The history of polysomnography. Sleep Med Clin. 2009;4(3):313–321. doi:10.1016/j.jsmc.2009.04.001

4. Mahowald MW, Schenck CH. Insights from studying human sleep disorders. Nature. 2005;437(7063):1279–1285. doi:10.1038/nature04287

5. Stone JL, Hughes JR. Early history of electroencephalography and establishment of the American clinical neurophysiology society. J Clin Neurophysiol. 2013;30(1):28–44. doi:10.1097/WNP.0b013e31827edb2d

6. Šušmáková K. Human sleep and sleep EEG. Meas sci rev. 2004;4(2):59–74.

7. Campbell IG. EEG Recording and Analysis for Sleep Research. Current Protocols in Neuroscience. 2009;49(1). doi:10.1002/0471142301.ns1002s49

8. Carskadon MA, Dement WC. Monitoring and staging human sleep. Princ pract sleep med. 2011;5:16–26. doi:10.1016/B978-1-4160-6645-3.00002-5

9. Moser D, Anderer P, Gruber G, et al. Sleep classification according to AASM and Rechtschaffen & Kales: effects on sleep scoring parameters. Sleep. 2009;32(2):139–149. doi:10.1093/sleep/32.2.139

10. Berry RB, Brooks R, Gamaldo C, et al. AASM scoring manual updates for 2017 (version 2.4). 2017;13:5. doi:10.5664/jcsm.6576

11. Cox R, Fell J. Analyzing human sleep EEG: a methodological primer with code implementation. Sleep Med Rev. 2020;54:101353. doi:10.1016/j.smrv.2020.101353

12. Ujma PP, Dresler M, Bódizs R. Comparing manual and automatic artifact detection in sleep EEG recordings. Psychophysiology. 2025;62(2):e70016. doi:10.1111/psyp.70016

13. Fiorillo L, Puiatti A, Papandrea M, et al. Automated sleep scoring: a review of the latest approaches. Sleep med rev. 2019;48:101204. doi:10.1016/j.smrv.2019.07.007

14. Motamedi-Fakhr S, Moshrefi-Torbati M, Hill M, Hill CM, White PR. Signal processing techniques applied to human sleep EEG signals—a review. Biomed Signal Process Control. 2014;10:21–33. doi:10.1016/j.bspc.2013.12.003

15. Rosenberg RS, Van Hout S. The American academy of sleep medicine inter-scorer reliability program: sleep stage scoring. J Clin Sleep Med. 2013;9(1):81–87. doi:10.5664/jcsm.2350

16. Younes M, Raneri J, Hanly P. Staging sleep in polysomnograms: analysis of inter-scorer variability. J Clin Sleep Med. 2016;12(6):885–894. doi:10.5664/jcsm.5894

17. Younes M. The case for using digital EEG analysis in clinical sleep medicine. Sleep Sci Pract. 2017;1(1):2. doi:10.1186/s41606-016-0005-0

18. Cox R, Mylonas DS, Manoach DS, Stickgold R. Large-scale structure and individual fingerprints of locally coupled sleep oscillations. Sleep. 2018;41(12):zsy175. doi:10.1093/sleep/zsy175

19. Tarokh L, Carskadon MA, Achermann P. Trait-like characteristics of the sleep EEG across adolescent development. J Neurosci. 2011;31(17):6371–6378. doi:10.1523/JNEUROSCI.5533-10.2011

20. Tucker AM, Dinges DF, Van Dongen HP. Trait interindividual differences in the sleep physiology of healthy young adults. J Sleep Res. 2007;16(2):170–180. doi:10.1111/j.1365-2869.2007.00594.x

21. Bódizs R, Körmendi J, Rigó P, Lázár AS. The individual adjustment method of sleep spindle analysis: methodological improvements and roots in the fingerprint paradigm. J Neurosci Methods. 2009;178(1):205–213. doi:10.1016/j.jneumeth.2008.11.006

22. Chen S, He M, Brown RE, Eden UT, Prerau MJ. Individualized temporal patterns drive human sleep spindle timing. Proc Natl Acad Sci. 2025;122(2):e2405276121. doi:10.1073/pnas.2405276121

23. Chen S, Redline S, Eden UT, Prerau MJ. Dynamic models of obstructive sleep apnea provide robust prediction of respiratory event timing and a statistical framework for phenotype exploration. Sleep. 2022;45(12):zsac189. doi:10.1093/sleep/zsac189

24. Dimitrov T, He M, Stickgold R, Prerau MJ. Sleep spindles comprise a subset of a broader class of electroencephalogram events. Sleep. 2021;44(9):zsab099. doi:10.1093/sleep/zsab099

25. Grigg-Damberger MM. The AASM scoring manual: a critical appraisal. Curr Opin Pulm Med. 2009;15(6):540–549. doi:10.1097/MCP.0b013e328331a2bf

26. Schulz H. Rethinking sleep analysis: comment on the AASM manual for the scoring of sleep and associated events. J Clin Sleep Med. 2008;4(2):99–103. doi:10.5664/jcsm.27124

27. Loomis AL, Harvey EN, Hobart G. Potential rhythms of the cerebral cortex during sleep. Science. 1935;81(2111):597–598. doi:10.1126/science.81.2111.597

28. Himanen SL, Hasan J. Limitations of Rechtschaffen and Kales. Sleep med rev. 2000;4(2):149–167. doi:10.1053/smrv.1999.0086

29. Rechtschaffen A, Kales A. A manual of standardized terminology, techniques and scoring system for sleep stages of human subjects. US department of health, education and welfare. Public Health Serv Bethesda. Published online 1968. doi:10.1001/archpsyc.1969.01740140118016

30. O’Reilly C, Nielsen T. Automatic sleep spindle detection: Benchmarking with fine temporal resolution using open science tools. Front Hum Neurosci. 2015;9:353. doi:10.3389/fnhum.2015.00353

31. Schimicek P, Zeitlhofer J, Anderer P, Saletu B. Automatic sleep-spindle detection procedure: aspects of reliability and validity. Clin Electroencephalogr. 1994;25(1):26–29. doi:10.1177/155005949402500108

32. Warby SC, Wendt SL, Welinder P, et al. Sleep-spindle detection: crowdsourcing and evaluating performance of experts, non-experts and automated methods. Nat Methods. 2014;11(4):385–392. doi:10.1038/nmeth.2855

33. He M, Das P, Hotan G, Purdon PL. Automatic segmentation of sleep spindles: a variational switching state-space approach. In: 2022 56th Asilomar Conference on Signals, Systems, and Computers. IEEE; 2022:1301–1305. doi:10.1109/IEEECONF56349.2022.10052015

34. Stokes PA, Rath P, Possidente T, et al. Transient oscillation dynamics during sleep provide a robust basis for electroencephalographic phenotyping and biomarker identification. Sleep. 2023;46(1):zsac223. doi:10.1093/sleep/zsac223

35. Biabani N, Ilic K, Birdseye A, et al. Disorder-specific alterations of transient oscillatory dynamics during sleep across cortical and subcortical networks. Scientific Reports. 2026;16(1):3630. doi:10.1038/s41598-025-33669-1

36. Hermans LW, Huijben IA, van Gorp H, et al. Representations of temporal sleep dynamics: review and synthesis of the literature. Sleep Med Rev. 2022;63:101611. doi:10.1016/j.smrv.2022.101611

37. Phan H, Mikkelsen K. Automatic sleep staging of EEG signals: recent development, challenges, and future directions. Physiol Meas. 2022;43(4):4TR01. doi:10.1088/1361-6579/ac6049

38. Prerau MJ, Brown RE, Bianchi MT, Ellenbogen JM, Purdon PL. Sleep Neurophysiological Dynamics Through the Lens of Multitaper Spectral Analysis. Physiology. 2017;32(1):60–92. doi:10.1152/physiol.00062.2015

39. Roebuck A, Monasterio V, Gederi E, et al. A review of signals used in sleep analysis. Physiol Meas. 2014;35(1):R1–R57. doi:10.1088/0967-3334/35/1/R1

40. Perez-Pozuelo I, Zhai B, Palotti J, et al. The future of sleep health: a data-driven revolution in sleep science and medicine. npj Digital Med. 2020;3(1):42. doi:10.1038/s41746-020-0244-4

41. Redline S, Purcell SM. Sleep and big data: harnessing data, technology, and analytics for monitoring sleep and improving diagnostics, prediction, and interventions—an era for sleep-omics? 2021;44:6. doi:10.1093/sleep/zsab107

42. Vallat R, Walker MP. An open-source, high-performance tool for automated sleep staging. elife. 2021;10:e70092. doi:10.7554/eLife.70092

43. Purcell L, Karkera T, Palanivelu S, et al. The LUNA ecosystem: integrated tools for sleep signal visualization and analysis. Published online May 2026. doi:10.5281/zenodo.20025864

44. Donoghue T, Haller M, Peterson EJ, et al. Parameterizing neural power spectra into periodic and aperiodic components. Nat Neurosci. 2020;23(12):1655–1665. doi:10.1038/s41593-020-00744-x

45. Percival DB, Walden AT. Spectral Analysis for Physical Applications. Vol 909. Cambridge; 2002. doi:10.1017/CBO9780511622762

46. Groppe DM, Urbach TP, Kutas M. Mass univariate analysis of event-related brain potentials/fields I: a critical tutorial review. Psychophysiology. 2011;48(12):1711–1725. doi:10.1111/j.1469-8986.2011.01273.x

47. Kovesi P. Good colour maps: how to design them. arXiv. Preprint posted online September 12, 2015:arXiv:1509.03700. doi:10.48550/arXiv.1509.03700

48. Wamsley EJ, Tucker MA, Shinn AK, et al. Reduced Sleep Spindles and Spindle Coherence in Schizophrenia: Mechanisms of Impaired Memory Consolidation? Biological Psychiatry. 2012;71(2):154–161. doi:10.1016/j.biopsych.2011.08.008

49. Redline S, Tishler PV, Tosteson TD, et al. The familial aggregation of obstructive sleep apnea. Am J Respir Crit Care Med. 1995;151(3):682–687. doi:10.1164/ajrccm/151.3_Pt_1.682

50. Zhang GQ, Cui L, Mueller R, et al. The National Sleep Research Resource: towards a sleep data commons. J Am Med Inf Assoc. 2018;25(10):1351–1358. doi:10.1093/jamia/ocy064

51. Cox R, Schapiro AC, Manoach DS, Stickgold R. Individual Differences in Frequency and Topography of Slow and Fast Sleep Spindles. Front Hum Neurosci. 2017;11:433. doi:10.3389/fnhum.2017.00433

52. Dehghani N, Cash SS, Halgren E. Topographical frequency dynamics within EEG and MEG sleep spindles. Clinical Neurophysiology. 2011;122(2):229–235. doi:10.1016/j.clinph.2010.06.018

53. Motamedi-Fakhr S, Moshrefi-Torbati M, Hill M, Hill CM, White PR. Signal processing techniques applied to human sleep EEG signals—A review. Biomedical Signal Processing and Control. 2014;10:21–33. doi:10.1016/j.bspc.2013.12.003

54. Aboalayon K, Faezipour M, Almuhammadi W, Moslehpour S. Sleep stage classification using EEG signal analysis: a comprehensive survey and new investigation. Entropy. 2016;18(9):272. doi:10.3390/e18090272

55. Armitage R. The distribution of EEG frequencies in REM and NREM sleep stages in healthy young adults. Sleep. 1995;18(5):334–341. doi:10.1093/sleep/18.5.334

56. Boostani R, Karimzadeh F, Nami M. A comparative review on sleep stage classification methods in patients and healthy individuals. Comput Methods Programs Biomed. 2017;140:77–91. doi:10.1016/j.cmpb.2016.12.004

57. Achermann P, Dijk DJ, Brunner DP, Borbély AA. A model of human sleep homeostasis based on EEG slow-wave activity: quantitative comparison of data and simulations. Brain Res Bull. 1993;31(1-2):97–113. doi:10.1016/0361-9230(93)90016-5

58. Achermann P, Borbély AA. Low-frequency (< 1 Hz) oscillations in the human sleep electroencephalogram. Neuroscience. 1997;81(1):213–222. doi:10.1016/s0306-4522(97)00186-3

59. Finelli LA, Baumann H, Borbély AA, Achermann P. Dual electroencephalogram markers of human sleep homeostasis: correlation between theta activity in waking and slow-wave activity in sleep. Neuroscience. 2000;101(3):523–529. doi:10.1016/S0306-4522(00)00409-7

60. Kim HJ, Chen S, Eden UT, Prerau MJ. A quantitative representation of continuous brain state during sleep. In: 2021 10th International IEEE/EMBS Conference on Neural Engineering (NER). IEEE; 2021:103–106. doi:10.1109/NER49283.2021.9441276

61. Riedner BA, Vyazovskiy VV, Huber R, et al. Sleep homeostasis and cortical synchronization: III. A high-density EEG study of sleep slow waves in humans. Sleep. 2007;30(12):1643–1657. doi:10.1093/sleep/30.12.1643

62. Helfrich RF, Mander BA, Jagust WJ, Knight RT, Walker MP. Old Brains Come Uncoupled in Sleep: Slow Wave-Spindle Synchrony, Brain Atrophy, and Forgetting. Neuron. 2018;97(1):221–230.e4. doi:10.1016/j.neuron.2017.11.020

63. Mölle M, Marshall L, Gais S, Born J. Grouping of spindle activity during slow oscillations in human non-rapid eye movement sleep. J Neurosci. 2002;22(24):10941–10947. doi:10.1523/JNEUROSCI.22-24-10941.2002

64. Staresina BP, Bergmann TO, Bonnefond M, et al. Hierarchical nesting of slow oscillations, spindles and ripples in the human hippocampus during sleep. Nat Neurosci. 2015;18(11):1679–1686. doi:10.1038/nn.4119

65. Steriade M, Nuñez A, Amzica F. A novel slow (<1 hz) oscillation of neocortical neurons in vivo: depolarizing and hyperpolarizing components. J Neurosci. 1993;13(8):3252–3265. doi:10.1523/JNEUROSCI.13-08-03252.1993

66. Winer JR, Mander BA, Helfrich RF, et al. Sleep as a potential biomarker of tau and β-amyloid burden in the human brain. J Neurosci. 2019;39(32):6315–6324. doi:10.1523/JNEUROSCI.0503-19.2019

67. Younes M, Gerardy B, Pack AI, Kuna ST, Castro-Diehl C, Redline S. Sleep architecture based on sleep depth and propensity: patterns in different demographics and sleep disorders and association with health outcomes. Sleep. 2022;45(6):zsac059. doi:10.1093/sleep/zsac059

68. Purcell SM, Manoach DS, Demanuele C, et al. Characterizing sleep spindles in 11,630 individuals from the National Sleep Research Resource. Nat Commun. 2017;8:15930. doi:10.1038/ncomms15930

69. Chen C, Wang K, Belkacem AN, et al. A comparative analysis of sleep spindle characteristics of sleep-disordered patients and normal subjects. Front Neurosci. 2023;17:1110320. doi:10.3389/fnins.2023.1110320

70. Crowley K, Trinder J, Kim Y, Carrington M, Colrain IM. The effects of normal aging on sleep spindle and K-complex production. Clin Neurophysiol. 2002;113(10):1615–1622. doi:10.1016/S1388-2457(02)00237-7

71. McClain IJ, Lustenberger C, Achermann P, Lassonde JM, Kurth S, LeBourgeois MK. Developmental changes in sleep spindle characteristics and sigma power across early childhood. Neural Plast. 2016;2016(1):3670951. doi:10.1155/2016/3670951

72. Borbély AA, Achermann P. Sleep homeostasis and models of sleep regulation. J Biol Rhythms. 1999;14(6):557–568. doi:10.1177/074873099129000894

73. Acharya UR, Faust O, Kannathal N, Chua T, Laxminarayan S. Non-linear analysis of EEG signals at various sleep stages. Comput Methods Programs Biomed. 2005;80(1):37–45. doi:10.1016/j.cmpb.2005.06.011

74. Coon W, Ogg M. Sleep EEG foundation models reveal within-stage microstructure that improves health screening beyond traditional stages. *npj Digital Med*. Published online July 9, 2026. doi:10.1038/s41746-026-02970-2

75. Zhu G, Li Y, Wen P. Analysis and classification of sleep stages based on difference visibility graphs from a single-channel EEG signal. IEEE J Biomed Health Inform. 2014;18(6):1813–1821. doi:10.1109/JBHI.2014.2303991

76. Adamantidis AR, Gutierrez Herrera C, Gent TC. Oscillating circuitries in the sleeping brain. Nat Rev Neurosci. 2019;20(12):746–762. doi:10.1038/s41583-019-0223-4

77. Buzsáki G. Rhythms of the Brain. Oxford University Press; 2006. doi:10.1093/acprof:oso/9780195301069.001.0001

78. Fogel S, Martin N, Lafortune M, et al. NREM sleep oscillations and brain plasticity in aging. Front Neurol. 2012;3:176. doi:10.3389/fneur.2012.00176

79. Girardeau G, Lopes-Dos-Santos V. Brain neural patterns and the memory function of sleep. Science. 2021;374(6567):560-564. doi:10.1126/science.abi8370

80. Niethard N, Ngo HVV, Ehrlich I, Born J. Cortical circuit activity underlying sleep slow oscillations and spindles. Proc Natl Acad Sci. 2018;115(39):E9220–E9229. doi:10.1073/pnas.1805517115

81. Steriade M, McCormick DA, Sejnowski TJ. Thalamocortical oscillations in the sleeping and aroused brain. Science. 1993;262(5134):679–685. doi:10.1126/science.8235588

82. Goodman LA. Kolmogorov-Smirnov tests for psychological research. Psychol Bull. 1954;51(2):160. doi:10.1037/h0060275

83. Benjamini Y, Yekutieli D. The control of the false discovery rate in multiple testing under dependency. Ann Stat. 2001;29(4):1165–1188. doi:10.1214/aos/1013699998

84. Fujisawa S, Amarasingham A, Harrison MT, Buzsáki G. Behavior-dependent short-term assembly dynamics in the medial prefrontal cortex. Nat Neurosci. 2008;11(7):823–833. doi:10.1038/nn.2134

85. Dean DA, Goldberger AL, Mueller R, et al. Scaling up scientific discovery in sleep medicine: the National Sleep Research Resource. Sleep. 2016;39(5):1151–1164. doi:10.5665/sleep.5774

86. Djonlagic I, Mariani S, Fitzpatrick AL, et al. Macro and micro sleep architecture and cognitive performance in older adults. Nat Hum Behav. 2021;5(1):123–145. doi:10.1038/s41562-020-00964-y

87. Kim J, Gulati T, Ganguly K. Competing roles of slow oscillations and delta waves in memory consolidation versus forgetting. Cell. 2019;179(2):514–526. doi:10.1016/j.cell.2019.08.040

88. Campos-Beltrán D, Zhang S, Marshall L. Differences in sleep spindles and polysomnography in humans: a meta-analysis on the influence of age, sex, and cognitive ability. Front Sleep. 2026;5:1802882. doi:10.3389/frsle.2026.1802882

89. Ujma PP, Konrad BN, Genzel L, et al. Sleep spindles and intelligence: evidence for a sexual dimorphism. J Neurosci. 2014;34(49):16358–16368. doi:10.1523/JNEUROSCI.1857-14.2014

90. Huupponen E, Himanen SL, Värri A, Hasan J, Lehtokangas M, Saarinen J. A study on gender and age differences in sleep spindles. Neuropsychobiology. 2002;45(2):99–105. doi:10.1159/000048684

91. Li Q, Wen S, Sun H, et al. The human sleep project: a multi-center clinical polysomnography dataset across the human lifespan. Sleep. Published online 2026:zsag215. doi:10.1093/sleep/zsag215

92. Silva GE, An MW, Goodwin JL, et al. Longitudinal evaluation of sleep-disordered breathing and sleep symptoms with change in quality of life: the Sleep Heart Health Study (SHHS). Sleep. 2009;32(8):1049–1057. doi:10.1093/sleep/32.8.1049

93. Ganglberger W, Sun H, Turley N, et al. Brain health from sleep EEG: a multicohort, deep learning biomarker for cognition, disease, and mortality. NEJM AI. 2026;3(3):AIoa2500487. doi:10.1056/AIoa2500487

94. Thapa R, Kjaer MR, He B, et al. A multimodal sleep foundation model for disease prediction. Nat Med. Published online 2026:1–11. doi:10.1038/s41591-025-04133-4

95. Cay G, Ravichandran V, Sadhu S, et al. Recent advancement in sleep technologies: a literature review on clinical standards, sensors, apps, and AI methods. IEEE access : pract innov open solut. 2022;10:104737–104756. doi:10.1109/ACCESS.2022.3210518

96. Goldstein CA, Berry RB, Kent DT, et al. Artificial intelligence in sleep medicine: an American Academy of Sleep Medicine position statement. J Clin Sleep Med. 2020;16(4):605–607. doi:10.5664/jcsm.8288

97. Satapathy SK, Brahma B, Panda B, Barsocchi P, Bhoi AK. Machine learning-empowered sleep staging classification using multi-modality signals. BMC Med Inf Decis Making. 2024;24(1):119. doi:10.1186/s12911-024-02522-2

98. Huang W, Wang Y, Cheng H, et al. A unified time-frequency foundation model for sleep decoding. Nat Commun. Published online 2026. doi:10.1038/s41467-025-67970-4

99. He M, Das P, Hotan G, Purdon PL. Switching state-space modeling of neural signal dynamics. PLOS Comput Biol. 2023;19(8):e1011395. doi:10.1371/journal.pcbi.1011395

