## Supplementary material for "The DYNAM-O Toolbox: Characterizing Individualized Neural Signatures in Sleep EEG"

### Appendix 1: Function calls in `runDYNAMO()`

```
runDYNAMO()
    ↓ [Part 1: Identifying TF-peaks from Spectrograms]
      ├─ computeTFPeaks()
            ↓ [First-pass: TF-peak detection optimizing for temporal resolution]
                ├─ computeSpectrogram()  [1 s window, 4 Hz spectral resolution]
                ├─ computeBaseline()
                ├─ runSegmentedData()
                │     ├─ removeBaseline()
                │     ├─ segmentData()
                │     └─ for each segment (parfor):
                │           └─ extractTFPeaks()
                │                 ├─ runWatershed()
                │                 ├─ mergeWshedSegment()
                │                 ├─ trimWshedRegions()
                │                 └─ computePeakStatsTable()
                └─ filterStatsTable()
            ↓ [Second-pass: TF-peak detection optimizing for spectral resolution]
                ├─ computeSpectrogram()  [2 s window, 2 Hz spectral resolution]
                ├─ computeBaseline()
                ├─ maskSpectrogram()
                ├─ runSegmentedData()
                ├─ filterStatsTable()
                └─ refinePeakFrequency()
    ↓ [Part 2: Computing Feature Properties of TF-peaks]
      ├─ computePeakStage()
      ├─ computePeakSOpower()
      └─ computePeakSOphase()
    ↓ [Part 3: Characterizing TF-peak Dynamics with Feature Histograms]
      ├─ SOpowerphaseHistogram()
      │     ├─ computeSOpower()
      │     ├─ computeSOphase()
      │     ├─ SOpowerHistogram()
      │     └─ SOphaseHistogram()
      ├─ fitParamBasis()
      └─ fitSplineBasis()
```

### Appendix 2: Comparison between Multi-taper Method and Wavelet Transform

In DYNAM-O, we currently employ the multi-taper method (MTM) with Fourier transform to obtain spectrograms [1,2]. There are different time-frequency transforms, such as the wavelet transform, to generate spectrograms. Both MTM for spectral estimation and wavelet transform provide valid time-frequency transforms to investigate the temporal and spectral information. Mathematically, the two spectral analysis approaches convey the same information about sleep EEG data [3], but the MTM spectrograms are more conducive to our subsequent imaging processing steps for identifying transient oscillation events based on two-dimensional (2D) peaks in the time-frequency domain.

Wavelet transform has been heavily used in sleep research for spindle analysis [4]. As frequency increases, wavelet transform offers finer temporal resolution and coarser spectral resolution [5]. In comparison, multitaper spectrogram allows for separate, constant controls of spectral and temporal resolutions through the time-half-bandwidth product parameter [2], and it offers equally spaced, discretized spectrograms on which imaging processing techniques such as the watershed algorithm can be readily applied [6]. Figure A1 shows a direct comparison of the same 30-s segment visualized on MTM spectrogram and wavelet scalogram. It is clear that both approaches can highlight strong transient oscillations (sleep spindles in this case), with peaks on the wavelet scalogram aligning with TF-peak centers on the MTM spectrogram. The DYNAM-O Toolbox is currently optimized for MTM spectrograms, but it is also compatible with using wavelet scalograms in place of MTM spectrograms to detect TF-peaks after slight modifications of parameters.

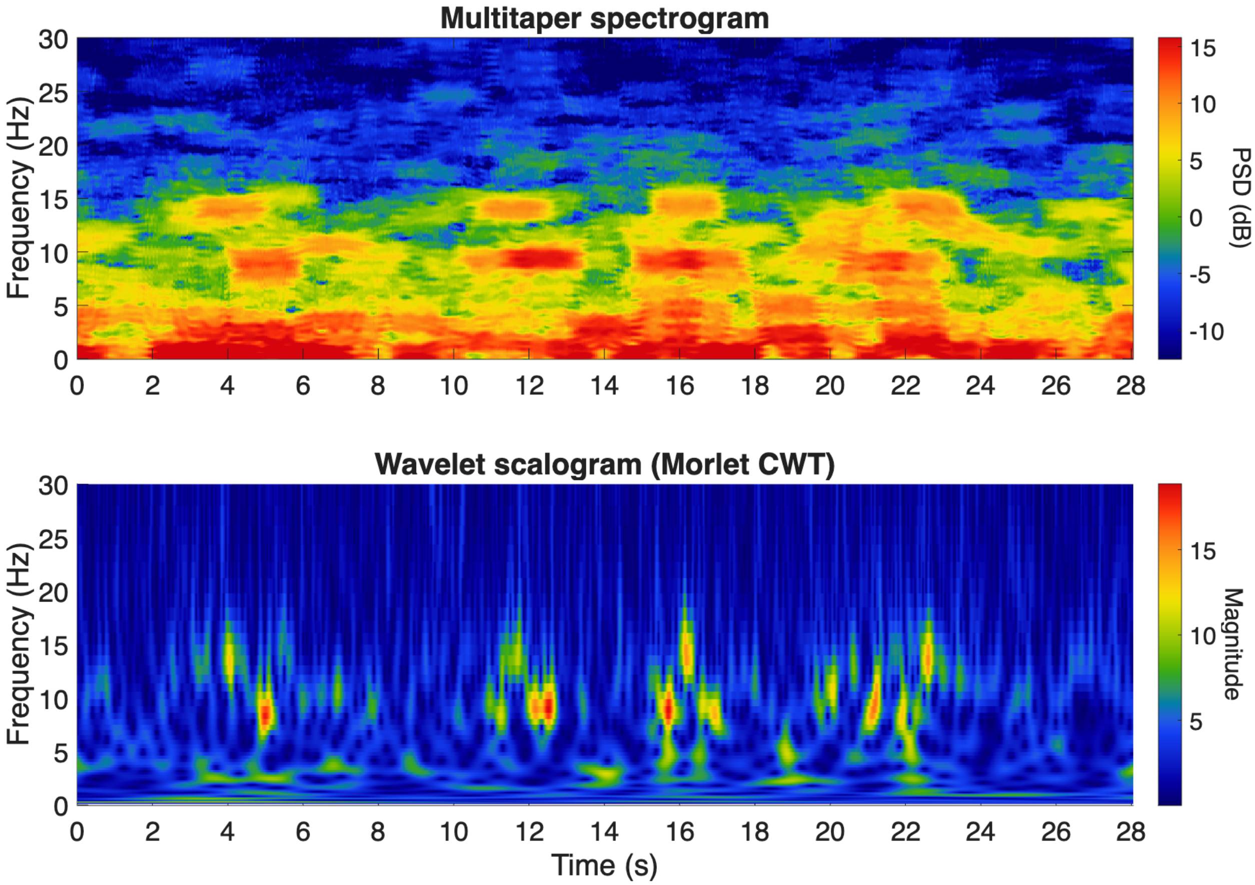


**Figure A1. Comparison of the multitaper spectrogram and the wavelet scalogram.** The same 30-s segment of sleep EEG containing sleep spindles, shown as a multitaper method (MTM) spectrogram **(top)** and a Morlet continuous wavelet transform scalogram **(bottom)**. Both transform plots highlight the same strong transient oscillations, with peaks on the wavelet scalogram aligning with TF-peak centers on the MTM spectrogram. Unlike the wavelet scalogram, whose temporal and spectral resolutions covary with frequency, the MTM spectrogram provides separate, constant control of both resolutions on an equally spaced time-frequency grid.

### Appendix 3: Background baseline normalization of spectrograms

Electrophysiological recordings are known to exhibit a 1/f-like drop-off in power with frequency [7,8]. As recently resurfaced as a methodological consideration for neuroscience studies using electrophysiology [9], it is important to distinguish between aperiodic and oscillatory signals when investigating neural activity in sleep EEG data [10]. Since the focus of our analytic approach is on transient neural oscillations during sleep, we consider the stationary and aperiodic signals to form the background baseline, to which oscillatory activity is added on the log-linear spectrum. Therefore, to emphasize transient increases in spectral power from oscillations, we normalize each spectrogram by removing the baseline spectrum [6,11].

By doing so, spectral peaks reflecting transient neural oscillations will be accentuated from the surrounding activity as 3D topographical peaks with more symmetric falling slopes on both high and low frequency ends. This allows the watershed algorithm to better identify the geometric boundaries of TF-peaks and extract peak centers. Figure A2 shows the background baseline spectrum estimated using the 2$^{nd}$ percentile of spectral power at each frequency across a night, which is largely flat and reflecting the 1/f-like spectral slope with no apparent peak structure. Removing this baseline via division on a linear scale allows TF-peaks to stand out on the normalized spectrogram for subsequent processing. Note that the watershed algorithm is applied on the linear-scale spectrogram while we visualize spectrograms on a dB scale for displaying transient oscillations at higher frequencies.

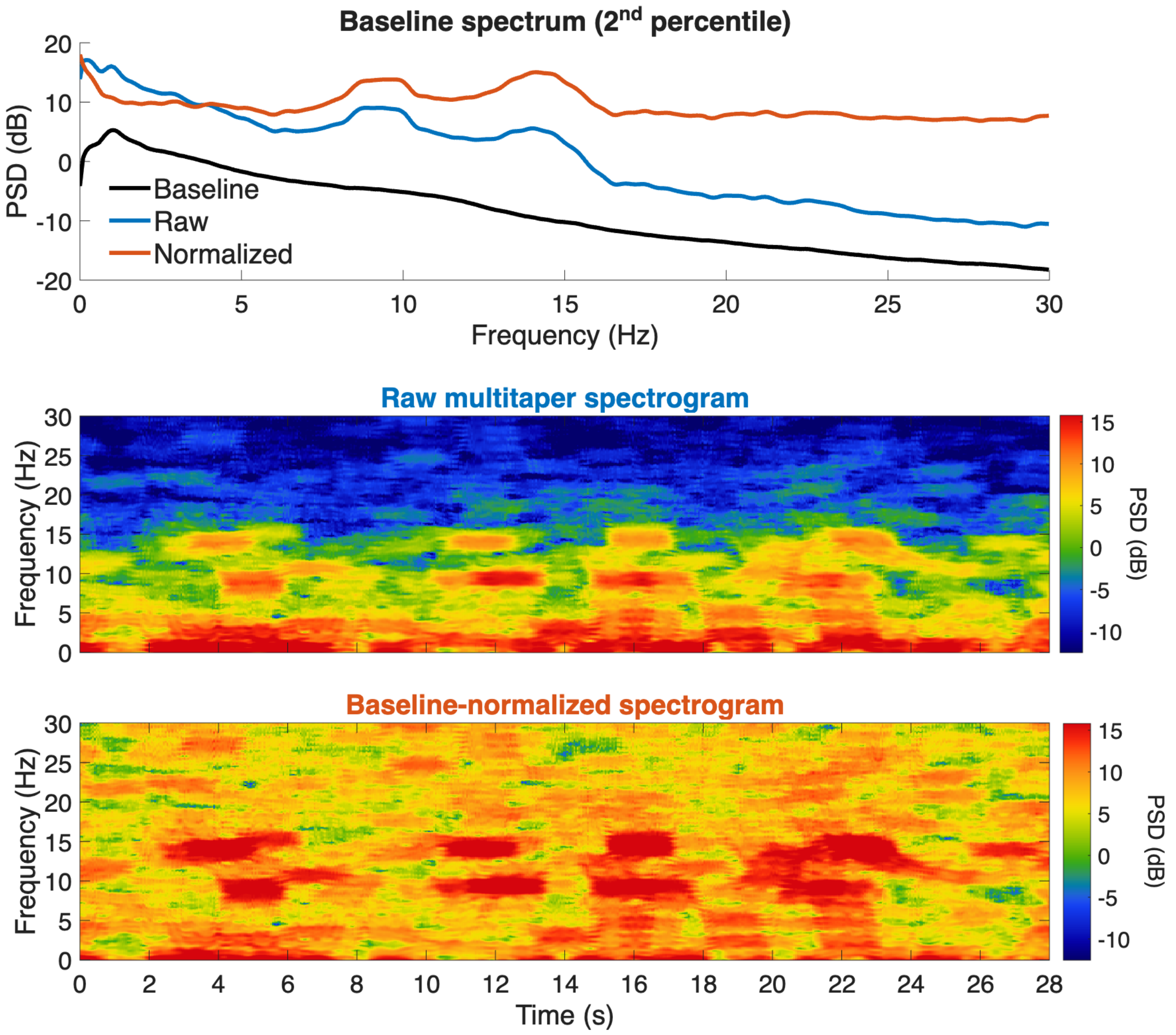


**Figure A2. Background baseline normalization of the spectrogram. (top)** Power spectra averaged across the night, shown as the raw multitaper spectrum, the background baseline estimated as the second percentile of spectral power at each frequency, and the resulting baseline-normalized spectrum; the baseline is largely flat with no apparent peak structure, reflecting the 1/f-like aperiodic spectral slope. **(middle)** The raw multitaper spectrogram of the same recording, with warmer colors indicating greater PSD. **(bottom)** The baseline-normalized spectrogram, obtained by dividing the raw spectrogram by the baseline spectrum on a linear scale.

### Appendix 4: Merging algorithm

The merging algorithm is required in DYNAM-O because applying the watershed algorithm on spectrograms results in an abundance of tiny watershed regions due to the presence of noise. This over-segmentation issue was first addressed in Stokes et al., 2023, which proposed the first version of the merging algorithm that forms the basis of the current optimized version.

The task of merging small and fragmented watershed regions can be conceptualized as a problem on a connected graph, where each watershed region forms a node. Thus, adjacent nodes should be merged when they likely represent fragments of a unified peak, while they should not be merged when they likely represent distinct peaks. This statement can be mathematically described as computing a merge weight between two regions, which consists of two parts: the first part reflects how likely the two regions (nodes) being considered are parts of the same 3D topographical peak; the second part captures how likely the two regions represent distinct peaks. The final merge weight, which corresponds to an edge weight on the connected graph, is simply the first part subtracting the second part in support of merging.

As shown in Figure A3, given two regions $i$ and $j$ , a straightforward measure of how likely the two regions are parts of the same peak can be calculated by taking the difference between the maximum of their adjoining boundary, denoted as $A_{i,j}$ and the minimum boundary points surrounding each of the two regions, $\min(B_i)$ and $\min(B_j)$. Conceptually, this measure reflects the fact that when a peak is complete, its surrounding boundaries will be at similarly low heights all around. Therefore, if a peak is broken into two fragments by watershed, the boundary line partitioning the two fragments will have an elevated height compared to the rest of the boundary of a topographical peak. Therefore, the first part of a directed merge weight indicating how critical it is to merge region $j$ into region $i$, denoted as $C_{i,j}$, and vice versa are:

$$C_{i,j} = \max(A_{i,j}) - \min(B_i)$$

$$C_{j,i} = \max(A_{i,j}) - \min(B_j)$$

The second part of the merge weight needs to indicate how likely the two regions represent distinct peaks. A straightforward measure can be obtained by taking the difference between the height of a watershed region $j$, denoted as $j_{peak}$, and the maximum of the adjoining boundary $A_{i,j}$. Conceptually, if a region's peak height is much more elevated compared to the adjoining boundary, it likely reflects a distinct peak. This is also a directed measure, denoted as $D_{i,j}$ and $D_{j,i}$, since each of the two regions $i$ and $j$ has its own peak maximum:

$$D_{i,j} = j_{peak} - \max(A_{i,j})$$

$$D_{j,i} = \mathrm{i}_{\mathrm{peak}} - \max(A_{i,j})$$

The merging action between any two regions is symmetric, regardless of which region shows the stronger evidence to be merged into the other. Therefore, we obtain the equation for an undirected edge weight, $w_{i,j}$, between two adjacent watershed regions $i$ and $j$ as:

$$E_{i,j} = C_{i,j} - D_{i,j} = 2\max(A_{i,j}) - \min(B_i) - j_{peak}$$

$$E_{j,i} = C_{j,i} - D_{j,i} = 2\max(A_{i,j}) - \min(B_j) - i_{peak}$$

$$w_{i,j} = \max(E_{i,j}, E_{j,i})$$

Given a set of watershed regions, we form an initial connected graph by computing the edge weights between all adjacent regions following the above equations. With this initial graph, we then perform merging in an iterative manner until a stopping criterion is reached: the two regions with the largest $w_{i,j}$, between them on the current graph are first merged together, and impacted edge weights with their neighbors are recomputed. This process repeats until the largest recomputed edge weight falls below a threshold. We employ a fixed threshold at 11. Alternatively, a dynamic threshold can be placed at the 95$^{th}$ percentile of all edge weights on the initial graph in order to accommodate different levels of noise in different sleep EEG datasets. Empirically, we have verified that this merging algorithm performs with satisfactory quality across multiple datasets on the NSRR database [12], and therefore it can be assumed to be functioning properly in new analyses using the current version of the DYNAM-O Toolbox.

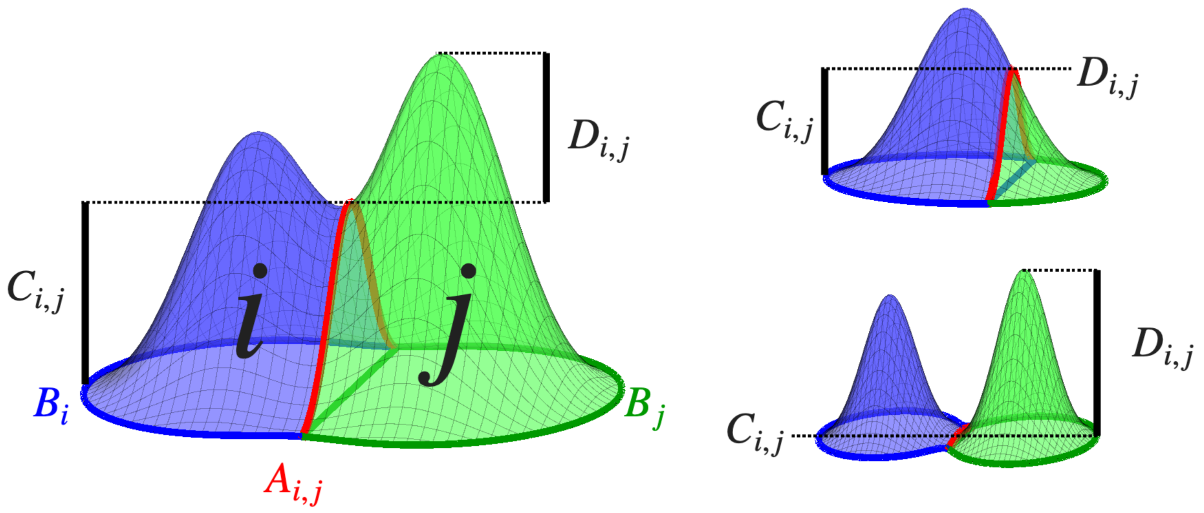


**Figure A3. The directed edge weight ($E_{i,j}$) for merging two regions is informed by two parts.** The first indicates how likely the two regions belong to the same topographical peak ($C_{i,j}$), and the second reflects how likely a region represents a distinct peak with a large peak height ($D_{i,j}$).

### Appendix 5: Separation between TF-peaks and noise watershed regions

Given the iterative merging procedure with a relatively high stopping criteria at 11 (or at the 95$^{th}$ percentile), most of the remaining watershed regions are still unmerged local maxima produced by noise that get segmented by the watershed algorithm. The vast majority of these watershed regions do not represent meaningful 3D topographical peaks in the time-frequency domain, and they unlikely reflect real transient neural oscillations during sleep. To separate TF-peaks from the rest of watershed regions that merely capture noise, we apply a set of detection thresholds that are defined by the MTM parameters used in spectrogram estimation.

When an MTM spectrogram is estimated with a set of window length and spectral resolution, we define the minimum duration of a detectable TF-peak as half of the window length, and the minimum bandwidth as half of the spectral resolution. These two lower cutoffs are justified since the MTM spectrogram is estimated with temporal and spectral spreading even for a perfect sinusoidal oscillation occurring briefly in time. Therefore, real TF-peaks that are meant to reflect prominent transient oscillations should not have finer resolutions than the limits of the spectrogram estimation, motivating the application of the detection thresholds.

The TF-peak detection step in DYNAM-O processing pipeline involves sequential optimization for temporal and spectral resolutions. Therefore, two rounds of watershed region filtering are performed, with corresponding thresholds matching the MTM parameters. Specifically, the first round has duration and bandwidth cutoffs at 0.5 s (1 s window) and 2 Hz (4 Hz spectral resolution), while the second round has cutoffs at 0.5 s (2 s window; kept at the first round value to tighten spectral resolution without raising duration threshold) and 1 Hz (2 Hz spectral resolution). Liberal upper bounds are also placed at 5 s and 15 Hz to reject unrealistic transient oscillations.

Finally, the multitaper spectral estimate is approximately chi-square distributed at large samples:

$$\hat{S}(f) \sim S(f)\frac{\chi_v^2}{v}$$

The degree of freedom of the chi-square distribution ($v$) is two times the number of tapers, and therefore the two-sided 95% confidence interval can be computed to set a lower bound on the minimum height of statistically significant TF-peaks on a linear scale:

$$h_{min} = \left(\frac{F_{\chi_v^2}^{-1}(0.975)}{v}\right)^2$$

The additional power of two is to account for the baseline normalization of removing the 2$^{nd}$ percentile spectrum, which effectively increases the floor of detectable watershed regions by squaring the error in the multitaper spectral estimation (or doubling the bound on a dB scale).

### Appendix 6: Sequential optimization for temporal and spectral resolutions

Given the uncertainty principle in spectral estimation using tapered Fourier transform, there is a tradeoff between temporal and spectral resolutions. The multitaper spectrogram estimation defines this tradeoff using the time-half-bandwidth product parameter, which is fixed at a minimal 2 to utilize 3 DPSS tapers for spectral estimation. As shown in the main manuscript, the previous MTM parameter used in Stokes et al., 2023 resulted in temporal and spectral resolutions of 1 s and 4 Hz, risking blending two concurrent transient oscillations close in frequency as one TF-peak event with a large bandwidth. One could sacrifice some temporal resolution to narrow the spectral resolution using a 2 s window, resulting in 2 Hz spectral resolution. However, under this second MTM parameter setting, two transient oscillations at the same frequency occurring close in time may appear as one TF-peak event with a long duration.

We achieve sequential optimization with a double loop of the watershed-merging-trimming processing, first using MTM parameters for (1 s, 4 Hz) resolutions. Once TF-peak regions are detected, these regions are converted into binary masks applied to a second spectrogram with MTM parameters for (2 s, 2 Hz) resolutions, using the function `maskSpectrogram()`. The resultant TF-peak events after the second round of processing are outputted as the final extracted TF-peaks for subsequent analyses in the DYNAM-O processing pipeline. This technique, while simple, successfully limits the tendency of TF-peaks detected on the (2 s, 2 Hz) spectrogram to link events close in time as long-duration events, since they exist within separate masked regions during the first round of TF-peak detection. This sequential optimization completely retains the same processing logics in each round, including appropriate noise filtering criteria defined by each round's MTM parameters (Appendix 5). Figure A4 shows a schematic diagram of this sequential optimization procedure with different MTM parameters for detecting TF-peaks on spectrograms.

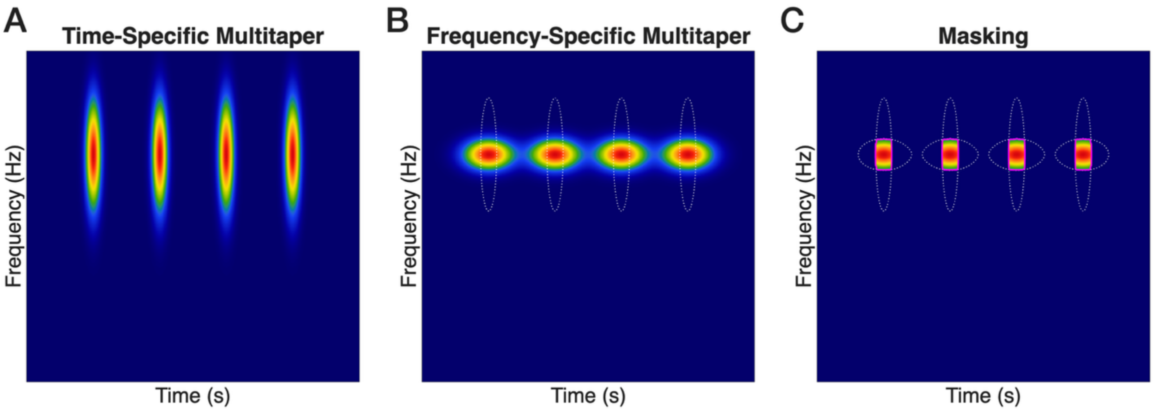


**Figure A4. Sequential optimization for temporal and spectral resolutions in TF-peak detection. (A)** The first-pass spectrogram computed with (1 s, 4 Hz) multitaper parameters resolves events that are close in time but spreads them across frequency. **(B)** The second-pass spectrogram computed with (2 s, 2 Hz) parameters resolves events that are close in frequency, but blends events close in time; dotted outlines mark the first-pass regions. **(C)** Restricting the second-pass detection to binary masks derived from the first-pass regions recovers each event as a distinct TF-peak.

### Appendix 7: TF-peak frequency refinement

Due to the inherent tradeoff between temporal and spectral resolutions during spectrogram estimation, the sequential optimization can only reach an approximate resolution of (1 s, 2 Hz) when detecting TF-peak events. However, EEG patterns of transient oscillations may have frequencies separated by less than 2 Hz. As the DYNAM-O Toolbox focuses on population distributions of TF-peaks, being able to separate clusters of TF-peaks close in frequencies requires a finer spectral resolution in the estimated peak frequency for each detected TF-peak event.

To address this need for improved frequency estimates, we perform a peak frequency refinement step after TF-peaks have been detected. For each identified TF-peak event, we extract the 4 s original recording centered at the peak time, which has been estimated from weighted centroid in the watershed-merging-trimming processing. Then we perform a tapered spectral estimation with a Hann window function, which gives approximately 1 Hz main lobe width with the 4 s analysis window, 0.05 Hz frequency resolution, and constant detrending in the window. The bounding box property is used to extract the spectrum segment corresponding to the TF-peak. A 1000-point spline interpolation is used to smooth the spectrum segment, with the frequency corresponding to the maximum on the interpolated curve identified as the refined peak frequency assigned to the TF-peak. If the maximum is identified on an edge of the bounding box, indicating a failed identification of a topographical peak within the box under a finer spectral resolution, the peak frequency is set to NaN, and the TF-peak is subsequently discarded. This frequency-refinement filtering typically removes less than 1% of the identified TF-peaks.

The choice of using the Hann window function over MTM DPSS tapers for frequency refinement requires some justifications. First, the improvement of the spectral resolution from 2 Hz to 1 Hz is entirely due to the extension of the analysis window from 2 s to 4 s. While this longer duration for spectral estimation results in smearing of the 3D spectral peak in the time-frequency domain, we are not re-detecting TF-peak events with the watershed algorithm. Instead, we only perform peak frequency refinement on already identified TF-peaks by analyzing the single spectrum obtained from the analysis window centered at each event’s peak time. Therefore, the temporal smearing is much less detrimental for the current processing objective.

Second, with a 4s analysis window, MTM spectral estimation with a time-half-bandwidth product of 2 and 3 tapers will also give a spectral resolution at 1Hz. However, the tradeoff between variance reduction and spectral leakage when averaging across multitaper tapers also hinders an accurate estimation of the event peak frequency. As can be observed on Figure A5a, the (4s, 1Hz) MTM spectrum using 3 tapers has a flat top of the main lobe peak showing a triplet structure. The two additional bumps flanking the center peak are due to spectral leakage from the last two tapers that have worse bias characteristics. While the entire triplet falls within the 1Hz bandwidth of the MTM estimate, the flat-top main lobe makes it challenging to accurately estimate the peak frequency of the event, as small perturbations due to noise or neighboring TF-peak events can result in the maximum value occurring off the "true" center of event peak frequency.

A natural solution is to only use a single DPSS taper in MTM spectral estimation, which avoids spectral leakage from the last two tapers. As shown in Figure A5a, (4 s, 1 Hz) MTM spectrum using 1 taper generates a unimodal peak centered at the true oscillation frequency (set to 14 Hz here), since the first DPSS taper maximizes the concentration of spectral energy within the +/-0.5 Hz band in the main lobe. However, a single DPSS taper no longer reduces variance, and the only gain over a Hann taper is the maximal spectral concentration within the bandwidth. As a tradeoff, the first DPSS taper still has higher side lobes compared to the Hann-windowed frequency response (Figure A5b). To accurately refine the peak frequency of an already identified TF-peak event, it is equally important to accentuate the main lobe curvature around the true peak frequency and to attenuate bias due to spectral leakage from side lobes. We use the Hann taper for this frequency refinement step, since it has better side lobe properties. Having lower side lobes means that neighboring TF-peaks will have more attenuated influence on the spectrum segment of the current event whose peak frequency is being estimated, allowing a slightly improved identification of the peak frequency. We are only able to exploit this characteristic of the Hann taper since we are refining the peak frequency by centering the analysis window at the peak time. The Hann-windowed Short Time Fourier Transform spectrogram is not suitable for the watershed-merging-trimming processing due to higher variance in the spectral estimator.

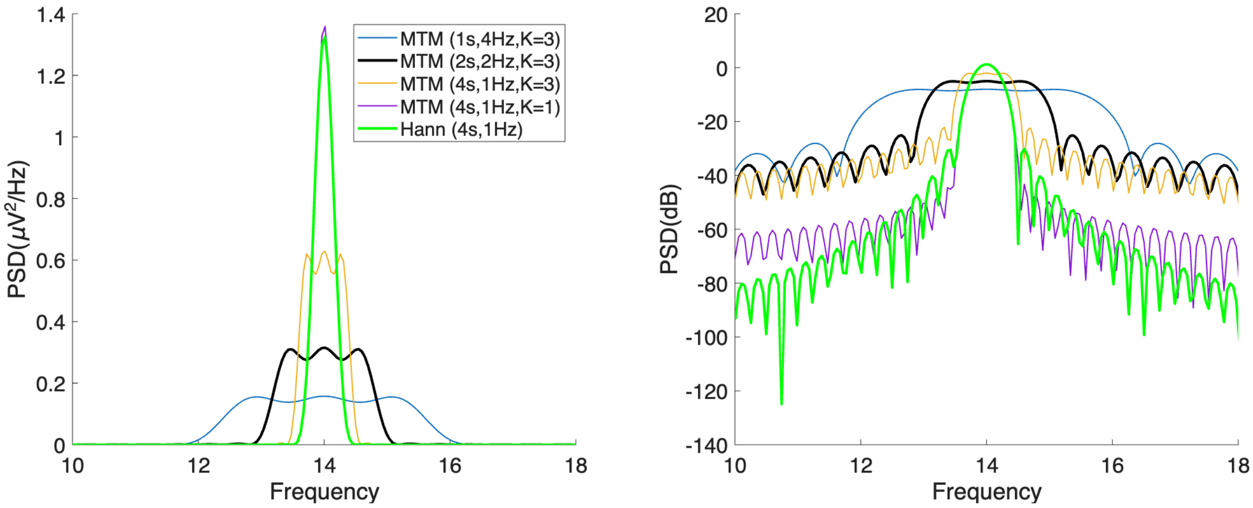


**Figure A5. Comparison of spectral estimates for peak frequency refinement.** Power spectra of a simulated 14-Hz oscillation estimated with multitaper (MTM) DPSS tapers at (1 s, 4 Hz), (2 s, 2 Hz), and (4 s, 1 Hz) resolutions using 3 tapers, with a single DPSS taper at (4 s, 1 Hz), and with a Hann taper at (4 s, 1 Hz). **(left)** Estimates on a linear PSD scale, highlighting the main-lobe shape. The (4 s, 1 Hz) three-taper MTM spectrum has a flat-topped, triplet-shaped main lobe arising from spectral leakage in the higher-order tapers, whereas the single-taper MTM and the Hann taper yield unimodal peaks centered at the true 14-Hz frequency. **(right)** The same estimates on a dB scale, highlighting that the Hann taper has the lowest side lobes, minimizing leakage from neighboring TF-peaks, while the single DPSS taper has higher side lobes despite a nearly identical main lobe.

### Appendix 8: Macroscopic sleep feature properties of TF-peaks

A key factor in understanding the overnight sleep dynamics of TF-peaks is the context in which each event occurs, specifically, whether there are patterns in the timings of discrete TF-peak events observed during a night. Many macroscopic properties can be used to describe such global brain states during sleep, and in the current version of the DYNAM-O Toolbox, we focus on three of them with well-established relationships to clinical sleep research and cortical states:

- Sleep stage at which a TF-peak event occurs
- Power of slow oscillation (0.3-1.5 Hz) as a proxy measure of depth of sleep
- Phase of slow oscillation (0.3-1.5 Hz) as a proxy measure of cortical states

The assignment of the corresponding sleep stage to each identified TF-peak is straightforward with available scoring of sleep stages: once the peak time is identified for each event, the closest previous scored sleep stage is used, which usually is available for every 30-sec epoch based on standard clinical practices for polysomnography recordings.

The total spectral power within the 0.3-1.5 Hz range of the raw recording is used to measure the strength of slow oscillations (SO). A MTM spectrogram is computed with a time-half-bandwidth product of 5, 9 DPSS tapers, a window length of 5 s and a window step-size of 0.5 s. Each 5 s window is also linearly detrended to remove the slowest frequency to reduce bleeding into the 0.3-1.5 Hz range. Once SO spectral power is calculated for the entire recording, an outlier exclusion is applied to remove SO power estimates more than 3 standard deviations away from the global mean, removing power spikes due to motion or environmental noise artifacts. The SO power is then linearly interpolated from steps of 0.5 s to the peak time of each TF-peak event. Finally, due to different gain settings across datasets and/or different impedance values at different channels, the raw SO power values vary greatly across recordings. To shift SO power to a similar scale, each recording's SO power is divided by the second percentile value on the linear scale (equivalent to subtraction on the dB scale) across a pre-specified set of sleep stages, which default to N1-3 and REM sleep, excluding wake and undefined periods. This normalization makes SO power measures from different recordings more comparable.

To compute the phase of slow oscillations, the raw recording is digitally zero-phase filtered to the 0.3-1.5 Hz frequency range with an infinite impulse response bandpass filter. The stopband frequencies are defined at 0.2 Hz and 1.6 Hz with at least 60dB attenuation in the stopband. After filtering, the analytic signal is extracted using the Hilbert transform, and the SO phase is obtained by taking the angle of the complex signal (phase of the real cosine projection). The SO phase is also linearly interpolated to the peak time of each TF-peak event.

Together, the three macroscopic properties `PeakStage`, `SOpower`, and `SOphase` are computed for each identified TF-peak event and included in the `stats_table` output.

### Appendix 9: Constructing slow oscillation-power and phase feature histograms

The slow oscillation feature histograms are computed by counting the numbers of TF-peak events falling within each bin in the two-dimensional space of slow oscillation feature by frequency. A main difference from a statistical histogram is that bins on SO feature histograms are overlapping to introduce smoothing in order to reveal prominent modes representing TF-peak dynamics. The default settings focus on TF-peaks identified during the NREM sleep stages: N1, N2, and N3. REM sleep is currently excluded from the SO feature histograms, since slow oscillations during REM sleep are weak and similar to those in awake state – superimposing distinct forms of transient oscillations prevents clear interpretations of mode structures on the SO feature histograms.

When generating a SO-power histogram, each bin has a frequency height of 1 Hz and a step size of 0.2 Hz, producing 80% frequency overlap across neighboring bins. The default normalized SO-power range included in the histogram is between -5 and 25 dB, and each bin has a SO-power width of 2.5dB with a step size of 0.25dB. Once the bounds of the histogram bins are defined, the numbers of TF-peaks falling within each bin are counted and divided by the summed duration of NREM sleep within the bin's SO power range to produce a density measure (event / minute). For example, if a total of 10 minutes during all NREM sleep are observed to produce a slow oscillation power of 15dB, and 20 TF-peaks within a frequency bin of 1 Hz are detected during these disjoint time points across the night, the bin's corresponding density value will be $20/10 = 2$ events per minute. Figure A6 compares histograms of averaged feature properties as different ways to identify structures among TF-peak events. Lastly, to prevent unstable density estimates at extreme values of SO power observed from very few time points of NREM sleep, a minimum duration of 10 minutes is imposed on columns in the SO-power histogram to mask them on the final SO-power histogram output.

When generating a SO-phase histogram, identical frequency bin height and step size as for the power histogram are used. The bin width along the phase dimension is a fifth of the full range (2π) and a step size of 0.02π (a hundredth of the range). Densities values are also computed by dividing the numbers of TF-peaks falling within each bin by the summed duration of NREM sleep for the bin. Since SO phase is a circular measure, it is very unlikely that any column with a phase range of 0.4π will have a low duration, therefore no minimum duration is imposed. Optionally, a minimum number of peaks can be required within the 1 Hz frequency bounds for each row of bins, which is turned off by default. In addition, we normalize each row on the histogram to sum to 1, producing a proportional density measure on the final SO-phase histogram output. This normalization for the circular phase is used to reveal structures of preferred phases of slow oscillations in transient oscillations after removing raw density differences at different frequencies. Without such normalization, frequency bins with more TF-peak events will dominate the SO-phase histogram, overshadowing any real phase locking among less frequent TF-peaks occurring at other frequencies.

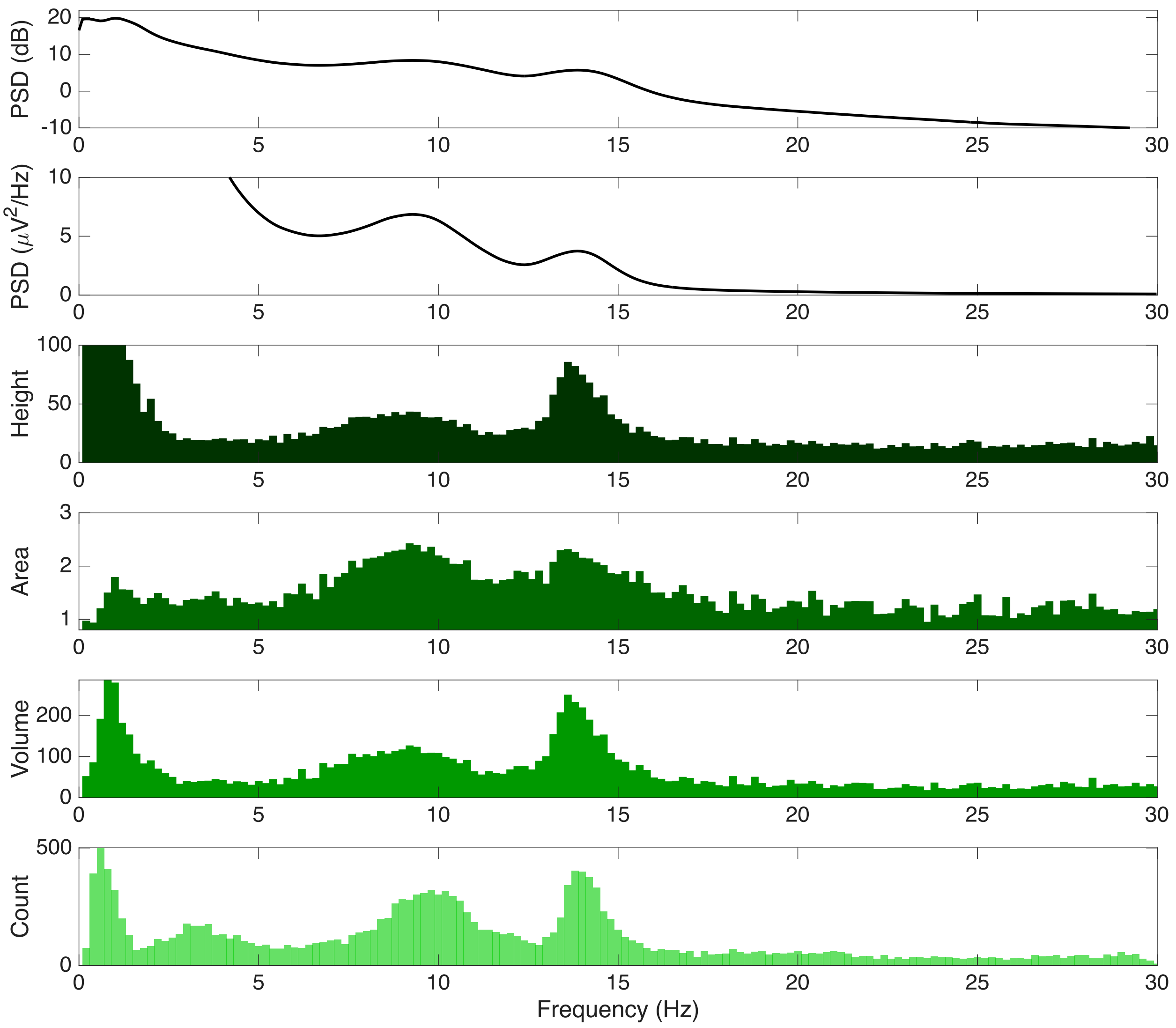


**Figure A6. Comparison of TF-peak feature histograms with averaged feature properties.** Different summaries of the same overnight TF-peaks, each plotted as a function of frequency. From top to bottom: the night-averaged power spectral density on a dB scale and on a linear scale ($\mu V^2/Hz$); followed by the TF-peak height, area, volume, and count aggregated within frequency bins. Averaging the power spectra obscures transient oscillation structure, whereas binning discrete TF-peak properties reveals prominent modes of activity in the low-frequency and sigma ranges. These comparisons motivate the use of TF-peak count histograms to characterize transient oscillation dynamics.

### Appendix 10: Dimensionality reduction with interpretable parametric modes

The objective of this dimensionality reduction approach is to construct a low-dimensional and easily interpretable representation of the SO-power and phase histograms using functions that mimic distinct mode structures. To achieve this, we formulate basis functions tailored to reflect the observed mode structures within the histograms, enabling us to express the entire histogram as a summation of these modes. Each mode is characterized by parameters that capture key structural attributes, including centroid, amplitude, width, and height, facilitating their interpretation within the context of the underlying systems. This characterization, while potentially limited in flexibility, yields a meaningful representation consisting of only a handful of parameters per mode, providing multiple orders of magnitude in dimensionality reduction.

The algorithm is summarized as follows:

1. Define a set of interpretable parametric basis functions.
2. Identify potential peaks in the histogram as initial conditions.
3. Estimate the histogram as a sum of the basis functions.

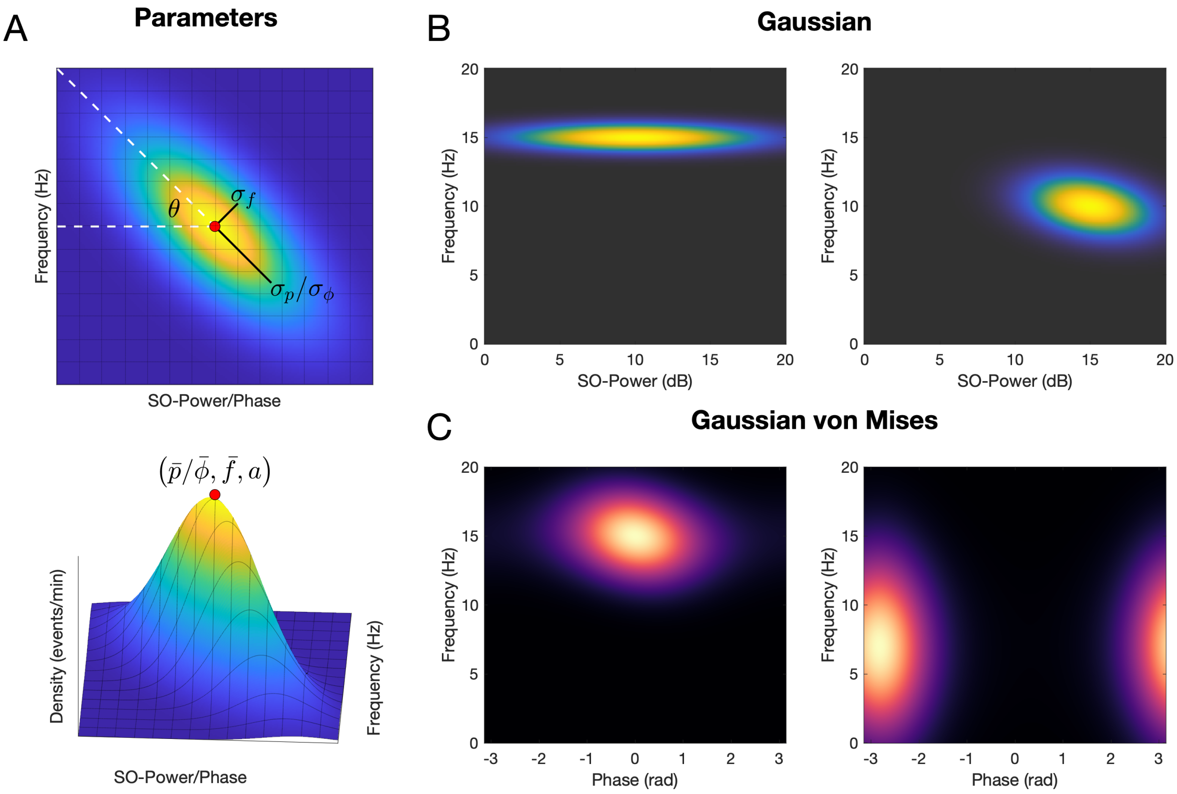


**Figure A7. Parametric mode basis functions. (A)** To estimate the structure of a mode within the SO-power and phase histograms, we use functions based on a 2D Gaussian with rotation, which provides estimates of interpretable parameters related to mode structure. This provides substantial dimensionality reduction given that there are only 6 parameters per peak. For the SO-power histograms, we use a 2D Gaussian with rotation. For the SO-phase histogram, we use a 2D hybrid von Mises-Gaussian basis function, which acts as a 2D Gaussian with rotation that is circular in the phase dimension. Examples of power and phase modes are shown in **(B)** and **(C)**, respectively.

### Model Form

To parameterize the SO-power and phase histograms, we devise basis functions to represent the structure of the density modes.

Given a series of $N$ modes $m_1, \dots m_N$, we define the parameterized histogram $H$ as:

$$H = \sum_{k=0}^{n} m_i \tag{1}$$

Each mode defines TF-peak density as a function of frequency and SO-power or SO-phase.

### Basis Functions

*SO-Power Modes*

We parameterize a density mode within the SO-power histogram as a two-dimensional Gaussian with rotation, where density $d$ at frequency $f$ and SO-power $p$ is defined as:

$$d(f,p) = a\, exp\left(-\frac{1}{2}\left(\frac{(f-\bar{f})\,cos(\theta)+(p-\bar{p})\,sin(\theta)}{\sigma_f}\right)^2 - \left(\frac{-(f-\bar{f})\,sin(\theta)+(p-\bar{p})\,cos(\theta)}{\sigma_p}\right)^2\right) \tag{2}$$

This describes a mode of amplitude $a$, centered at frequency/SO-power location $(\bar{f}, \bar{p})$, with frequency/power variances of $\sigma_f^2$ and $\sigma_p^2$, with rotation of $\theta$.

We additionally include a plane to the SO-power histogram:

$$d(f,p) = \beta_0 + \beta_1 f + \beta_2 p \tag{3}$$

to provide a baseline offset for the modes.

*SO-Phase Modes*

We parameterize a density mode within the SO-phase histogram analogously to the SO-power histogram, using a hybrid structure that is Gaussian with rotation across the frequency dimension and von Mises (circular Gaussian) across the phase dimension to allow for periodicity.

We can start by defining a basis based on the von Mises distribution as a function of phase $\phi$:

$$d(\phi) = \exp(\kappa \cos(\phi - \bar{\phi})) \tag{4}$$

where the central phase is $\bar{\phi}$ and the von Mises dispersion parameter is $\kappa = 1/\sigma_\phi^2$.

While we could include the standard pdf normalization factor as a denominator, as phase is normalized in our model, the normalization will end up being incorrect for the summation of multiple phase modes. We therefore model the maximum density of the mode in terms of a free amplitude parameter $a$, with the assumption that, given normalized data input, the parameterized output will be approximately normalized.

To do so, we can normalize the von Mises distribution such that the maximum of the function will be at 1 and then multiply by the amplitude parameter $a$. Given that the maximum will be $\exp(\kappa)$ at $\phi = \bar{\phi}$, we normalize to get:

$$d(\phi) = a \exp(\kappa \cos(\phi - \bar{\phi}) - \kappa) \quad (5)$$

We then multiply by the Gaussian in the frequency dimension to get:

$$d(f, \phi) = a \exp\left(-\frac{1}{2}\frac{(f-\bar{f})^2}{\sigma_f^2}\right) \exp(\kappa \cos(\phi - \bar{\phi}) - \kappa) \quad (6)$$

Finally, to add rotation we add the following modification to define the mode density as:

$$d(f, \phi) = a \exp\left(-\frac{1}{2}\frac{(f-\bar{f})^2}{\sigma_f^2}\right) \exp\left(\kappa \cos\left(\phi - \bar{\phi} + \left(f - \bar{f}\right)\sin(\theta)\right) - \kappa\right) \quad (7)$$

This describes a mode of amplitude $a$, centered at frequency/SO-phase location $\left(\bar{f}, \bar{\phi}\right)$, with frequency/power variances of $\sigma_f^2$ and $\sigma_\phi^2$, with rotation of $\theta$.

**Parameter Estimation**

All parameters are estimated using non-linear least squares, as implemented in the MATLAB *fit()* function. Upper and lower bound constraints for parameters can be set for each parameter during the fit process. In general, lower bounds of 0 are set for any positive parameters (e.g. standard deviation, amplitudes, etc.) and parameters such as frequency and SO-power can be constrained into the range of the computed histogram. Dataset-specific bounds can be applied as needed.

To seed the initial conditions of the model, specifically the number of modes and associated parameters, we use the same watershed algorithm and merge procedure as used in a single step of the TF-peak detection—this time applied to the SO-power and phase histograms. In this way, the algorithm can detect salient modes with minimal assumptions. The properties of the detected watershed regions (i.e. centroids, amplitude, width, and height) are used to define the initial conditions of a given mode. To fit the histogram, the detected watershed regions are first sorted from largest to smallest peak amplitude. Then the model is seeded with the largest mode and fit. Then each mode, in order of peak amplitude, is added to the model and fit one by one, saving adjusted r-squared as a metric of goodness-of-fit. If the addition of the mode produces a peak that is either too small, overlaps too much with an existing peak, or does not improve goodness-of-fit enough, the mode is not added.

To handle the periodicity in the SO-phase histograms, three identical copies of the histogram are horizontally concatenated together before the watershed procedure to complete all modes spanning – π/π. Duplicate modes across periods are then removed.

**Parametric Mode Fitting Algorithm**

Given $R$ watershed regions $\{r_1, r_2, \ldots, mr_M\}$, sorted from largest to smallest amplitude, a minimum mode amplitude, $\mathrm{A}_{\mathrm{min}}$, and a maximum mode spatial overlap, $\mathrm{O}_{\mathrm{max}}$,

Repeat for iterations $i \in \{1, \ldots, N\}$ where $N$ is equal to $R$ or some maximum number of allowable modes:

1) For iteration 1, fit a single mode model, $m_1$, with initial parameters defined by $r_1$
   a) If the fit model mode has an amplitude greater than $\mathrm{A}_{\mathrm{min}}$
      i) Save the baseline model, $m_1$ and adjusted r-squared $g_1$
   b) Otherwise
      i) Terminate fit and return empty model
2) For iteration $i$
   a) Fit a new model, $m_i$, with initial conditions from model fit $m_{i-1}$ with the addition of a new mode with initial conditions from $r_i$
   b) If the fit model modes all have amplitudes greater than $\mathrm{A}_{\mathrm{min}}$ and there are no modes with spatial overlap more than $\mathrm{O}_{\mathrm{max}}$
      i) Save the iteration model, $m_i$ and adjusted r-squared $g_i$
   c) Otherwise
      i) Revert to the previous model by setting $m_i = m_{i-1}$

After all models have been fit, the optimal model iteration is selected based on the adjusted r-squared $g_i$. This can be done either by selecting some minimum change in r-square between iterations, or by more sophisticated "knee" identification methods.

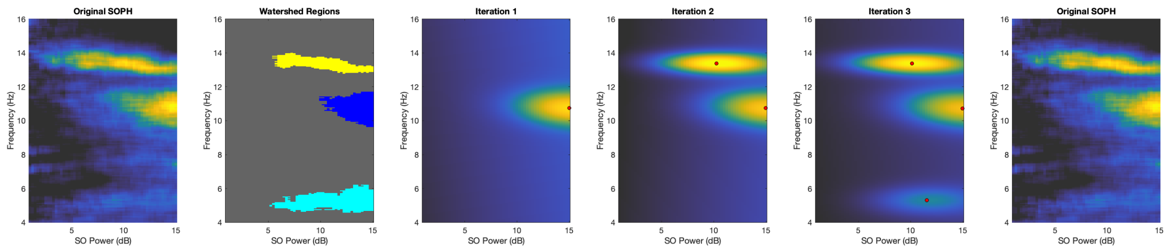


**Figure A8. Iterative fit of the SOPH using parametric basis functions to represent the modes.** The watershed regions are used to seed the initial conditions of an iterative algorithm that adds peaks to the model. Stopping criteria are evaluated as each additional peak gets added to the model.

#### Default Display of Parametric Mode Fitting Results

When invoked directly or indirectly through the `runDYNAMO()` entry point, the parametrized modes obtained from running the above fitting algorithm in the `fitParamBasis()` function are displayed in a dedicated figure as shown in **Figure A9**. This visualization shows the watershed-based seeding of parametric modes and the contours of fitted 2D Gaussian peaks. It can be seen that there are overlaps in the parameterized modes, reflecting ambiguous region partitioning in the feature histograms observed from real sleep EEG recordings.

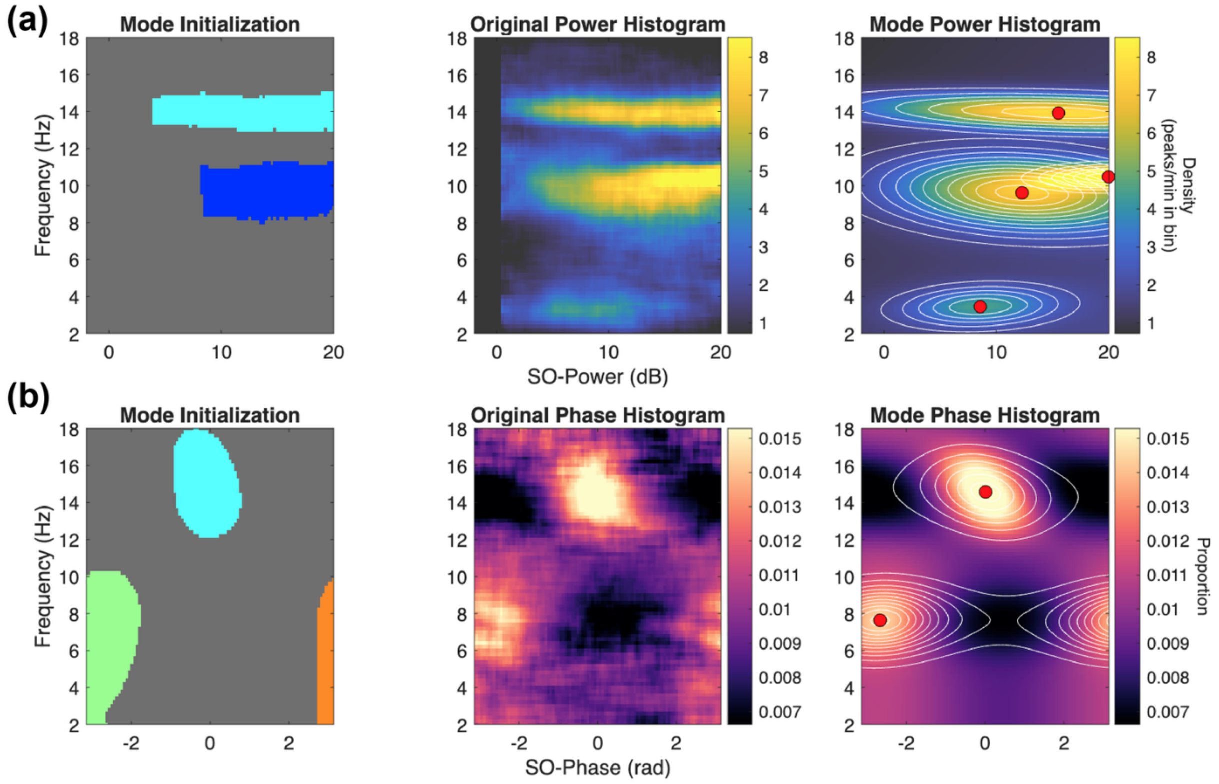


**Figure A9. Default display of parametric mode fitting to SO feature histograms in DYNAM-O. (a)** SO-power histogram, shown from left to right as the watershed-based mode initialization that seeds the fit (colored regions), the original histogram, and the fitted reconstruction overlaid with contours of the 2D Gaussian modes and their centers (red dots). **(b)** SO-phase histogram, shown in the same layout with von Mises-Gaussian basis functions.

### Appendix 11: Dimensionality reduction with 2D spline functions

Similar to the parametric modes fitted to SO feature histograms, 2D spline functions can be employed as the basis to reduce the histogram dimensionality in a nonparametric manner. Given a SO feature histogram, we perform a least-square spline interpolation to approximate the observed histogram values using the `spap2()` function in MATLAB.

To obtain a smooth low-dimensional representation of the SO-power histogram, we fit a two-dimensional tensor-product B-spline surface to the histogram values across the SO power and frequency axes. Cubic spline knot sequences are constructed along each axis using specified knot locations augmented to enforce cubic boundary conditions. The 2D spline surface is then estimated with degree-3 polynomials in both dimensions. A total of 5 knots are used for varying SO power values, and a total of 18 knots are used for the frequency dimension.

Fitting 2D spline functions to the SO-phase histogram follows the same processing, with the only exception being that 9 knots instead of 18 knots are used along the frequency dimension. This difference is due to coarser spectral variations in clusters of TF-peaks occurring at similar phases of slow oscillations. Unlike the SO-power histogram where distinct clusters of TF-peaks can be observed with small separations in frequency, SO-phase modes tend to occupy larger frequency ranges and vary slowly in their proportional density values. Therefore, the number of knots along the frequency dimension is reduced to prevent overfitting to fluctuations due to noise and misrepresenting structures of the underlying TF-peak dynamics on the SO-phase histogram.

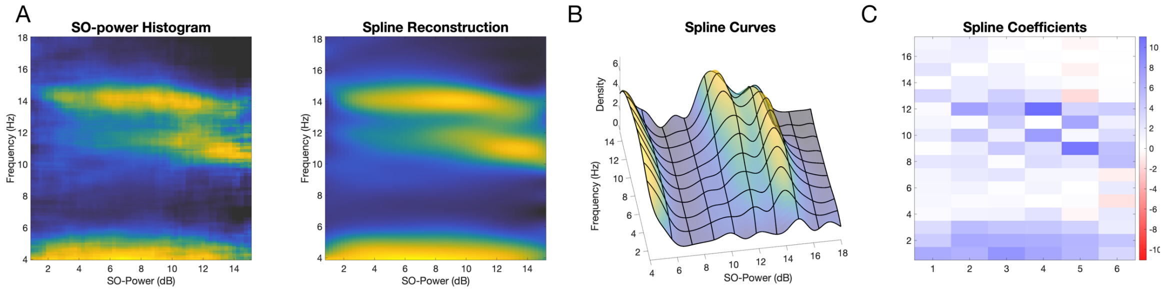


**Figure A10. Fitting with spline basis functions.** We can create a flexible surface that captures a wide variety of structures, at the expense of parameter interpretability. **(A)** Given the raw data (left), we can reconstruct the SO-power histogram (right) using spline functions. **(B)** Cardinal splines are used to fit SO feature histograms. **(C)** The resulting spline coefficients provide a low-dimensional representation of the fitted histogram.

### References

1. Percival DB, Walden AT. *Spectral Analysis for Physical Applications*. Vol 909. Cambridge; 2002. doi:10.1017/CBO9780511622762

2. Prerau MJ, Brown RE, Bianchi MT, Ellenbogen JM, Purdon PL. Sleep Neurophysiological Dynamics Through the Lens of Multitaper Spectral Analysis. *Physiology*. 2017;32(1):60-92. doi:10.1152/physiol.00062.2015

3. Bruns A. Fourier-, hilbert-and wavelet-based signal analysis: are they really different approaches? *J Neurosci Methods*. 2004;137(2):321-332. doi:10.1016/j.jneumeth.2004.03.002

4. Purcell SM, Manoach DS, Demanuele C, et al. Characterizing sleep spindles in 11,630 individuals from the National Sleep Research Resource. *Nat Commun*. 2017;8:15930. doi:10.1038/ncomms15930

5. Cohen MX. A better way to define and describe morlet wavelets for time-frequency analysis. *Neuroimage*. 2019;199:81-86. doi:10.1016/j.neuroimage.2019.05.048

6. Stokes PA, Rath P, Possidente T, et al. Transient oscillation dynamics during sleep provide a robust basis for electroencephalographic phenotyping and biomarker identification. *Sleep*. 2023;46(1):zsac223. doi:10.1093/sleep/zsac223

7. Miller KJ. Broadband spectral change: evidence for a macroscale correlate of population firing rate? *J Neurosci*. 2010;30(19):6477-6479. doi:10.1523/JNEUROSCI.6401-09.2010

8. Miller KJ, Sorensen LB, Ojemann JG, Den Nijs M. Power-law scaling in the brain surface electric potential. Sporns O, ed. *PLOS Comput Biol*. 2009;5(12):e1000609. doi:10.1371/journal.pcbi.1000609

9. Donoghue T, Haller M, Peterson EJ, et al. Parameterizing neural power spectra into periodic and aperiodic components. *Nat Neurosci*. 2020;23(12):1655-1665. doi:10.1038/s41593-020-00744-x

10. Mander BA, Dave A, Lui KK, et al. Inflammation, tau pathology, and synaptic integrity associated with sleep spindles and memory prior to β-amyloid positivity. *Sleep*. 2022;45(9):zsac135. doi:10.1093/sleep/zsac135

11. Dimitrov T, He M, Stickgold R, Prerau MJ. Sleep spindles comprise a subset of a broader class of electroencephalogram events. *Sleep*. 2021;44(9):zsab099. doi:10.1093/sleep/zsab099

12. Dean DA, Goldberger AL, Mueller R, et al. Scaling up scientific discovery in sleep medicine: the National Sleep Research Resource. *Sleep*. 2016;39(5):1151-1164. doi:10.5665/sleep.5774
